# Story about honest mistakes: The cyanobacterium *Synechocystis* has a promiscuous Entner-Doudoroff (ED) aldolase but no functional ED pathway

**DOI:** 10.64898/2026.04.01.715859

**Authors:** Ravi Shankar Ojha, Marius Theune, Ruben Fritsche, Alexander Makowka, Marko Boehm, Carmen Peraglie, Christopher Bräsen, Jacky L. Snoep, Martin Hagemann, Bettina Siebers, Kirstin Gutekunst

**Author notes:** shared first authorship (Ravi Shankar Ojha, Marius Theune).

## Abstract

In 2016, the glycolytic Entner-Doudoroff (ED) pathway was reported in cyanobacteria and plants (1). The claim was based on the biochemical characterization of its key enzyme the 2-keto-3-deoxy-6-phosphogluconate (KDPG) aldolase also named ED aldolase (EDA), on protein sequence alignments, physiological data from cyanobacterial mutants, and the *in vivo* detection of an ED pathway specific metabolite (1). However, two enzymes 6-phosphogluconate (6PG) dehydratase (EDD) and EDA are unique to this route. A recent study suggests that EDD (Slr0452) from *Synechocystis* sp. PCC 6803 most likely encodes an enzyme involved exclusively in amino acid synthesis, indicating that a complete ED pathway would be missing (2). To address the presence or absence of the ED pathway in *Synechocystis*, we conducted extended biochemical and physiological studies, revisited old data and resolved contradictions. These investigations reveal that *Synechocystis* lacks both an ED pathway and a glucose dehydrogenase/glucokinase (GDH/GK) bypass but contains a promiscuous aldolase EDA. EDA prefers KDPG as substrate but also decarboxylates oxaloacetate (OAA) and cleaves 2-keto-4-hydroxyglutarate (KHG). Synthesis of KDPG from pyruvate and glyceraldehyde 3-phosphate (GAP) is catalyzed with very low efficiency. These *in vitro* data suggest that EDA might be involved in the phosphoenolpyruvate (PEP)-pyruvate-OAA node and proline catabolism, which requires further clarification. The previous misconception was based on missing enzymatic characterizations, the oversight of a secondary mutation in a deletion strain, and an outdated view on carbohydrate fluxes. We conclude with a list of lessons and provide a solid foundation for future investigations into the role of EDA in cyanobacteria and other photoautotrophs.

**Significance statement:** This study provides a retrospective on why, for many years, it was mistakenly assumed that the glycolytic Entner-Doudoroff (ED) pathway exists in the cyanobacterium *Synechocystis* sp. PCC 6803. It shows that the first enzyme of this pathway, ED dehydratase EDD is absent, while the second enzyme, 2-keto-3-deoxy-6-phosphogluconate (KDPG) aldolase EDA, is present but is promiscuous, cleaving KDPG in addition to 2-keto-4-hydroxyglutarate (KHG) and decarboxylating oxaloacetate (OAA) *in vitro*. Finally, valuable lessons are drawn from prior misconceptions and experimental limitations. This study provides a solid foundation for future studies on the role of the ED aldolase in absence of the ED pathway in cyanobacteria and other photoautotrophs.

## Introduction

The carbon metabolism of cyanobacteria includes anabolic processes that build carbohydrates, like the Calvin-Benson-Bassham (CBB) cycle, as well as catabolic processes such as glycolytic pathways, in which these carbohydrates are consumed to provide intermediates, energy and reducing equivalents. In the day-night rhythm, carbohydrate fluxes alternate between anabolic and catabolic directions. In 2016, it was claimed that cyanobacteria and plants possess the Entner-Doudoroff (ED) pathway as a glycolytic route besides classical glycolysis (also known as Embden-Meyerhof-Parnas (EMP) pathway) and the oxidative pentose phosphate (OPP) pathway (Fig. 1A) (1). The ED pathway utilizes two enzymes that are unique to this route. The first enzyme is 6-phosphogluconate (6PG) dehydratase, abbreviated as EDD (ED dehydratase), which converts 6PG to 2-keto-3-deoxy-6-phosphogluconate (KDPG). The second enzyme is KDPG aldolase, abbreviated as EDA (ED aldolase), which splits KDPG into pyruvate and glyceraldehyde 3-phosphate (GAP). This claim was mainly based on the biochemical characterization of purified KDPG aldolases (EDAs) from the cyanobacterium *Synechocystis* sp. PCC 6803 (hereafter *Synechocystis*) and the plant *Hordeum vulgare*, in combination with protein sequence alignments, growth experiments with selected cyanobacterial deletion mutants and the detection of gluconate and KDPG in *Synechocystis* and *H. vulgare,* from which KDPG is a metabolite that is indicative of an active ED pathway (1). In the following years, *Synechocystis* mutants in which EDA was deleted either alone or in combination with other glycolytic enzymes, such as 6PG dehydrogenase (GND), which belongs to the OPP pathway, were characterized in detail and yielded contradictory results (3–7). These observations raised the question whether EDA could be either a promiscuous aldolase or an enzyme with moonlighting functions (3). Slr0452 from *Synechocystis* and its homologues in plants were considered to be EDDs (1). This consideration was based on protein sequence alignments, and in addition on a report of a promiscuous dihydroxy acid dehydratase (DHAD) with gluconate dehydratase (GAD) activity in *Sulfolobus solfataricus* also discussing similarities between the two dehydratases DHAD and EDD (1, 8). However, a recent study reported that the assumed EDDs in cyanobacteria and plants are instead in most cases exclusively DHADs (2). DHADs are specifically involved in the synthesis of the branched amino acids valine, leucine and isoleucine. Accordingly, this raises further questions about the presence of the ED pathway in cyanobacteria and plants.

**Figure 1:**
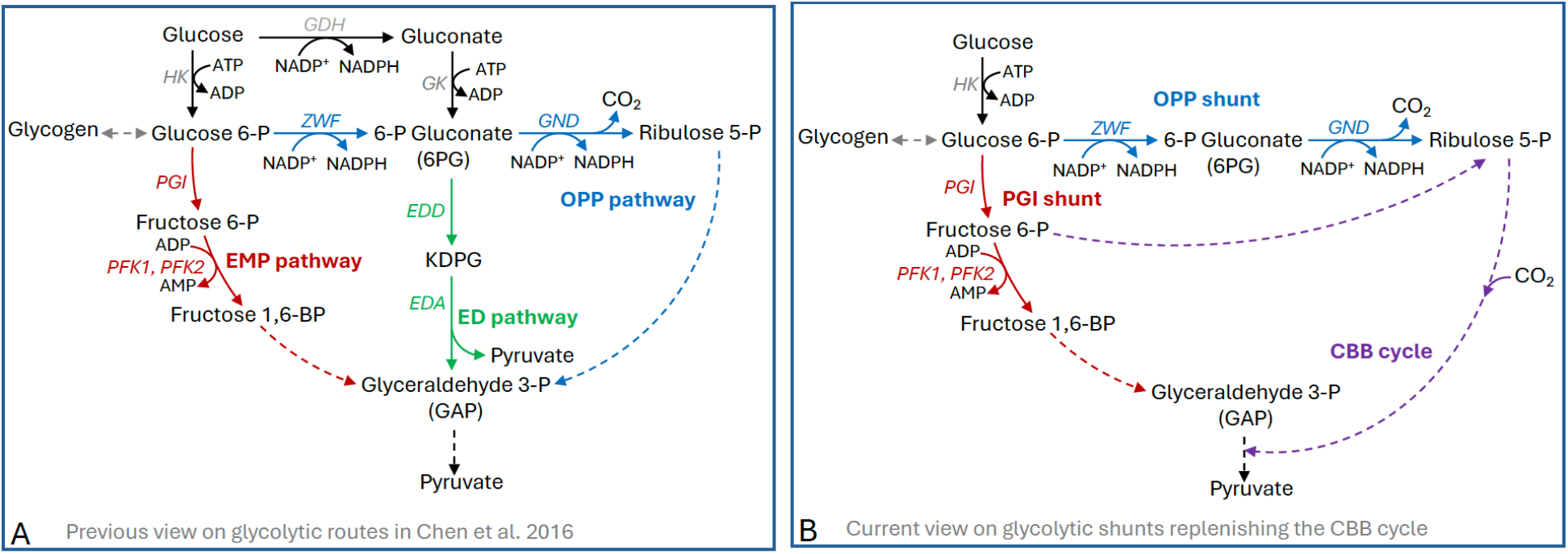
Previous and current view on glycolytic routes, glycolytic shunts and the CBB cycle in *Synechocystis*. (A) The central carbohydrate metabolism in *Synechocystis* as described by Chen et al (1). The shown GDH/GK bypass and ED pathway were assumed to be present based on the misinterpretation of experimental data (1). It was unclear at that time how glycolytic routes and the CBB cycle are organized under photomixotrophic conditions on glucose in continuous light. (B) Updated view where glycolytic routes and the CBB cycle are intertwined under photomixotrophic conditions so that glucose is fed via the glycolytic PGI and OPP shunt into the CBB cycle based on flux analyses (3). ED pathway and GDH/GK bypass are omitted based on results of this study. CBB, Calvin–Benson–Bassham; ED, Entner-Doudoroff; EDA, Entner-Doudoroff aldolase; EDD, Entner-Doudoroff dehydratase, EMP, Embden–Meyerhof–Parnas; GDH, glucose dehydrogenase; GK, gluconate kinase; GND, 6-phosphogluconate dehydrogenase; HK, hexokinase; KDPG, 2-keto-3-deoxy-6-phosphogluconate; OPP, oxidative pentose phosphate; PFK, phosphofructokinase; PGI, phosphoglucose isomerase; ZWF, glucose-6-phosphate dehydrogenase.

The anabolic CBB cycle and the catabolic glycolytic EMP and OPP pathways share several enzymes, requiring a sophisticated regulation of metabolic routes. Under photomixotrophic conditions, organic carbon (e.g., glucose) is consumed from the medium and metabolized in parallel with CO2 fixation, accelerating growth compared to photoautotrophic conditions (9). It was previously observed that growth of the *Synechocystis* mutant Δ*pfk*Δ*zwf,* in which the EMP and the OPP pathway were interrupted by deleting both phosphofructokinases (*pfk-A1* and *pfk-A2*) and glucose-6-phosphate dehydrogenase (*zwf*), was still able to enhance its growth in the presence of glucose (see Fig. 1) (1). This observation suggested the presence of an alternative route for carbohydrate degradation. When bioinformatic protein sequence analyses revealed the presence of both enzymes that are unique to the glycolytic ED pathway: EDD (Slr0452) and the KDPG aldolase EDA (Sll0107), this contradiction seemed to be resolved. EDA was purified and verified biochemically as KDPG aldolase (1). In addition, two putative glucose dehydrogenases (GDHs) (Sll1709 and Slr1608) were identified through bioinformatic studies. Since ZWF, which supplies the substrate 6PG for EDD, was deleted in Δ*pfk*Δ*zwf*, it was assumed that a putative GDH, together with a gluconate kinase (GK), would convert glucose (bypassing hexokinase (HK) and ZWF) to gluconate and further to 6PG (Fig. 1). Growth experiments with the respective deletion mutants seemed to support these assumptions (1). Furthermore, gluconate and KDPG were detected in cells of the *Synechocystis* wild type (WT) and Δ*pfk*Δ*zwf* via IC-ESI-MS/MS (1). Hence, it was concluded that the accelerated growth of the Δ*pfk*Δ*zwf* mutant was made possible by the existence of a previously overlooked glycolytic ED pathway in combination with a GDH/GK bypass (1). Notably, at that time, it was not known whether the CBB cycle and glycolytic routes are spatially separated in the cytoplasm or rather intertwined with sophisticated regulation to enable a switch between anabolic and catabolic reactions. Subsequently, flux analyses with labelled glucose under photomixotrophic conditions with WT and Δ*eda* yielded interesting, but also contradictory results, as will be outlined in detail below (3). It was observed that glycolytic routes and the CBB cycle are highly intertwined in that sense that catabolic glycolytic routes are shortened to shunts that feed carbohydrates as anaplerotic reactions into the anabolic CBB cycle and thereby accelerate CO2 fixation (3, 4). Two glycolytic shunts were identified: the OPP shunt, comprising the first two reactions of the OPP pathway catalyzed by ZWF and GND, and the PGI shunt, which includes only the first enzyme of the EMP pathway, phosphoglucose isomerase (PGI) (see Fig. 1) (3, 4). Whereas 27.6% of the labeled glucose was directed to the glycogen pool, the majority, 63.1%, was metabolized via the PGI shunt into the regenerative CBB cycle, and 9.3% were channeled via the OPP shunt to the CBB cycle (3). Contrary to our expectations, no flux via the ED pathway was detected, and no KDPG was measurable, despite the use of ultrasensitive LC–MS/MS.

The earlier assumption that the accelerated growth on glucose of the glycolytic Δ*pfk*Δ*zwf* mutant requires the GDH/GK bypass and the ED pathway, had been proven wrong by the described flux analyses, as this growth behavior can easily be explained by the existence of the PGI shunt which is responsible for the main flux of glucose into the CBB cycle and is still intact in the Δ*pfk*Δ*zwf* mutant (Fig. 1) (3). However, although there was no detectable flux via the ED pathway in the WT, carbohydrate fluxes were still altered in Δ*eda*. The flux via the PGI shunt was not modified, but a higher portion of labeled glucose was directed to the glycogen pool, and the flux through the OPP shunt was reduced (3, 4). Further studies yielded similarly conflicting results concerning the existence of the ED pathway and particularly concerning the phenotypes of Δ*eda* and the double mutant Δ*eda*Δ*gnd*. Both Δ*eda*Δ*gnd* and Δ*eda* were impaired in their ability to recover from nitrogen starvation, Δ*eda* had a reduced ability to respond to CO2 shifts, the reactivation of the dark arrested CBB cycle was delayed and growth of mutants in which either *eda* was deleted alone or in combination with other glycolytic enzymes was not understandable on the mere interruption of glycolytic fluxes (4, 5, 7). Therefore, these observations suggested that alterations in Δ*eda* and Δ*eda*Δ*gnd* might rather be based on regulatory aspects such as a moonlighting functions or substrate promiscuity of EDA instead of missing fluxes (3). Enzymes with substrate promiscuity accept a broad range of substrates because their active sites can accommodate molecules with different shapes or chemical groups. Therefore, they catalyze several reactions in addition to the primary, most efficient one (10). Moonlighting enzymes, on the other hand, perform additional functions that are independent of the catalytic activity and active site of an enzyme, such as protein-protein interactions, regulation of gene expression, or binding to RNA or DNA as transcription factors (11).

The most puzzling contradiction that remained was the observation that the intermediate 6-P gluconate (6PG) accumulated only in the double mutant Δ*eda*Δ*gnd* (where both 6PG effluxes should be interrupted in the case of an existing ED pathway) but neither in the single mutant Δ*eda* nor in Δ*gnd* (see Fig. 1A) (4).

In order to understand previous results in detail against the background of contradictory data and the new bioinformatic information that EDD is absent in most cyanobacteria (2, 12), we went back to old data, purified and tested the putative EDD (Slr0452), checked the existence of the previously presumed GDH/GK bypass, performed a series of control measurements with different mutants and resolved the accumulation pattern of 6PG in deletion strains. A detailed biochemical characterization of the KDPG aldolase EDA (Sll0107) reveals its promiscuity and provides initial evidence for its possible role in the absence of EDD in the cyanobacterium *Synechocystis*.

## Results and Discussion

### *In vitro* characterization of the predicted EDD/DHAD in *Synechocystis*

Based on the biochemical characterization of the KDPG aldolase EDA in *Synechocystis*, we had previously assumed that Slr0452 encodes for an enzyme that is involved in both amino acid synthesis as dihydroxy acid dehydratase (DHAD) and the ED pathway as EDD, as it was shown that the bifunctional DHAD (DHAD, IlvD/EDD superfamily) in the (hyper)thermophilic archaeon *Sulfolobus solfataricus*, also accepts D-gluconate - and to a lesser extent, galactonate - as alternative substrates in addition to its canonical role in branched-chain amino acid biosynthesis (8). Similarly, DHAD from *Rhizobium leguminosarum* bv. *trifolii* exhibits activity toward several C5 and C6 sugar acids, including gluconate. In addition, and the DHAD from *Fontimonas thermophila* has been engineered to increase its gluconate-dehydration activity by an order of magnitude. Together, these observations underscore the intrinsic substrate promiscuity of IlvD enzymes in the IlvD/EDD superfamily (13, 14).

However, although at least some DHADs exhibit activity toward gluconate (sugar acids), they have not been shown to accept 6-phosphogluconate (6PG), the physiological substrate of EDD in the canonical ED pathway, from any organism. One possible explanation for this limited overlap in substrate specificity was proposed by Evans et al.(2), who suggested that gluconate, being smaller and unphosphorylated, may fit within the DHAD substrate-binding pocket normally occupied by dihydroxy-isovalerate. However, the lack of crystal structures of DHAD bound to dihydroxy-isovalerate or of EDD bound to 6PG precludes direct structural comparisons, leaving the precise geometric and electrostatic determinants of substrate discrimination between these enzymes unresolved. Of note, archaea like *S. solfataricus* utilizing modified, branched ED pathway versions involving the dehydration of gluconate to KDG, do not employ a DHAD but a dedicated gluconate dehydratase (GAD) (phylogenetically grouped within the enolase superfamily, specifically a subgroup related to the mandelate racemase/muconate lactonizing enzyme (MR/MLE)) to catalyze this conversion (15, 16). This indicates that DHAD side activity alone might be insufficient to sustain physiologically relevant flux through the modified ED pathway. To date, there is no evidence that DHAD can functionally replace a bona fide GAD from the enolase superfamily in these organisms *in vivo*.

A recent study supposes that Slr0452 from *Synechocystis* which was assumed to be an EDD is an exclusive DHAD based on the biochemical characterization of its homolog in soybean (*Glycine max*) (2). Notably, organisms harboring both DHAD and EDD activity typically encode an additional IlvD/EDD superfamily gene, likely originating from gene duplication. Thus, the presence of only a single DHAD candidate in *Synechocystis* argues against an EDD function. DHAD activity was indeed shown for Slr0452 in another study, however, its potential activity toward 6PG as a substrate was not evaluated, leaving unresolved whether Slr0452 encodes a strict DHAD or a bifunctional DHAD/EDD enzyme (17). To address this question, the *slr0452* gene from *Synechocystis* was heterologously expressed in *E. coli* and recombinant protein was purified (Fig. S1). For comparison, a DHAD from the cyanobacterium *Synechococcus elongatus* was purified and assayed under identical conditions as a positive control (12) (Fig. S2). Enzymatic assays confirmed that Slr0452 catalyzes the dehydration of 2R-dihydroxyisovalerate (2R-DHIV) to 2-ketoisovalerate (KIV), consistent with DHAD function (Fig. S3A). Slr0452 exhibited a specific activity of 2.0 U mg⁻¹, whereas DHAD from *S. elongatus* showed 0.8 U mg⁻¹ with 2R-DHIV as substrate (Fig. S3A).

To test for potential dual DHAD/EDD activity, Slr0452 was subsequently assayed using 6PG as substrate. A biochemically characterized EDD from *Caulobacter crescentus* (18) served as a positive control. Despite screening a range of divalent metal cofactors (Mg²⁺, Mn²⁺, Zn²⁺, Co²⁺, Cu²⁺, and Fe²⁺), no measurable activity toward 6PG was detected for Slr0452 under any tested condition (Fig. S3B). In contrast, EDD from *C. crescentus* displayed robust 6PG dehydratase activity as expected (Fig. S2, Fig. S3B). To evaluate the possibility of a modified ED pathway in *Synechocystis*, Slr0452 and EDA (Sll0107) activities were also assayed using gluconate and KDG as substrates, respectively. *S. solfataricus* DHAD with GAD activity (*Sso*DHAD) and *Sulfolobus acidocaldarius* KD(P)G aldolase (*Saci*KD(P)GA), previously shown to catalyze reactions with gluconate and KDG, respectively (19, 20), were used as positive controls (Fig. S2, Fig. S4). Slr0452 and EDA from *Synechocystis* showed no activity on gluconate or KDG indicating that they are not involved in a modified ED pathway (Fig. S4).

Collectively, these results demonstrate that *slr0452* encodes a monofunctional, metal-dependent DHAD rather than an EDD or a bifunctional DHAD/EDD enzyme. This confirms that Slr0452 is exclusively involved in amino acid synthesis and does not catalyze the first reaction of the ED pathway as previously suggested (2, 17).

Above all, these results show that *Synechocystis* lacks a canonical 6-phosphogluconate (6PG) or gluconate dehydratase and therefore does not contain either the canonical or modified glycolytic ED pathway.

### Testing for the existence of the previously presumed GDH/GK bypass

We furthermore tested for the existence of the previously presumed glucose dehydrogenase/gluconate kinase (GDH/GK) bypass as an alternative route to hexokinase and ZWF for gluconate and 6PG production. Two putative GDHs (Sll1709, Slr1608) were previously identified in the genome based on protein sequence alignments (1). Crude cell extracts were analyzed for GDH and GK activity; however, analysis of crude extracts from *Synechocystis* WT, Δ*zwf,* as well as a putative GDH1 (Sll1709) overexpression mutant *sll1709*:oe revealed no detectable GDH activity under either photoautotrophic or photomixotrophic conditions (Fig. S5, Fig. S6, Fig. 2A-B).

**Figure 2:**
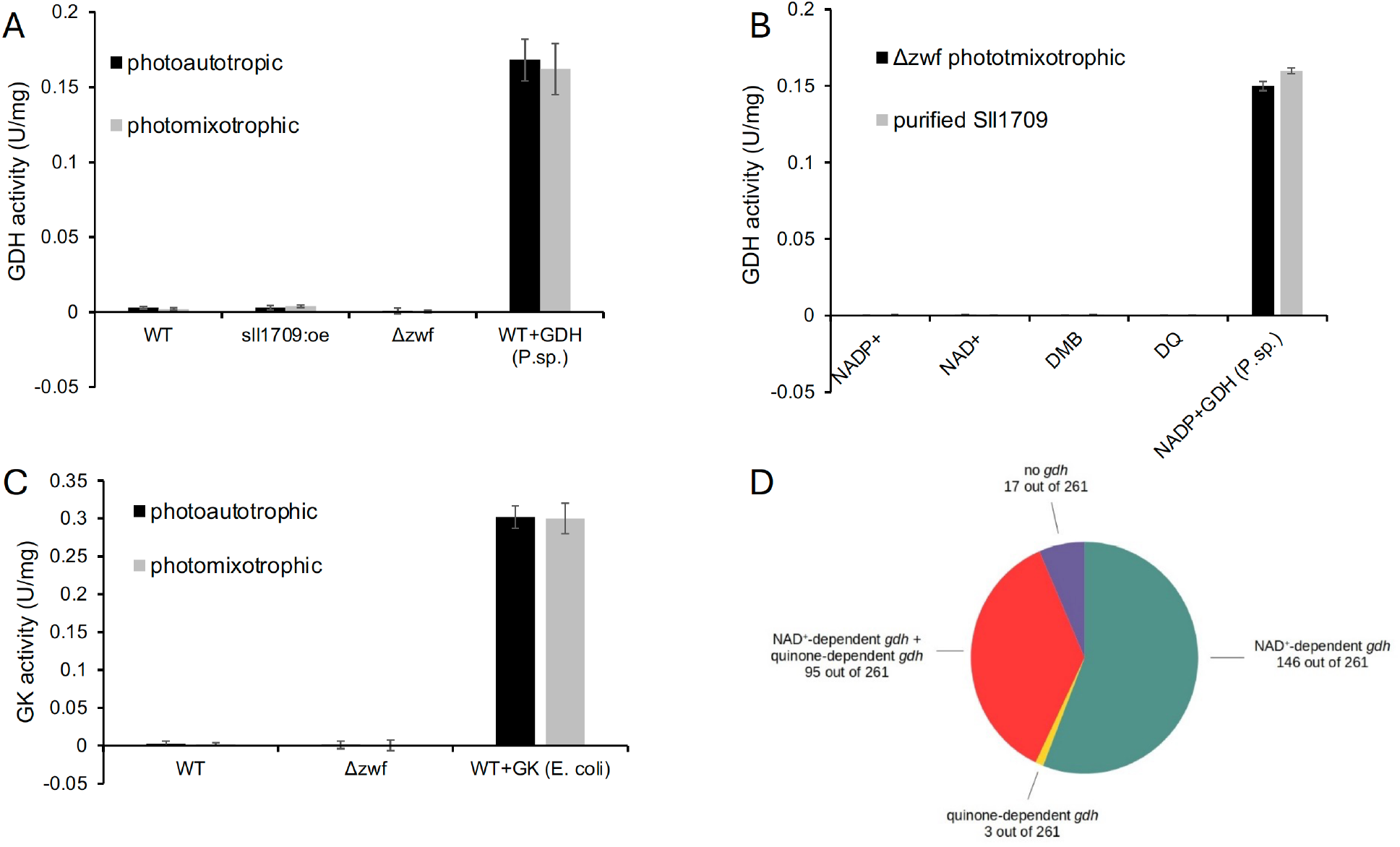
Evaluation of the proposed glucose dehydrogenase/gluconate kinase (GDH/GK) bypass in *Synechocystis* and representative cyanobacteria. (A) GDH activity measurements in crude cell extracts of *Synechocystis* WT, Δ*zwf* and a strain overexpressing putative GDH1 (*sll1709*:oe) under photoautotrophic or photomixotrophic conditions. As a positive control, 0.05 U glucose dehydrogenase from *Pseudomonas sp.* was added to WT cell extract. (B) GDH activity measurements in crude cell extracts of photomixotrophically grown Δ*zwf* cultures and with purified putative GDH1 (Sll1709). NADP^+^, NAD^+^, and the two quinones DMB (2,6-Dimethoxy-1,4-benzoquinone) and DQ (2,3-Dimethoxy-1,4-naphthoquinone) were tested as electron acceptors. As a positive control, 0.05 U glucose dehydrogenase from *Pseudomonas sp.* was added to Δ*zwf* cultures and to purified putative GDH1 (Sll1709). (C) Gluconate kinase activity measurements in cell extracts of *Synechocystis* WT and Δ*zwf* under photoautotrophic or photomixotrophic conditions. As a positive control, 0.05 U gluconate kinase from *E. coli* was added to WT cell extract. (D) Blast analyses of putative glucose dehydrogenase (GDH) in cyanobacteria. Protein databases of 261 cyanobacteria that also carry a putative gluconate kinase (GK) were analyzed for GDH homologues.

Unfortunately, we were not able to overexpress putative GDH2 (Slr1608). Crude cell extracts of Δ*zwf* were tested on the assumption that the GDH/GK route, if existent, might be upregulated in this mutant. The *sll1709*:oe overexpression mutant was tested to ensure sufficient enzyme amounts. Cell crude extracts of Δ*zwf* and purified putative GDH1 (Sll1709) were furthermore tested for GDH activity in an *in vitro* enzyme assay (Fig. S6, Fig. 2B). In this analysis, the two quinones 2,3-dimethoxy-5-methyl-1,5-benzoquinone (DMB) and duroquinone (DQ) were also tested as electron acceptors in addition to NADP^+^ and NAD^+^ since some glucose dehydrogenases are known to reduce quinones instead of NAD(P)^+^. No significant amount of GDH activity was detected for any tested electron acceptor in the GDH eluate, as well as in a cell extract of photomixotrophically grown Δ*zwf* (Fig. 2B). Taken together, these results indicate that GDH activity, which converts glucose to gluconate, is absent in *Synechocystis in vivo.* However, the inability to detect enzymatic activity is never a completely safe proof for its absence as the conditions might not have been chosen correctly, the enzyme activity might be below detection limits, or the purified enzyme might have lost its activity during purification. Furthermore, we were not able to purify and test the second putative GDH2 (Slr1608). Therefore, the second enzyme of the putative GDH/GK route was sought. The search for homologues of GK in the *Synechocystis* genome, using GKs from *E. coli* and *Pseudomonas aeruginosa* as baits, was not successful. In line with the *in silico* analyses, no gluconate kinase activity was detectable in either crude cell extracts from WT or Δ*zwf* (Fig. 2C). These results furthermore strengthen the assumption that a GDH/GK bypass is absent in *Synechocystis*.

In order to further investigate if the GDH/GK bypass is absent from all cyanobacteria, we conducted a second blast (BlastP) analysis. Similarity thresholds were set at a query score > 30 %, an identity score > 30 % and an e-value < 1^-10^. At these threshold settings we found 261 cyanobacteria that possess putative homologs of GKs from *E. coli* and *P. aeruginosa* (Table S1). We searched for homologs of GDHs in these 261 cyanobacteria next to *Synechocystis*. As baits the NAD^+^-dependent GDH 1GCO from *Bacillus megaterium* and the quinone-dependent GDH from *Acinetobacter calcoaceticus* were used (21, 22). While the results again showed a rather poor similarity for both putative GDHs (Sll1709 and Slr1608) from *Synechocystis* and thereby confirmed the lack of GDH activity in enzyme tests, good homologies for proteins were found for 244 (93 %) of the other 261 analyzed cyanobacteria (Fig. 2D, Table S1). Out of these 244 cyanobacteria that likely possess both a putative GDH and a putative GK homologue, we picked two strains, namely *Spirulina* sp. SIO3F2 and *Lyngbya aestuarii* and overexpressed and purified their putative GDHs (NEO87985.1 and WP_238987112) and the putative GK from *Lyngbya aestuarii* (WP_023068790.1) in *E. coli* (Fig. S7 to Fig. S9). Purification of the putative GK from *Spirulina* sp. SIO3F2 (NEO86388.1 MAG) was not successful. Enzyme activity assays showed a clear GDH activity for both the GDH from *Spirulina* and the GDH from *Lyngbya* (Table 1). Both NADP^+^ and NAD^+^ could function as cofactors, while higher activity was observed with NADP^+^ for both enzymes. The *Spirulina* and *Lyngbya* GDHs also used 2-Deoxy-D-glucose as substrate with 20-40% residual activity compared to glucose. Marginal activity with D-mannose was observed with the *Lyngbya* GDH (Table 1). All other tested substrates including fructose 6-phosphate, D-fucose, glucose 6-phosphate, lactose, D-glucosamine, D-lyxose, D-allose, D-arabinose, L-arabinose, D-xylose, L-xylose, and D-ribose were not used as substrates.

**Table 1:** Specific activities of the two cyanobacterial GDHs from *Spirulina* and *Lyngbya* and the gluconate kinase from *Lyngbya*. Enzyme assays were performed after recombinant expression in *E. coli* and following purification.

| Glucose dehydrogenase (GDH) | Activity (U/mg)<br>with NADP<sup>+</sup> | Activity (U/mg)<br>with NAD<sup>+</sup> |
| --- | --- | --- |
| **GDH (NEO87985.1) from**<br>***Spirulina* sp. SIO3F2** | 5.63 (Glucose)<br>2.12 (2-Deoxy-D-glucose) | 0.15 (Glucose) |
| **GDH (WP_238987112) from**<br>***Lyngbya aestuarii*** | 20.2 (Glucose)<br>4.04 (2-Deoxy-D-glucose)<br>0.26 (D-Mannose) | 2.95 (Glucose) |
| **Gluconat kinase (GK)** | **Activity (U/mg)**<br>**with ATP** |  |
| **GK (WP_023068790.1) from**<br>***Lyngbya aestuarii*** | 24.79 (10 mM Gluconate) |  |

For both the GK from *Lyngbya* and *Spirulina,* expression in *E. coli* could be observed (Fig. S9), however, purification was only achieved for the GK from *Lyngbya.* For the Lyngbya GK a specific activity of 24.8 U/mg could be determined (Table 1). In combination with already shown GDH activity, this proves the presence of the GDH/GK bypass in this cyanobacterium.

Taken together, these results indicate that there is no GDH/GK bypass in *Synechocystis,* whereas it seems to be present in other cyanobacteria, as e.g. *Lyngbya*.

Based on these results, we currently have no explanation for the gluconate that was detected in previous IC-ESI-MS/MS measurements in *Synechocystis* (1). Supplementation of the growth medium with gluconate did not enhance growth of the WT, which furthermore supports the absence of a gluconate kinase or alternatively gluconate transport into the cells (Fig. 3A). An additional control was performed by testing photomixotrophic growth of a hexokinase mutant (Δ*hk*, *sll0539*) assuming that a GDH/GK bypass if present might be upregulated in this mutant.

**Figure 3:**
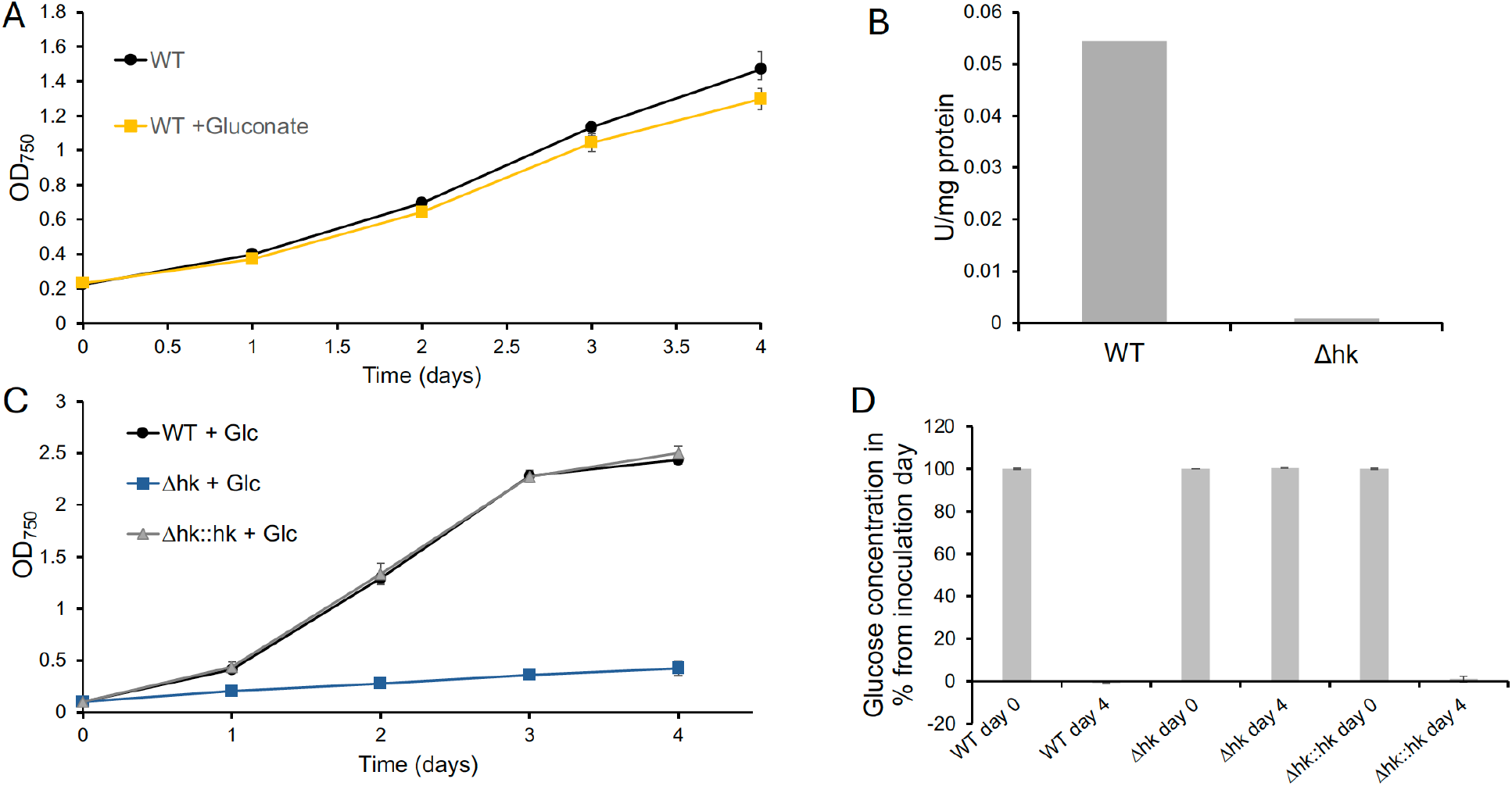
Utilization of gluconate and glucose as carbon source for growth in WT, Δ*hk* (Δ*sll0539*), and Δ*hk*::*hk*. (A) Growth of WT in the absence and presence of gluconate. (B) Hexokinase activity in WT and Δ*hk* with 10 mM glucose. (C) Photomixotrophic growth of WT, Δ*hk*, and Δ*hk*::*hk*. (D) Glucose concentration (%) in the growth medium after inoculation (day 0; 10 mM glucose) and on day 4 in WT, Δ*hk*, and Δ*hk*::*hk*.

Two enzymes (Sll0539 and Slr0329) were previously identified as putative glucose phosphorylating enzymes, with Sll0539 as the main contributor, while the deletion of Slr0329 did not affect glucose uptake or photomixotrophic growth (23, 24). Therefore, only Δ*hk* (Δ*sll0539*) was studied (25). Enzyme assays confirmed that the Δ*hk* mutant lost its ability to phosphorylate glucose (Fig. 3B). It was furthermore unable to metabolize glucose from the medium and to accelerate its growth (Fig. 3C-D). The described phenotypes could be complemented in Δ*hk::hk*. These data further support the assumption that a GDH/GK bypass is absent in *Synechocystis*, because it should allow glucose utilization in the absence of hexokinase (Hk, Sll0539) otherwise.

### Investigation into the causes why 6PG accumulates in Δ*eda*Δ*gnd* in the absence of the ED pathway, but not in Δ*eda* or Δ*gnd*

As our data indicate that the GDH/GK bypass is not present in *Synechocystis*, the only known source for 6PG would be the reaction catalyzed by ZWF (see Fig. 1). Based on this assumption, the accumulation of 6PG in Δ*eda*Δ*gnd* in contrast to Δ*eda* and Δ*gnd* is unexpected in the absence of an existing ED pathway (see Fig. 1) (4). This pattern of 6PG enrichment had led to believe for many years, despite conflicting results, that there must be at least a small flux through the ED pathway in *Synechocystis* (4). So, to resolve this apparent contradiction, we attempted to resolve the origin of the 6PG accumulation pattern in Δ*eda*Δ*gnd,* Δ*eda* and Δ*gnd*.

In order to test if ZWF is responsible for 6PG production in Δ*eda*Δ*gnd,* this enzyme was deleted in addition resulting in the mutant Δ*eda*Δ*gnd*Δ*zwf*. 6PG accumulation was abolished in Δ*eda*Δ*gnd*Δ*zwf* as expected, indicating, that ZWF activity was responsible for 6PG production in Δ*eda*Δ*gnd* (Fig. 4A).

**Figure 4:**
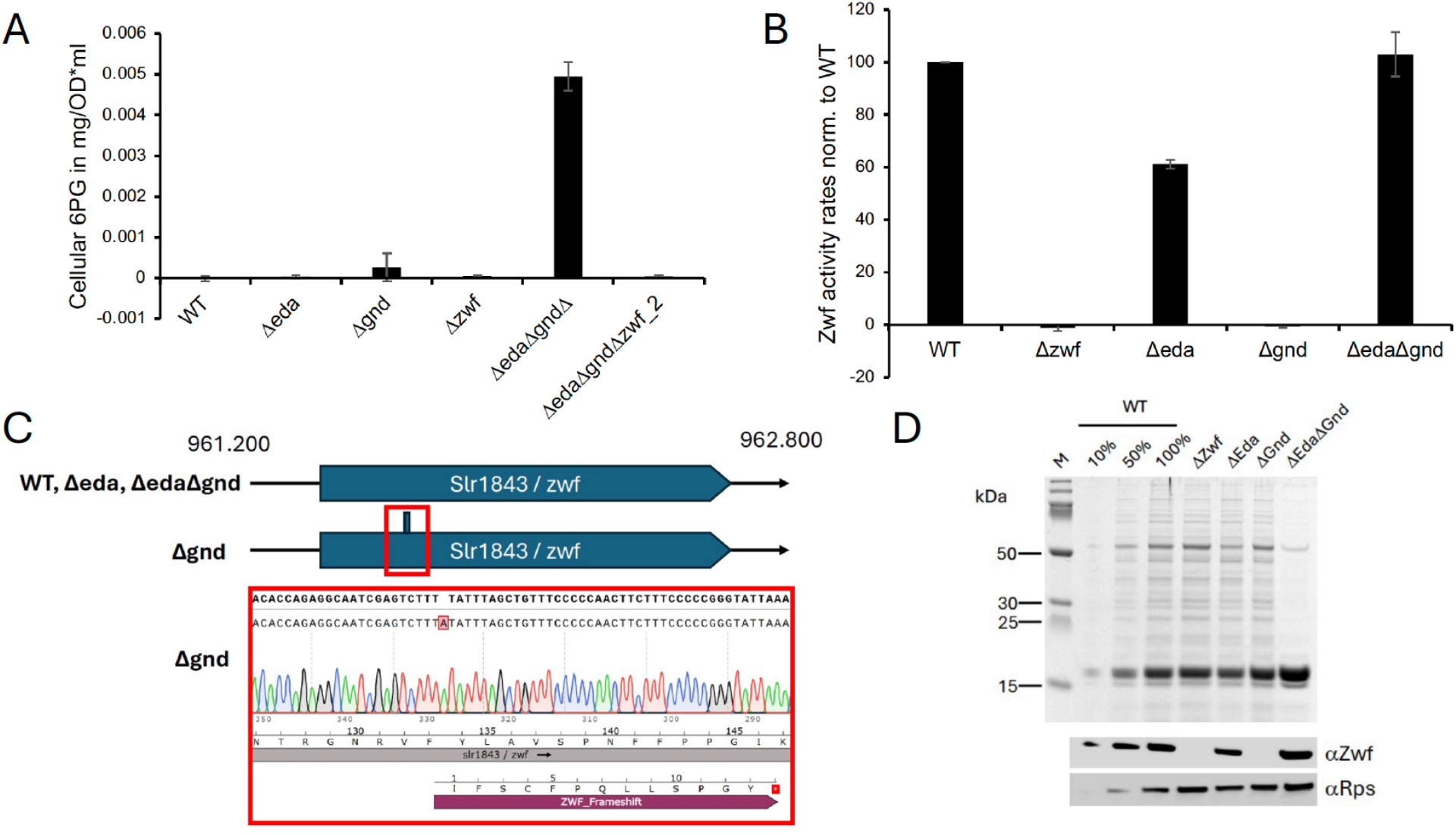
The accumulation of 6-phosphogluconate (6PG) in Δ*eda*Δ*gnd* in contrast to Δ*eda* and Δ*gnd* is not a result of a missing ED pathway flux but a secondary mutation in the *zwf* gene in Δ*gnd.* (A) 6PG accumulation in WT, Δ*eda*Δ*gnd,* and Δ*eda*Δ*gnd*Δ*zwf* under photoautotrophic conditions. (B) ZWF activity measurements in cell crude extracts of WT, Δ*zwf*, Δ*gnd*, Δ*eda*, and Δ*eda*Δ*gnd* reveals no ZWF activity in Δ*gnd.* The values are normalized to Zwf enzyme activity found in the WT based on protein content. (C) Sequencing of the *zwf* genes of WT, Δ*gnd*, Δ*eda*, and Δ*eda*Δ*gnd* revealed the insertion of an adenine in position 399 in Δ*gnd,* which results in a premature stop codon. (D) Immunoblots of soluble cell extracts of WT, Δ*zwf*, Δ*gnd*, Δ*eda*, and Δ*eda*Δ*gnd* with antibodies specific against ZWF (αZwf). An antibody against 30S ribosomal protein (αRps) was utilized as a loading control. The WT was loaded in three different concentrations (10%, 50%, 100%) to facilitate relative quantification of the αZwf signals. ZWF was neither expressed in Δ*zwf* nor in Δ*gnd*.

We subsequently tested ZWF activities in Δ*eda*Δ*gnd,* Δ*eda,* Δ*gnd,* and Δ*zwf* as a control and found to our surprise that ZWF activity was completely lacking in Δ*gnd* (Fig. 4B). Sequencing of the *zwf* gene in Δ*gnd*, revealed an insertion of an adenine in position 399, which results in a premature stop codon, and obviously completely abolishes ZWF activity in the Δ*gnd* mutant, whereas the mutants Δ*eda*Δ*gnd* and Δ*eda* lacked this insertion in their *zwf* genes (Fig. 4C). Immunoblots with specific antibodies against ZWF confirmed that ZWF is present in WT, Δ*eda* and Δ*eda*Δ*gnd* but is no longer expressed in Δ*gnd* (Fig. 4D). Whereas Δ*eda*Δ*gnd* showed ZWF activity that resembled WT levels, ZWF activity was reduced to approximately 60% in Δ*eda* (Fig. 4B). This observation is in line with the reduced photomixotrophic flux through the OPP shunt in this mutant that we had observed earlier (3). The construction of corresponding complemented mutants (Δ*eda::eda*) is underway to test whether these differences are indeed due to the absence of EDA.

In summary, these data demonstrate that the accumulation pattern of 6PG in Δ*eda*Δ*gnd,* Δ*eda* and Δ*gnd* is not caused by an active ED pathway, as previously assumed. Instead, the pattern can be explained by a secondary mutation in *zwf* in the Δ*gnd* mutant which abolishes 6PG production in the Δ*gnd* mutant.

### *In vitro* characterization of EDA (Sll0107)

Taken together, all data so far support the conclusion that the ED pathway is absent in the cyanobacterium *Synechocystis*. One remaining puzzling observation is the reported detection of KDPG in *Synechocystis* in one study using IC-ESI-MS/MS, which can be indicative of an active ED pathway, whereas another study using LC–MS/MS failed to detect this metabolite (1, 3). Analytical artifacts can never be fully excluded. To clarify whether EDA (Sll0107) might catalyze the reversible aldol condensation of pyruvate and GAP to KDPG, which could explain the detection of KDPG in the absence of EDD and to test for potential promiscuity of EDA with several substrates, the *sll0107* gene (KEGG annotation) was expressed in *E. coli*. Although SDS–PAGE and immunoblot analysis confirmed the expression of a ∼23 kDa protein, it could not be eluted from the Ni–TED resin, and crude extracts lacked detectable KDPG aldolase activity. These observations suggested expression of a non-functional or truncated protein. Genomic inspection of *sll0107 (eda*) revealed an alternative start codon (genome position 2,972,167) 72 nucleotides upstream of the annotated start codon (genome position 2,972,095) (Fig. 5). Comparative sequence and AlphaFold 3 analyses (26) using *E. coli* and *C. crescentus* KDPG aldolases (27) as a reference, showed that only the upstream-initiated longer sequence retained the N-terminal α-helix characteristic of class I KDPG aldolases (Fig. 5A).

**Figure 5.**
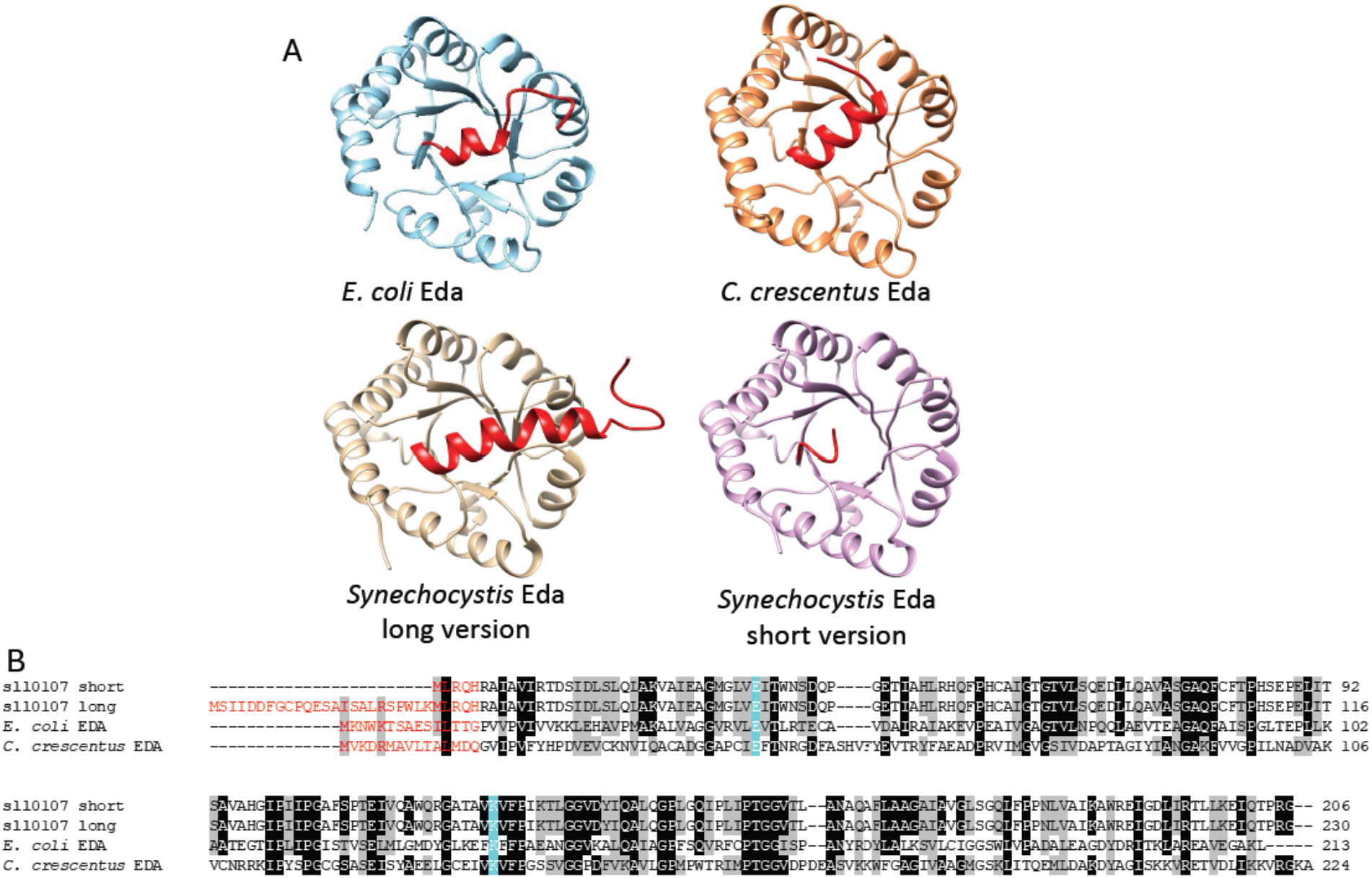
Structural comparison and sequence alignments of EDA enzymes including the genome-annotated version and an N-terminally extended version with an alternative start codon of the *Synechocystis* enzyme. (A) Structures of *E. coli* and *C. crescentus* EDA enzymes are compared with AlphaFold 3 models of the N-terminally extended (long) and genome-annotated (short) *Synechocystis* EDA variants. The conserved N-terminal α-helix is highlighted in red. Amino acid sequences were retrieved from KEGG, AlphaFold 3 models were generated using AlphaFold Server v3.0 (26) and structures were visualized and edited using UCSF ChimeraX. (B) Sequence alignment highlighting the N-terminal residues of the conserved α-helix in red. The catalytic lysine responsible for Schiff-base formation and the glutamate acting as the acid/base catalyst are indicated in cyan. The alignment was performed using Clustal Omega (28) and manually visualized in BioEdit.

Therefore, the extended Sll0107 version was expressed in *E coli*, purified and resulted in active protein (Fig. S10-S12). SDS–PAGE showed a subunit mass of ∼25 kDa, consistent with the calculated 23 kDa, and size-exclusion chromatography indicated a native mass of ∼64 kDa, corresponding to a homotrimeric assembly typical of bacterial class I KDPG aldolases (13, 29, 30).

Recombinant Sll0107 was assayed using the LDH-coupled continuous assay and exhibited robust activity toward its canonical substrate KDPG, catalyzing its cleavage into pyruvate and GAP with a catalytic efficiency of 24.4 s^-1^ mM^-1^ (Fig. 6, Table 2). Inspired by recent reports that metal-independent class I KDPG aldolases from the cyanobacterium *Synechococcus elongatus* also accept oxaloacetate (12), we tested this substrate and found that EDA (Sll0107) catalyzed its cleavage to pyruvate and CO₂ with a catalytic efficiency of 0.58 s^-1^ mM^-1^ (Fig. 6, Table 2). Since elevated proline levels were previously reported in Δ*eda*, we next examined activity with 2-keto-4-hydroxyglutarate (KHG), an intermediate of the proline degradation pathway as substrate (5, 31). EDA efficiently cleaved KHG to pyruvate and glyoxylate with a catalytic efficiency of 0.18 s^-1^ mM^-1^ (Fig. 6, Table 2).

**Figure 6.**
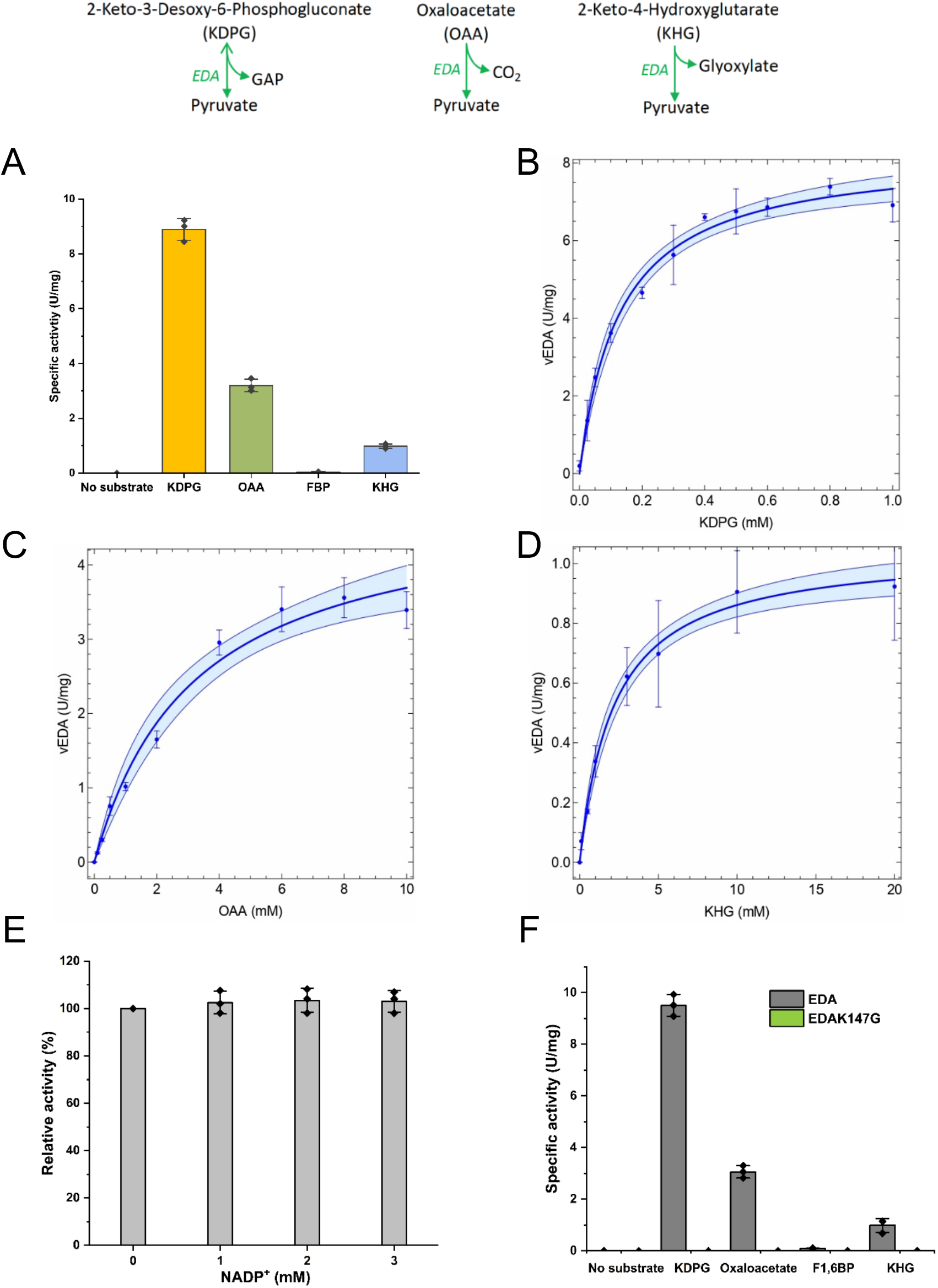
Substrate specificity and characterization of EDA (Sll0107) from *Synechocystis.* **(A)** Substrate specificity was assessed with 5 mM of each KDPG, oxaloacetate, FBP, and KHG using the continuous coupled assay (see Methods). **(B-D) Enzymatic characterization of EDA using KDPG, OAA and KHG as substrates.** The activity of EDA was measured across a range of (B) KDPG (0–1 mM), (C) OAA (0-10 mM) and (D) KHG (0-20 mM) concentrations. Colored bands indicate 95% confidence, and data are shown as mean ± standard deviations from three technical replicates. The kinetic parameters are given in table 2. **(E) Effect of NADP^+^ on EDA activity**. Different NADP^+^ concentrations (0, 1, 2 and 3 mM) were tested for their effect on EDA activity. Assays were performed with sub-saturating KDPG concentration (0.2 mM). The relative activity (%) in comparison to the control without effector (100%; specific activity of 4.3 U/mg) is shown. **(F) Effect of K147G mutation on EDA activity.** Enzymatic activity of wild-type EDA and the catalytic mutant EDAK147G was measured with 5 mM KDPG, oxaloacetate, FBP, and KHG using continuous coupled assays (see Methods). Data are presented as mean ± SD from three technical replicates (n = 3).

**Table 2:** Kinetic parameters of EDA (Sll0107) from *Synechocystis*.

| Substrates | $V_{max}$ | $K_m$ | $k_{cat}$ | $k_{cat}/K_m$ |
| --- | --- | --- | --- | --- |
| KDPG | $8.28 \pm 0.25$ U<br>mg<sup>-1</sup> | $0.13 \pm 0.01$<br>mM | $3.2$ s<sup>-1</sup> | $24.4$ s<sup>-1</sup> mM<sup>-1</sup> |
| OAA | $4.87 \pm 0.37$ U<br>mg<sup>-1</sup> | $3.19 \pm 0.64$<br>mM | $1.86$ s<sup>-1</sup> | $0.58$ s<sup>-1</sup> mM<sup>-1</sup> |
| KHG | $1.05 \pm 0.03$ U<br>mg<sup>-1</sup> | $2.18 \pm 0.24$<br>mM | $0.4$ s<sup>-1</sup> | $0.18$ s<sup>-1</sup> mM<sup>-1</sup> |
| FBP | No activity |  |  |  |
| Malate | No activity |  |  |  |
| Pyruvate + D-GAP | $0.18 \pm 0.01$ U mg<sup>-1</sup> (2.2% of the KDPG cleavage activity) | | | |
| Pyruvate + CO<sub>2</sub> | No activity |  |  |  |
| Pyruvate + glyoxylate | No activity |  |  |  |

EDA exhibited a strong preference for KDPG as its primary substrate but also catalyzed reactions with OAA and KHG, albeit with substantially lower catalytic efficiency. The effect of NADP⁺ on EDA activity was examined, as NADP⁺ has been reported to act as an activator of *S. elongatus* EDA (12). Under the assay conditions used in this study, the addition of NADP⁺ at concentrations up to 3 mM did not result in any measurable increase in EDA activity (Fig. 6E). These findings are consistent with curated enzyme data reported in BRENDA Enzyme Database (32), in which NADP⁺ is not listed as an activator for EDA.

To assess reaction reversibility, a hallmark of class I aldolases, reported for archaeal KD(P)G aldolases (15), we tested the enzyme in the condensation direction using (i) pyruvate + GAP for KDPG formation, (ii) pyruvate + CO₂ for oxaloacetate, and (iii) pyruvate + glyoxylate for KHG synthesis. Only a low level of reverse activity, 0.18 U/mg (∼2% of the cleavage activity) was detected for KDPG formation using a discontinuous TBA assay, whereas no reverse activity was observed for oxaloacetate or KHG synthesis (Fig. S12).

Given that a soybean KDPG aldolase was recently reported to accept fructose 1,6-bisphosphate (FBP) (2) we also tested FBP as a substrate. However, EDA (Sll0107) displayed no detectable activity with FBP (Fig. 6A, 6F, Table 2). Also, malate was not used as substrate (Table 2).

Class I KDPG aldolases are metal-independent enzymes that utilize a catalytic lysine residue to form a Schiff base intermediate during catalysis. In EDA this lysine (Lys147, based on the extended *sll0107* sequence) was identified as essential for activity toward multiple sugar acid substrates (33). To generate a catalytically inactive variant, Lys147 was substituted with glycine (K147G), which lacks both side-chain and net charge. The mutant enzyme EDAK147G was expressed and purified at comparable levels to the WT protein but exhibited no detectable activity with any tested substrate (Fig. 6F).

Taken together, these data show that the KDPG aldolase EDA (Sll0107) of *Synechocystis* is an aldolase with substrate promiscuity, which *in vitro* cleaves KDPG to GAP and pyruvate, OAA to CO2 and pyruvate, and KHG to glyoxylate and pyruvate (Fig. 6, Table 2). In addition, it synthesizes KDPG from GAP and pyruvate with very low efficiency (∼2% of the cleavage activity) (Fig. S12), which might explain the detection of KDPG in our initial study (1).

Since 6PG dehydratase (EDD) seems to be absent in *Synechocystis*, the canonical substrate of EDA, 2-keto-3-deoxy-6-phosphogluconate (KDPG) is not produced, and no alternative enzymatic source of KDPG is known to the best of our knowledge. Our *in vitro* enzyme assays therefore suggest that EDA may instead participate in alternative metabolic reactions, potentially linked to the PEP-pyruvate-OAA node and/or proline catabolism, as illustrated in Figure 7. However, further studies are required to clarify the physiological role of EDA *in vivo*.

**Figure 7:**
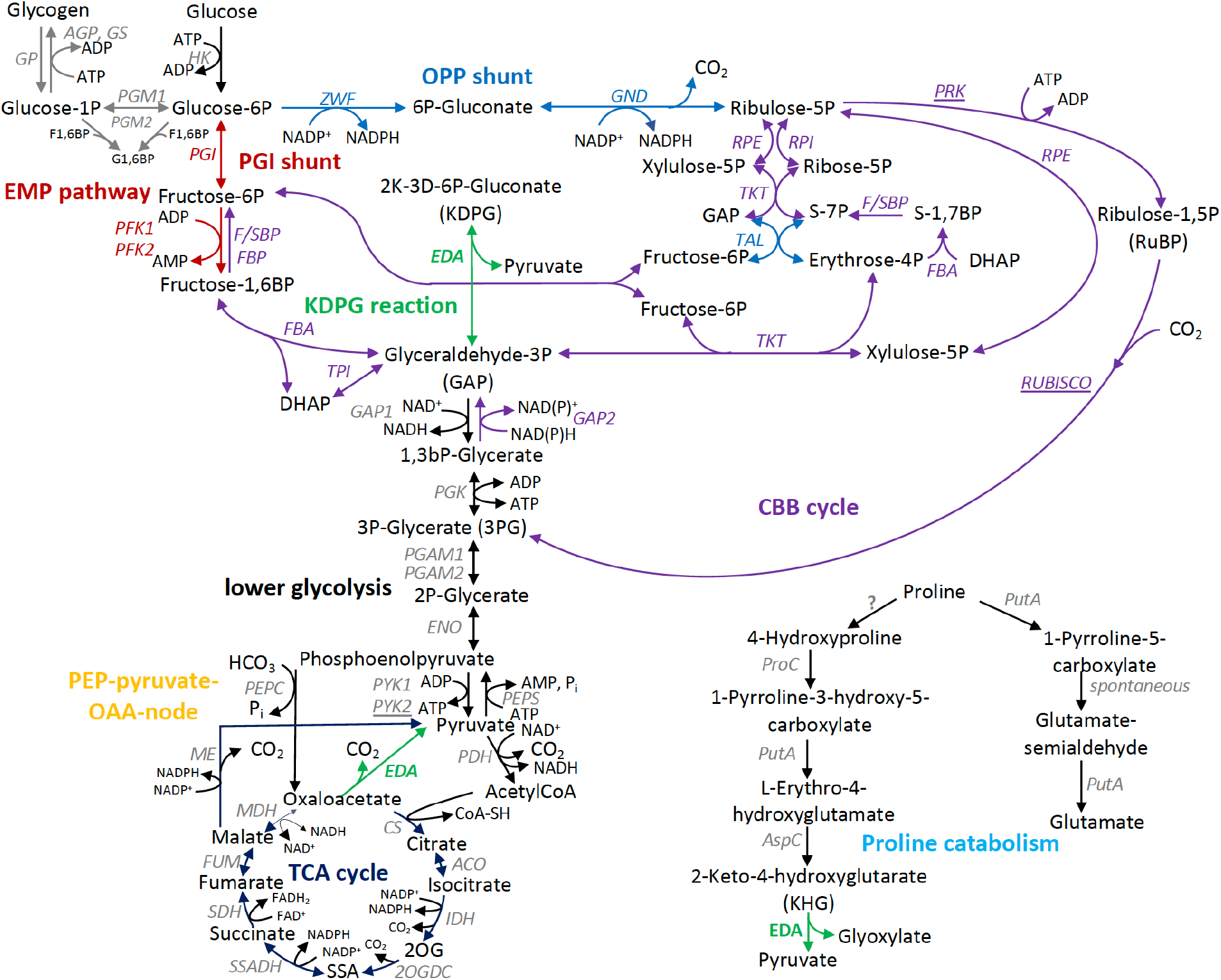
Proposed metabolic map of the central carbon metabolism in *Synechocystis* with all suggested *in vivo* roles of the promiscuous aldolase Sll0107 (EDA) highlighted in green, based on substrate promiscuity and activities observed *in vitro*. ACO, Acontinase; AGP, ADP-glucose pyrophosphorylase; CBB, Calvin–Benson–Bassham; DHAP, dihydroxyacetone phosphate; E4P, erythrose 4-phosphate; EMP, Embden–Meyerhof–Parnas; EDA, Enter-Doudoroff aldolase; ENO, enolase; F6P, fructose 6-phosphate; FBA, fructose-bisphosphate-aldolase; FBP, fructose 1,6-bisphosphate; FBPase, fructose-1,6-bisphosphatase; F/SBPase, fructose-1,6-bisphosphatase/sedoheptulose-1,7-biphosphatase; FUM, fumarase; GAP, glyceraldehyde 3-phosphate; GAPDH, glyceraldehyde-3-phosphate dehydrogenase; GBP, glucose 1,6-bisphosphate; GND, 6-phosphogluconate dehydrogenase; GP, glycogen phosphorylase; GS, glycogen synthase; HK, hexokinase; IDH, isocitrate dehydrogenase; ME, malic enzyme, MDH, malate dehydrogenase; OPP, oxidative pentose phosphate; PDH, pyruvate dehydrogenase complex, PEPC, phosphoenolpyruvate carboxylase, PEPS, phosphoenolpyruvate synthetase, PFK, phosphofructokinase; PGAM, phosphoglycerate mutase; PGI, phosphoglucose isomerase; PGK, phosphoglycerate kinase; PGL, 6-phosphogluconolactonase; PGM, phosphoglucomutase; PRK, phosphoribulokinase; PUTA, proline dehydrogenase; PYK, pyruvate kinase; R-5P, ribose 5-phosphate; RPE, ribulose-5-phosphate epimerase; RPI, ribose-5-phosphate isomerase; SDH, succinate dehydrogenase; TalB, transaldolase; TKT, transketolase; TPI, triosephosphate isomerase; X-5P, xylulose 5-phosphate; ZWF, glucose-6-phosphate dehydrogenase.

## Conclusions

Our data demonstrate that *Synechocystis* lacks a functional ED pathway, due to the absence of both EDD and the proposed GDH/GK bypass, which is also supported by photomixotrophic flux analyses (3). Instead, it possesses a promiscuous ED aldolase, whose physiological role remains to be elucidated. The dehydratases EDD and DHAD have a common origin, a high degree of amino acid and structural similarity, and the same catalytic mechanism. However, they differ in two domains that contribute to substrate selection, which is why EDD and DHAD are enzymes with high substrate specificity that cannot replace one another (2). Apart from four identified cyanobacteria (*Nostoc* sp. 335mG, *Synechococcus moorigangali* CMS01, *Leptolyngbya valderiana* BDU 20041, and *Leptolyngbya* sp. 15MV) that possess potential EDDs (based on the described domains, PXA97457.1 for *Nostoc*; MBV5257376.1 for *S. moorigangali*; OAB60313.1 for *L. valderiana*; QYU66659.1 for *Leptolyngbya.* sp), most cyanobacteria lack EDD despite the presence of EDA, which likewise applies to algae, moss and plants (2, 31). Our data demonstrate that Slr0452 in *Synechocystis*, which we previously assumed as EDD (1), lacks 6PG dehydratase activity and instead dehydrates DHIV exclusively, consistent with recent reports (17). The same experimental results were recently obtained for the cyanobacterium *S. elongatus* PCC 7942 that also lacks EDD but contains EDA (12). This raises the question of what role EDA could play in the absence of EDD and the ED pathway in cyanobacteria, moss, algae and plants. In soybean (*Glycine max*), GmEDA was found to catalyze three reactions *in vitro*: firstly, the cleavage of KDPG to GAP and pyruvate, secondly the cleavage of fructose 1,6-bisphosphate (FBP) to dihydroxyacetone phosphate (DHAP) and GAP and thirdly the condensation of erythrose 4-phosphate (E4P) and GAP to sedoheptulose 1,7-bisphosphate (SBP) (2). Accordingly, the idea was put forward that GmEDA is a promiscuous aldolase that catalyzes aldolase reactions in the CBB cycle and glycolysis *in vivo*, which is well in line with its localization in chloroplasts (2). In diatoms the ED pathway was suggested to operate in mitochondria providing GAP and pyruvate to lower glycolysis and the TCA cycle, based on bioinformatic analyses, bacterial complementation experiments and measurements on the conversion of 6PG to pyruvate in soluble protein extracts (34–36). However, as neither the EDD nor EDA from diatoms were thoroughly characterized biochemically yet, further studies would be informative. The proteomes of three marine *Prochlorococcus* and three marine *Synechococcus* strains were investigated upon exposure to glucose (37). Protein levels of EDA were either downregulated or not effected while proteins involved in OPP shunt and CBB cycle were upregulated indicating their involvement under photomixotrophy in these cyanobacteria possibly similar to *Synechocystis* (3, 4, 37). However, flux analyses are still missing for the marine strains.

Based on our *in vitro* data *Synechocystis* EDA might be connected to the TCA cycle and the PEP-pyruvate-OAA node by decarboxylating oxaloacetate to pyruvate and/or in proline catabolism by splitting KHG to glyoxylate and pyruvate (Fig. 7). The latter assumption is supported by earlier findings of elevated proline levels in Δ*eda* mutants of *Synechocystis* (5). For EDA from *S. elongatus* PCC 7942, it has recently been demonstrated that, in addition to cleaving KDPG, it also decarboxylates oxaloacetate (12). In contrast to *Synechocystis* EDA, EDA from *S. elongatus* PCC 7942 prefers OAA over KDPG. The catalytic efficiency of *S. elongatus* PCC 7942 EDA on oxaloacetate is rather low (OAA 0.437 s^-1^ mM^-1^), however, its activity can be enhanced by NADP^+^(12). The *Synechocystis* EDA showed a 42-fold higher catalytic efficiency with KDPG compared to oxaloacetate (KDPG 24.4 s^-1^ mM^-1^, OAA 0.58 s^-1^ mM^-1^) and no activation by NADP^+^ (1-3 mM) was observed. As the *S. elongatus* PCC 7942 Δ*eda* mutant was compromised in terms of photosynthesis and growth under diurnal day/night cycles, the authors hypothesized that *S. elongatus* PCC 7942 EDA might play a role in balancing NADPH/NADP^+^ ratios and photosynthesis by modulating TCA cycle activities (12). For *Synechocystis* EDA, the catalytic efficiency for KDPG compared to KHG was 136-fold higher (KDPG 24.4 s^-1^ mM^-1^, KHG 0.18 s^-1^ mM^-1^). However, EDA displayed no detectable activity with FBP (Fig. 6), which is consistent with the fact that class I KDPG aldolases are members of a different enzyme class (EC 4.1.2.14). Likewise, class I and class II FBP aldolases (EC 4.1.2.13) are not known to exhibit activity toward KDPG (16, 30, 38).

Interestingly, EDA was able to condense GAP and pyruvate to KDPG *in vitro* with low efficiency. It remains to be investigated whether this reaction could play a role *in vivo*, with KDPG potentially acting as a regulatory metabolite at low concentrations. This could explain why KDPG was detected in WT and Δ*pfk*Δ*zwf* in one study (1), although this observation could not be replicated later (3). Based on the current data, it is not possible to determine whether the detected KDPG was due to a measurement artifact or not. The accumulation pattern of 6PG in WT, Δ*eda*, Δ*gnd* and Δ*eda*Δ*gnd*, which contradicted the flux analyses and had been interpreted as evidence of an active ED metabolic pathway in *Synechocystis* (3, 4), could be explained by a secondary mutation in the Δ*gnd* mutant (Fig. 4C) and now agrees with the flux analyses.

Our systematic re-examination of data on the proposed ED pathway in *Synechocystis* demonstrates how complementary approaches can contribute to elucidating cyanobacterial metabolism. By combining detailed biochemical enzyme characterization, genetic deletion and complementation, careful mutant validation, and taking into account metabolic flux analyses (3), we were able to clarify that neither EDD nor the GDH/GK bypass is present, and that the ED pathway is incomplete and not functional in *Synechocystis*. These integrated studies further revealed the presence of a promiscuous KDPG aldolase whose broader metabolic involvement now opens exciting new research directions.

This study also highlights the importance of best practices to strengthen metabolic pathway reconstruction and provides the following lessons: Although protein sequence alignments provide valuable information, they cannot replace detailed biochemical enzyme tests, which are required to verify the predicted enzyme functions and to further test for substrate promiscuity. Furthermore, to demonstrate the presence of a metabolic pathway, it is advisable to characterize all enzymes of this pathway biochemically, rather than limiting analysis to individual key enzymes, and to also include flux analyses. Routine complementation of deletion mutants and sequencing of selected genes or the entire genome are effective means of identifying secondary mutations that can lead to misleading phenotypes (39). A final lesson is to exercise great caution when interpreting data and drawing conclusions, even if the data sets appear completely logical and conclusive at first glance (1). Over time, contradictory results have accumulated regarding the claim that there is an ED pathway in *Synechocystis*. In the present study, we have succeeded in clarifying a number of these contradictions. Taken together, these advances provide a solid foundation for future investigations into the *in vivo* role of the promiscuous aldolase EDA in cyanobacteria.

## Materials and Methods

### Growth experiments

*Synechocystis* sp PCC 6803 WT (glucose tolerant strain) and mutants (Table S2) were grown in 200 ml BG11 medium (pH 8) in custom-made glass tubes placed in a photobioreactor (manufactured by Willi Hilke, Uslar, Germany), at 28°C under constant light (50 µmol m^-2^ s^-1^). Standard BG11 medium contained: 17.6 mM NaNO3, 5 mM Tris, 0.304 mM MgSO4, 0.245 mM CaCl2, 0.189 mM NaCO3, 0.175 mM K2HPO4, 46.3 µM H3BO3, 31.2 µM citric acid, 22.8 µM FeNH4 citrate, 9.1 µM MnCl2, 2.79 µM Na2EDTA, 1.61 µM NaMoO4, 0.77 µM ZnSO4, 0.32 µM CuSO4, 0.17 µM Co(NO3)2 according to (40). FeNH4 citrate was added after autoclaving. Cultures were gassed with filter-sterilized ambient air. Photomixotrophic cultures were supplemented with 10 mM D-glucose. Aliquots were taken under sterile conditions every 24 h to measure optical density photometrically at 750 nm.

### Generation of mutants

Constructs for deletion mutants, complementation of Δhk and edaK147G were made by Gibson Assembly cloning (41), using the primers listed in Table S3. An antibiotic resistance cassette the was placed in the gene to be deleted and was fused for homologous recombination to two DNA fragments approximately 200 bp directly up- and downstream of the gene and cloned into pBluescript. *Synechocystis* was transformed with these constructs to replace the gene with the antibiotic resistance cassette via homologous recombination. Mutants were checked via PCR for segregation.

### Generation of a putative GDH1 (sll1709) His-tagged over-expression mutant (sll1709:oe) and overexpression of GDH homologues of Spirulina and Lyngbya

Two DNA fragments from the genome of *Synechocystis* were synthesized by Genescript: 1. a fragment containing 212 bp up- and 212 bp downstream of the s*ll1709* start codon, with a *BamHI*, *XhoI* and *NdeI* site in its middle and 20 bp of overlapping sequences with the pBluescript SK(+) vector at its ends (Fig. S5A). 2. a fragment containing a modified petE promotor, followed by His-tag, TEV cleavage recognition and linker encoding sequences, various restriction sites and 20 bp of overlapping sequences with the pBluescript SK(+) vector at its ends (Fig. S5B). Both fragments were separately cloned into the pBluescript SK(+) vector by Gibson Assembly cloning. A kanamycin antibiotic resistance cassette was inserted into the *EcoRV* site of the plasmid containing the modified petE promotor. Both plasmids were then digested with *BamHI* and *NdeI* and the promoter cassette was ligated into the alkaline phosphatase treated GDH1 plasmid to yield the final mutagenesis vector. This plasmid was transformed into *Synechocystis* and mutant segregation was confirmed by PCR analysis. GDH homologues of *Spirulina* and *Lyngbya* were overexpressed in *E.coli* (Fig. S7-8).

### P3-His-GDH1 (sll1709) over-expression and His-tag purification

For the purification of GDH1 from *Synechocystis*, a 6 L photoautotrophic culture of the P3-His-GDH1 overexpression strain was grown in a 10 L glass flask at 28°C, illuminated with constant light (50 µmol m^-2^ s^-1^) and gassed with filter sterilized ambient air to an OD750 of about 1. Cells were harvested by centrifugation at a RCF (relative centrifugal force) of 3,992 x g in a Beckman Coulter with a JLA-8.1000 Rotor for 20 minutes at 4°C. Initially, His-GDH1 over-expression in the 6 L culture was assessed by SDS PAGE analysis followed by immunoblotting with a His-tag specific antibody (Fig S6) as described in (42). A specific band could be detected in the over-expression mutant, confirming expression and stable accumulation of the over-expressed and N-terminally His-tagged GDH1 protein. A small-scale purification (using a 50-mL aliquot of the large culture and treating this sample as described below for the remaining sample) demonstrated that the protein could be purified (Fig S6C). For the large-scale purification the remaining cells were resuspended in lysis buffer (50 mM NaPO4 pH=7.0; 250 mM NaCl; 1 tablet complete protease inhibitor EDTA free (Roche) per 50 mL) and broken by passing them through a French Press cell at 1250 p.s.i. twice. Unbroken cells and membranes were removed by centrifugation in a Beckman ultracentrifuge using a 70 Ti rotor at 30.000 rpm for 45 min at 4°C. The decanted soluble extract was adjusted to a volume of 45 mL with lysis buffer and incubated with 5 mL TALON cobalt resin (Takara) for 1 h at 4°C. The resin was then washed extensively with 200 mL lysis buffer and subsequently with 100 mL lysis buffer containing 5 mM imidazole. Bound proteins were eluted with 10 mL elution buffer (50 mM NaPO4 pH=7.0; 250 mM NaCl; 250 mM imidazole) and after dialysis overnight against the reaction buffer 50 mM Tris pH=7.5; 20 mM MgCl, the protein was concentrated in a Vivaspin Turbo 4 Ultrafiltration Unit (5 kDa MWCO). Commercially available Glucose dehydrogenase from *Pseudomonas* sp. (19359, Sigma) was resuspended in the reaction buffer for control measurements. Finally, protein concentrations were determined by Bradford assay using the Roti-Quant reagent (Carl Roth) and adjusted to 0.25 mg/mL. Samples taken during the purification procedure were assessed by SDS PAGE analysis followed by immunoblotting with antibodies specific against the His-tag.

### Preparation of crude cell extracts

Enzyme activity was measured in crude cells extracts. For the generation of crude cell extracts of *Synechocystis*, an OD volume of 30-50 OD*mL sample culture was pelleted by centrifugation (5 min, 9000 *g*, 4 °C) and the pellet resuspended in 0.5 mL 100 mM Tris buffer (pH 7.6). The suspension was then transferred to centrifugation tubes with screwed lids (max 500 µl/cup) and glass beads (0.17-0.18 mm diameter; Sartorius, Göttingen, Germany) were added so that 1-2 mm liquid remained above the dispersion. Samples were vortexed for 10 min at 4 °C. Afterwards, the glass beads were pelleted by centrifugation in two consecutive centrifugation steps (1 min, 1000 *g*, 4 °C and 10 min, 1600 *g*, 4 °C). The supernatant was transferred to new centrifugation cups in between centrifugations. In order to precipitate cell debris, the supernatant was centrifuged (15 min, 18000 *g*, 4 °C) one more time. The supernatant constituted the final cell extract and was measured for its protein concentration and kept on ice until further enzyme activity measurements.

### Protein quantification

Protein concentrations were determined according to Bradford (43). 1 μL protein sample (cell extract) was mixed in triplicates with 799 μL water. Afterwards, 200 μL Bradford reagent, consisting of 100 mg/L Coomassie Serva Blue G-250 (Bio-Rad, Munich, Germany), 5 % (v/v) ethanol and 8.5 % (v/v) H3PO4 was added to all samples as well as to 800 µL bovine serum albumin (BSA) standards in concentrations of 2, 5, 10, 15 and 20 μg/mL. After an incubation at RT for 10 min, absorption at 595 nm was measured and sample protein concentration calculated by using the standard curve.

### Glucose dehydrogenase activity measurements

For glucose dehydrogenase (GDH) activity measurements in *Synechocystis*, NADPH turnover was monitored photometrically at 340 nm. Measurements were carried out in 50 µl crude cell extract or purified GDH in 100 mM Tris/HCl buffer (pH 7.4) in the presence of 20 mM glucose, 5 mM NAD(P)^+^ and 10 mM MgCl2. Reactions were started by addition of glucose. The kinetics of the reaction were monitored with a double-beam spectrophotometer (Uvikon 810, Kontron, Augsburg, Germany) for 5-10 minutes. In case quinones were tested as electron acceptors, the GDH reaction was enzymatically coupled to the gluconate kinase (GK) and 6PG dehydrogenase (GND) reaction, the latter providing the required NADP^+^ turnover. This was achieved by adding 5 mM ATP, 0.02 U gluconate kinase from *E. coli* (Megazyme, Wicklow, Ireland) and 0.005 U 6-phosphogluconate dehydrogenase from *E. coli* (Megazyme, Wicklow, Ireland). The two tested quinones 2,3-dimethoxy-5-methyl-1,5-benzoquinone (DMB) and duroquinone (DQ) were supplied in a concentration of 5 mM. As a positive control, 0.05 U glucose dehydrogenase from *Pseudomonas sp.* (Sigma Aldrich, Steinheim, Germany) was added.

The GDH activity of recombinant enzymes from *Spirulina* sp. SIO3F2 and *Lyngbya* aestuarii was determined after GST (Glutathione S-Transferase) - His tag removal by TEV protease cleavage and purification. Activity was determined spectrophotometrically by monitoring glucose dependent reduction of NAD(P)^+^ to NAD(P)H at 340 nm and 30°C (extinction coefficient of NAD(P)H=6.22 mM^-1^cm^-1^). The standard assay mixture (final volume of 500 µL) contained 0.1 mM Tris/HCl (pH 7.7 at 30°C), 10 mM MgCl2, 5 mM NAD(P)^+^, 20 mM Glucose, and 2.5 – 5 µg of recombinant protein. To determine the substrate specifity glucose was substituted by other sugars (for details see text).

### Gluconate kinase activity measurements

For gluconate kinase (GK) activity measurements in *Synechocystis* crude cell extracts the GK reaction was enzymatically coupled to 6-phosphogluconate dehydrogenase (GND) reaction, the latter providing NADP^+^ reduction, which was monitored photometrically at 340 nm. Measurements were carried out in 50 µl crude cell extract in 100 mM Tris buffer (pH 7.4) in the presence of 20 mM gluconate, 5 mM NADP^+^, 10 mM MgCl2., 5 mM ATP and 0.005 U 6-phosphogluconate dehydrogenase. Reactions were started by addition of gluconate and kinetics were monitored with a double-beam spectrophotometer (Uvikon 810, Kontron, Augsburg, Germany) for 5-10 minutes. As a positive control, 0.05 U gluconate kinase from *E. coli* (Megazyme, Wicklow, Ireland) was added.

Recombinant GK activity in *E. coli* was determined spectrophotometrically in a continuous coupled pyruvate kinase/lactate dehydrogenase (PK/LDH) assay by monitoring NADH oxidation at 340 nm (30 °C) (extinction coefficient of NAD(P)H=6.22 mM^-1^cm^-1^). Gluconate phosphorylation was followed indirectly via ADP formation, which was converted by PK/LDH in a NADH-dependent reaction. The standard assay (500 µl) contained 100 mM Tris/HCL (pH 7.7), 10 mM MgCl2, 0.2 mM NADH, 1 mM PEP, 2 mM ATP, 1-10 mM gluconate, auxiliary enzymes PK and LDH (1U each), and 0.7-15µg purified recombinant enzyme (after GST-tag cleavage).

### Glucokinase/hexokinase activity measurements

For glucokinase/hexokinase (GLK/HK) activity measurements, the GLK/HK reaction was enzymatically coupled to the glucose-6-phosphate dehydrogenase (ZWF) reaction, whose NADP^+^ turnover was monitored photometrically at 340 nm. Measurements were carried out in 50 µl crude cell extract in 100 mM Tris buffer (pH 7.4) in the presence of 10 mM glucose or fructose, 5 mM NADP^+^, 10 mM MgCl2., 5 mM ATP and 0.005 U 6-phosphogluconate dehydrogenase from *E. coli* (Megazyme, Wicklow, Ireland). Reactions were started by addition of glucose and kinetics were monitored with a double-beam spectrophotometer (Uvikon 810, Kontron, Augsburg, Germany) for 5-10 minutes.

### Quantification of 6-phosphogluconate

Quantification of 6-phosphogluconate (6PG) was achieved photometrically by enzymatic conversion of sample 6PG to ribulose-5-phosphate and CO2, which goes hand in hand with equimolar formation of NADPH. 5 to 50 OD*mL cell culture was pelleted by centrifugation (5 min, 9000 g, 4 °C) and cell lysis was achieved by solving the pellet in 1 mL 0.2 M HCl, followed by a 15 min incubation at 95 °C. The lysates were centrifuged (10 min, 18000 g, 4°C) and the supernatants transferred to new tubes where they were neutralized by addition of 1 mL 1 M Tris/HCl buffer (pH 8.0). Samples were then split into two (each sub-pool 900 μL), that were supplied with 90 μL of an 11.11 mM NADP^+^ solution. 10 μL of a 5 U/mL 6-PG dehydrogenase from (Megazyme, Wicklow, Ireland) solution was added to one of the samples, whereas 10 μL of water was added to the second sample as a control. All samples were then incubated at 37 °C for 30 min and the absorption at 340 nm was measured using a double-beam spectrophotometer (Uvikon 810, Kontron, Augsburg, Germany). 6PG standards were prepared (0 mM, 0.05 mM, 0.1 mM, 0.2 mM, 0.4 mM and 0.8 mM) and measured in the same way as samples. Δabsorption values at 340 nm of standards were used to create a standard curve, which was then used to convert sample Δabsorption values into 6PG concentrations. The resulting 6PG concentration (mM) was converted into specific cellular 6PG content (μg/OD*mL) by multiplying it with the molar mass and dividing it by the respective OD volume.

### Quantification of glucose

For quantification of medium glucose concentrations, 1 mL *Synechocystis* culture was pelleted by centrifugation (5 min, 10000 g, RT) and 50 μL of supernatant were transferred to a new cup. If medium glucose concentration was expected to exceed 5 mM, samples were diluted. 50 µL glucose standards with the following concentrations were prepared: 0 mM, 0.625 mM, 1.25 mM, 2.5 mM and 5 mM. Furthermore, a ‘NADP buffer’, containing 150 mM NADP^+^, 150 mM ATP, 300 mM MgCl2 in 100 mM Tris-HCl (pH 7.4,) was prepared. 940 μL ‘NADP buffer’ were added to all samples and standards, which were then transferred to acryl-cuvettes and absorption at 340 nm was measured in a double-beam spectrophotometer (baseline absorption) (Uvikon 810, Kontron, Augsburg, Germany). After that, 0.17 U Hexokinase as well as 0.085 U glucose-6-phosphate dehydrogenase were added to all samples and standards. Samples and standards were then incubated at 37 °C for 60 min to ensure complete enzymatic conversion. Finally, absorption at 340 nm was measured again (endpoint absorption). Baseline absorbance values were subtracted from endpoint readings to calculate ΔA₃₄₀. A glucose standard curve was generated from the standards and used to determine glucose concentrations in the culture supernatants.

### Blast analysis

In order to search for the existence of a gluconate kinase in the genome of *Synechocystis* and cyanobacteria in general, the fasta protein sequences of the well characterized gluconate kinases from *Escherichia coli* (KIG95398.1) and from *Pseudomonas aeruginosa* (WP134279162.1) were used as baits for the protein/protein blast tool at NCBI. Similarly, the fasta sequences of the NAD^+^ dependent glucose dehydrogenase 1GCO from *Bacillus megaterium* and the quinone dependent glucose dehydrogenase 1C9U from *Acinetobacter calcoaceticus* were used as baits for the search of gluconate dehydrogenases in cyanobacteria. Next to *Synechocystis*, 261 cyanobacterial species were analyzed. Homology thresholds were set at a query score > 30 %, an identity score > 30 % and an e-value < 1^-10^.

## SDS PAGE and immunoblot analyses

To assess the level of ZWF protein in various strains, soluble protein extracts were prepared and analysed by SDS-PAGE and immunoblotting. 5 µg of protein were loaded per lane on a 12.5% (w/v) polyacrylamide BisTris gel using MES running buffer (250 mM MES, 250 mM Tris, 5 mM EDTA, 0.5 % (w/v) SDS). Gels were Coomassie-stained or electroblotted onto nitrocellulose membrane. For immunoblotting, 5% (w/v) milk powder in 1x PBS-T was used as blocking solution and all washing steps were performed with 1x PBS-T. For detection a ZWF antibody (αZwf, Agrisera, Sweden, Phyto AB, PHY5113A) in a 1:5k dilution in 1x PBS-T (phosphate-puffered saline with Tween-20) and a horseradish peroxidase-conjugated secondary antibody (anti-rabbit, 1:10k in 1x PBS-T Cytiva (phosphate-buffered saline with Tween 20, 0.1 µm sterile filterized), Thermo Fisher Scientific, Germany) were used. As a loading control an antibody against 30S ribosomal protein S1 (αRps, Agrisera, Sweden) was utilized.

## Gene cloning and protein overexpression

Genes were codon-optimized for expression in *E. coli*. Codon-optimized genes from *Synechocystis* (*eda*, *sll0107*, longer *sll0107* variant, extended by 24 amino acids, *edd/dhad*, *slr0452*) and *Synechococcus elongatus* PCC 7942 (*dhad*, *syc0898_c*) and *Caulobacter crescentus* (*eda*, *ccna_0156* and *edd*, *ccna_02134*) for expression in *E. coli* were synthesized by BioCat GmbH (Germany). The *dhad* gene from *Sulfolobus solfataricus* (*sso3107*), encoding a DHAD with GAD activity was synthesized by Eurofins (Eurofins Genomics, Ebersberg, Germany). The *sll0107* and longer *sll0107* variant were inserted into pet15B vectors encoding an N-terminal 6xHis-tag, while *slr0452* and *syc0898_c* genes were inserted into pET24a vectors encoding a C-terminal 6×His-tag. The *saci_0225* gene from *Sulfolobus acidocaldarius* (encoding KDPG aldolase) was cloned into the pET15b vector using *NdeI* and *BamHI* restriction sites. The primers used for cloning are listed in Table S3. Genes encoding *eda* (*ccna_01562*) and *edd* (*ccna_02134*) from *C. crescentus* were cloned into pET28b and pET15b using *NdeI*/*BamHI* and *NdeI*/*HindIII*, respectively. All constructs were verified by Sanger sequencing (LGC Genomics, Berlin, Germany). For protein expression, plasmids were transformed into *E. coli* Rosetta (DE3) (Agilent Technologies). Cultures were grown in Terrific Broth (TB; 22 g L⁻¹ yeast extract, 12 g L⁻¹ tryptone, 4 mL L⁻¹ glycerol) supplemented with the appropriate antibiotics (100 µg mL⁻¹ ampicillin or kanamycin, and 30 µg mL⁻¹ chloramphenicol) at 37 °C with shaking at 180 rpm. Protein expression was induced at an OD₆₀₀ of 0.6–0.8 by addition of 1 mM isopropyl-β-D-thiogalactopyranoside (IPTG), followed by incubation at 18 °C for 18–22 h. Cells were harvested by centrifugation (15 min, 8630 × *g*, 4 °C) and stored at −70 °C until further use.

## EDA (KDPG aldolase) mutation at the active center

Lysine at position 147 in EDA (Sll0107) was substituted with glycine (K147G). Site-directed mutagenesis was performed as described (44). Q5® High-Fidelity DNA Polymerase and its recommended reaction buffer (NEB) were utilized for the amplification process. The primers used for mutagenesis are listed in Table S3, with the nucleotide substitutions introducing the K147G exchange underlined. As a template the codon-optimized *eda* gene from *Synechocystis* (*sll0107*) that was synthesized by BioCat GmbH (Germany), was utilized. Successful mutagenesis was confirmed by DNA sequencing (LGC genomics, Berlin, Germany). The resulting EDA variant was named EDAK147G.

## Protein purification

Recombinant *Synechocystis* EDA (both versions), EDAK147G, and DHAD (Slr0452), recombinant *Caulobacter crescentus* CcEDA and CcEDD, *Sulfolobus acidocaldarius Saci*KD(P)GA, and *Sulfolobus solfataricus SsoDHAD* were purified by immobilized metal affinity chromatography (IMAC) followed by size-exclusion chromatography (SEC) for EDA and Slr0452. Frozen cell pellets (2.4 g wet weight) were resuspended in 50 mM HEPES–NaOH (pH 7.8, 30 °C), 300 mM NaCl at a ratio of 1 g wet cells per 3 mL buffer. For EDD/DHADs purification buffer was supplemented with 10 mM MgCl2 and 1 mM DTT. Cells were disrupted by sonication (3 × 5 min; amplitude 50; cycle 0.5) using a UP200S homogenizer (Hielscher Ultrasonics, Brandenburg, Germany). Cell debris was removed by centrifugation (45 min, 21,130 × *g*, 4 °C). His-tagged proteins, except *Synechocystis* DHAD (Slr0452), were purified from the supernatant using Ni–TED columns (Macherey-Nagel, Düren, Germany) according to the manufacturer’s protocol. *Synechocystis* DHAD (Slr0452) was purified on a self-packed Ni–NTA affinity column. Elution fractions containing the target protein were pooled and concentrated using Vivaspin 20 centrifugal concentrators (Sartorius Stedim Biotech; 30 kDa cutoff for EDD/DHAD and 3 kDa cutoff for EDA). Concentrated samples of *Synechocystis* EDA (6 mg protein) were further purified by SEC on a HiLoad 16/600 Superdex 200 prep grade column (Cytiva, Marlborough, MA, USA) equilibrated with 50 mM HEPES–NaOH (pH 7.8, 30 °C), 300 mM NaCl and additionally contained 10 mM MgCl2 and 1 mM DTT for DHADs purification. Protein-containing fractions (4.5 mg after SEC) were analyzed by SDS–PAGE and enzymatic activity assays. Purified proteins were stored at −70 °C in 25% (v/v) glycerol. Protein concentrations were determined by the Bradford assay (45) using bovine serum albumin (Merck, Darmstadt, Germany) as the standard.

## Determination of the native molecular mass of EDA

Following IMAC purification, as stated above, the column was calibrated under identical buffer conditions using standard proteins from the LMW and HMW gel-filtration calibration kits (Cytiva, Marlborough, MA, USA): Aprotinin (6.5 kDa), Ribonuclease A (13.7 kDa), Ovalbumin (44 kDa), Aldolase (158 kDa), and Ferritin (440 kDa), with blue dextran to determine the column void volume. The native molecular mass of EDA was estimated from the resulting calibration curve.

## *In vitro* EDA enzyme characterization

The catalytic activities of *EDA*, EDAK147G, *Saci*KD(P)GA, and *Cc*EDA in the KDPG cleavage direction were determined using a continuous, coupled spectrophotometric assay. EDA catalyzes the cleavage of 2-keto-3-deoxy-6-phosphogluconate (KDPG) into pyruvate and glyceraldehyde 3-phosphate (GAP). Pyruvate formation was monitored indirectly by its reduction to lactate using lactate dehydrogenase (LDH; rabbit muscle, Merck, Darmstadt, Germany), coupled to the oxidation of NADH to NAD⁺. The decrease in absorbance at 340 nm was recorded in 96-well plates (BRANDplates®, BRAND, Wertheim, Germany) using a Tecan Infinite M200 plate reader (Tecan Group AG, Männedorf, Switzerland) at 30 °C. NADH concentration was quantified against a standard curve ranging from 0–0.7 mM.

For KDPG cleavage assays (200 µL final volume), reactions contained 0.1 M HEPES–NaOH (pH 7.8, 30 °C), 0.7 mM NADH, 2 mM KDPG, 2 U LDH, and the indicated enzyme (typically 1 µg EDA, EDAK147G, *Saci*KD(P)GA, or 0.06 µg *Cc*EDA). Substrate specificity toward oxaloacetate (OAA) and 2-keto-4-hydroxyglutarate (KHG) was assayed under the same conditions as KDPG assay, as they all generate pyruvate as one of the two product, however, for the KHG assay, 6 µg of EDA was used.

FBP cleavage activity forming GAP and DHAP was assayed using a coupled system containing glycerol-3-phosphate dehydrogenase (G3PDH; rabbit muscle, Merck, Darmstadt, Germany), which converts the DHAP product to glycerol 3-phosphate coupled to the oxidation of NADH to NAD^+^. Reaction mixtures (200 µL) contained 0.1 M HEPES–NaOH (pH 7.8, 30 °C), 0.7 mM NADH, 5 mM FBP and 2 U G3PDH. FBP aldolase (from *Saccharomyces cervisiae*; Merck, Darsmstadt, Germany) served as the positive control. Reactions were initiated by addition of enzyme, and all measurements were performed in triplicate. Negative controls omitted either enzyme or substrate. To obtain the kinetic parameters (Table 2), a classical Michaelis-Menten rate equation was fit to the complete dataset using NonlinearModelFit function in Wolfram Mathematica v14.2. EDA activity in the reverse (condensation) direction—forming KDPG from pyruvate and D-GAP was quantified using the colorimetric thio-barbituric acid (TBA) assay as described (46). Reactions (1 mL) contained each 20 mM pyruvate and 10 mM D-GAP in 0.1 M HEPES–NaOH (pH 7.8, 30 °C) with 15 µg EDA or 3 µg *Saci*KD(P)GA, *Saci*KD(P)GA was assayed at 70°C. At two-minute intervals, 50 µL aliquots were quenched with 5 µL of 12% trichloroacetic acid. Absorbance of the TBA–KDPG adduct was measured at 549 nm (ε = 67.8 mM⁻¹ cm⁻¹), and *Saci*KD(P)GA served as a positive control. EDA activity in reverse direction – forming OAA was assayed using malate dehydrogenase (MDH, *S. cervisiae*, Merck, Darmstadt, Germany) that forms malate from OAA by oxidation of NADPH to NADP^+^. EDA activity in reverse direction – forming KHG from pyruvate and glyoxylate was assayed by measuring the pyruvate consumption in a discontinuous assay. Pyruvate (5 mM) and glyoxylate (5 mM) were incubated with Eda (5 µg) in 1 mL of 0.1 M HEPES–NaOH buffer (pH 7.8) at 30 °C. At defined time points (0, 2, 4, 6, 8, 10, 20, and 30 min), 50 µL aliquots were withdrawn and quenched by the addition of 5 µL of 12% (v/v) TCA. Samples were immediately placed on ice and centrifuged at 8,130 × g for 10 min to remove precipitated protein. Supernatants (50 µL) were diluted into 950 µL of 0.1 M HEPES–NaOH (pH 7.8). Pyruvate consumption was quantified by a lactate dehydrogenase (LDH)–coupled assay using 0.7 mM NADH and 1 U of LDH (rabbit muscle; Merck, Darmstadt, Germany). The decrease in NADH absorbance reflected the conversion of pyruvate to lactate by LDH and was used to compare pyruvate levels across time points. It was ensured that auxiliary enzymes were not rate limiting. One unit (1 U) of enzyme activity is defined as 1 µmol substrate consumed or product formed per minute.

To test whether EDA exhibits activity toward 2-keto-3-deoxygluconate (KDG), an additional continuous coupled assay was employed. Reaction mixtures (200 µL final volume) contained 0.1 M HEPES–NaOH (pH 7.8, at 30 °C), 20 mM gluconate, 20 µg *Sulfolobus solfataricus* DHAD (*Sso*DHAD; SSO3107), 10 mM MgCl₂, and 1 mM DTT. The mixtures were incubated with 20 µg of either *Sso*DHAD at 50 °C or Slr0452 at 30 °C for 2 h. Reactions were initiated by the addition of EDA or *Sulfolobus acidocaldarius* KD(P)GA (*Saci*KD(P)GA; Saci_0225) as positive control, which converts the reaction product 2-keto-3-deoxygluconate (KDG) into pyruvate and glyceraldehyde. Pyruvate formation was detected as described above for the KDPG cleavage assay. All assays were performed in triplicate. Negative control reactions were performed in the absence of either the enzyme or the substrate.

## *In vitro* EDD enzyme assay

EDD catalyzes the dehydration of 6PG to form KDPG in the classical Entner–Doudoroff pathway. Enzyme activity was monitored using a continuous, coupled spectrophotometric assay in which the KDPG product was subsequently cleaved by excess *Cc*EDA to yield pyruvate and GAP. Pyruvate formation was detected as described above for the KDPG cleavage assay. NADH concentrations were quantified against a standard calibration curve (0–0.7 mM). Reaction mixtures (200 µL final volume) contained 0.1 M HEPES–NaOH (pH 7.8, 30 °C), 0.7 mM NADH, 10 mM MgCl₂, 5 mM 6PG, 5 µg *CcEDA* (recombinantly produced and characterized in this study, 150 U mg⁻¹), 2 U LDH, and the respective amounts of Slr0452 (20 µg) as the test sample and *Cc*EDD (5 µg) as positive control (47). To test whether Slr0452 exhibits activity toward gluconate, as reported for the bifunctional EDD/DHAD from *Sulfolobus solfactorius* (19), an additional continuous coupled assay was employed. Reaction mixtures (200 µL final volume) contained 0.1 M HEPES–NaOH (pH 7.8, at 30 °C), 0.7 mM NADH, 10 mM MgCl₂, and 1 mM DTT. The mixtures were incubated with 20 µg of either *S. solfataricus* DHAD (*Sso*DHAD) at 50 °C or Slr0452 at 30 °C. Reactions were initiated by the addition of *Sulfolobus acidocaldarius* KD(P)GA aldolase, which converts the reaction product 2-keto-3-deoxygluconate (KDG) into pyruvate and glyceraldehyde (20). Pyruvate formation was detected as described above for the KDPG cleavage assay. All assays were performed at least in triplicate. Negative controls omitted either the enzyme or substrate. It was ensured that auxiliary enzymes were not rate limiting.

## *In vitro* DHAD enzyme assay

Dihydroxyacid dehydratase (DHAD; *ilvD*) catalyzes a key step in the biosynthesis of branched-chain amino acids (BCAAs: leucine, isoleucine, and valine). The activities of Slr0452 and *S. elongatus* DHAD (*Se*DHAD) were assayed using the physiological substrate 2R-dihydroxyisovalerate (DHIV; Merck, Darmstadt, Germany), which is dehydrated to yield 2-ketoisovalerate (KIV). Product formation was quantified using a discontinuous colorimetric assay as described previously (17).

Reaction mixtures (100 µL) contained 50 mM Tris–HCl (pH 7.8), 1 mM dithiothreitol (DTT), 10 mM MgCl₂, and 1 mM 2R-DHIV. Reactions were initiated by the addition of 0.25 µM enzyme, incubated aerobically at room temperature for 30 min, and terminated with 25 µL of 2 M HCl. The resulting KIV was derivatized by addition of 50 µL of saturated 2,4-dinitrophenylhydrazine (DNPH; Häberle, Germany) in 2 M HCl and incubated for 30 min at room temperature. Subsequently, 25 µL of 10 M NaOH were added, samples were mixed, and precipitates were removed by centrifugation (5,000 × *g*, 10 min). Supernatants (100 µL) were transferred to clear 96-well microplates, and absorbance of the KIV-DNPH adduct was measured at 540 nm. Quantification was based on a KIV-DNPH calibration curve (0–1 mM), which exhibited linear response across this range. All enzyme assays were performed in triplicate.

## Bioinformatic analysis

Crystal structures were obtained from the Protein Data Bank (PDB), and structural models were retrieved from the AlphaFold Protein Structure Database (version 3.0) (26). Structural analyses and comparative assessments were performed using UCSF ChimeraX (48). Amino acid sequences for alignment were retrieved from UniProt, and multiple sequence alignments were generated using the EMBL-EBI Job Dispatcher framework (28), followed by manual refinement and editing in BioEdit.

## Acknowledgement

This study was supported by grants from and the German Science Foundation (DFG Gu1522/2-2, Gu1522/5-1, SI 642/14-1, SI 642/14-2, FOR2816, GRK2749/1). We acknowledge financial assistance from the DSI/NRF in South Africa (grant NRF-SARCHI 82813 to J. L. S.). We thank Markus Brand, Frauke Caliebe, Friederike Hörsch, Carolin Kremer and Maria Patti for engaged discussions.

## Authoŕs contributions

RO, MT, RF, AM, MB, and CP, performed the experiments; MB, CB, JLS, BS, KG supervised, RO, MT, RF, AM, MB, CP, CB, JLS, MH, BS, and KG analyzed data; RO, AM, and KG wrote the original draft; all authors wrote, reviewed, and edited; BS, and KG conceptualized the study; BS and KG acquired funding.

## Declaration of generative AI and AI-assisted technologies in the manuscript preparation process

During the preparation of this work the author(s) used ChatGPT (Open AI 2025), ChatGPT Team (5.1) and DeepL in order to optimize language. After using this tool/service, the authors reviewed and edited the content as needed and take full responsibility for the content of the published article.

## Conflict of interest

The authors declare to have no conflict of interest regarding the contents of this article.

## Data Availability

All study data are included in the article and SI Appendix. In addition, the kinetic data and model simulations are available as Excel files and Mathematica notebooks on the FAIRDOMHub (https://fairdomhub.org/investigations/775) and will be made public upon publication of the manuscript.

## Notes

### Competing Interest Statement

The authors have declared no competing interest.

### Summary of Updates

The manuscript was submitted to eLife and peer reviewed. The revised version was modified based on the recommendations made by the reviewers according to the responses of the authors. The public reviews and recommendations including the responses of the authors will be publicly available at eLife.

