## Supplementary Information for "Story about honest mistakes: The cyanobacterium *Synechocystis* has a promiscuous Entner-Doudoroff (ED) aldolase but no functional ED pathway"

**Figure S1**: Purification of recombinant Slr0452 from *Synechocystis* sp. PCC 6803 after expression in *E. coli*.
**Figure S2**: Purification of recombinant EDAK147G, EDA from *Caulobacter crescentus* (*Cc*EDA), KD(P)G aldolase from *Sulfolobus acidocaldarius* (*Saci*KD(P)GA), DHAD from *Synechococcus elongatus* (*Se*DHAD), and bifunctional dihydroxy acid dehydratase (DHAD) with gluconate dehydratase (GAD) activity from *Sulfolobus solfataricus* (*Sso*DHAD) expressed in *E. coli*.
**Figure S3**: (A) Substrate specificity of Slr0452 from *Synechocystis* sp. PCC 6803 and DHAD from *Synechococcus elongatus* (*Se*DHAD). (B) Enzymatic activity of Slr0452 and *Caulobacter crescentus* EDD *(Cc*EDD*)* with 6-phosphogluconate (6PG) as substrate.
**Figure S4**: Slr0452 and EDA (Sll0107) activities with gluconate and 2-keto-3-deoxygluconate (KDG) as substrates.
**Figure S5**: Sll1709 gene fragments synthesized by Genescript:
**Figure S6**: Verification of mutant overexpressing putative GDH1 (Sll1709) in *Synechocystis*
**Figure S7**: Cloning strategy and sequences for overexpression of putative GDHs (NEO87985.1and WP_238987112 and) and GKs (NEO86388.1 and WP_023068790.1) from *Spirulina sp. SIO3F2* and *Lyngbya aestuarii* and codon optimized sequence of EDA (Sll0107).
**Figure S8**: Purification of the recombinant GDH homologues from *Spirulina* sp. SIO3F2 (NEO87985.1) and *Lyngbya aestuarii* (WP_238987112) after expression in *E.coli*.
**Figure S9**: Purification of the GK from Spirulina sp. SIO3F2 and Lyngbya aestuarii after recombinant expression in *E. coli*.
**Figure S10**: Purification of the recombinant extended version of Sll0107 (EDA) from *Synechocystis* sp. PCC 6803 after expression in *E. coli*.
**Figure S11:** Native molecular mass of Sll0107 (EDA) from *Synechocystis*.
**Figure S12**: Enzymatic activity of *Synechocystis* EDA in the condensation (reverse) direction, determined by the thiobarbituric acid (TBA) assay.
**Table S1**: Result of blast analyses with 261 cyanobacteria next to *Synechocystis* based on a gluconate kinase (GK) of *Escherichia coli* and two glucose dehydrogenases (GDHs) as baits
**Table S2**: List of *Synechocystis* strains used in this study
**Table S3**: List of used primers for *Synechocystis* including primers for mutagenesis of extended codon optimized EDA

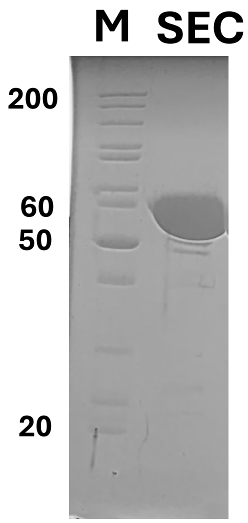


**Figure S1. Purification of recombinant Slr0452 from *Synechocystis* sp. PCC 6803 after expression in *E. coli*.** Purified protein (2 µg) following IMAC and size-exclusion chromatography was analyzed by SDS–PAGE and visualized with Coomassie Brilliant Blue. M, molecular weight marker (PageRuler™ Unstained Protein Ladder, Thermo Fisher Scientific). The apparent molecular mass (~60 kDa) is consistent with the calculated mass of 56 kDa.

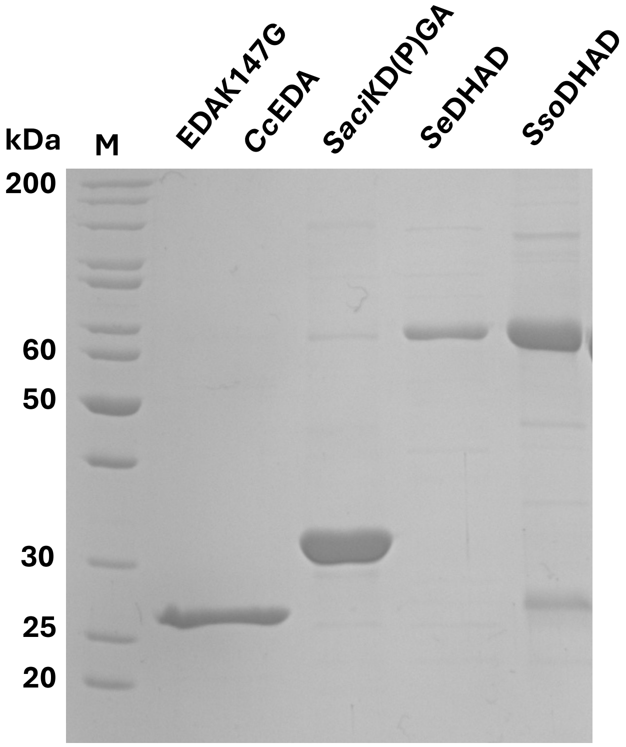


**Figure S2: Purification of recombinant EDAK147G, EDA from *Caulobacter crescentus* (*Cc*EDA), KD(P)G aldolase from *Sulfolobus acidocaldarius* (*Saci*KD(P)GA), DHAD from *Synechococcus elongatus* (*Se*DHAD), bifunctional dihydroxy acid dehydratase (DHAD) with gluconate dehydratase (GAD) activity from *Sulfolobus solfataricus* (*Sso*DHAD) after expression in *E. coli*.** Purified protein (2 µg) following IMAC was analyzed by SDS-PAGE and visualized with Coomassie Brilliant Blue. M, molecular weight marker (PageRuler™ Unstained Protein Ladder, Thermo Fisher Scientific). The apparent molecular mass of EDAk147G (25 kDa), *Cc*EDA (26 kDa), *Saci*KD(P)GA (32 kDa), *Se*DHAD (65 kDa), *Sso*DHAD (60 kDa) is consistent with calculated molecular mass of 23kDa, 24 kDa, 32.5 kDa, 66 kDa and 60 kDa, respectively.

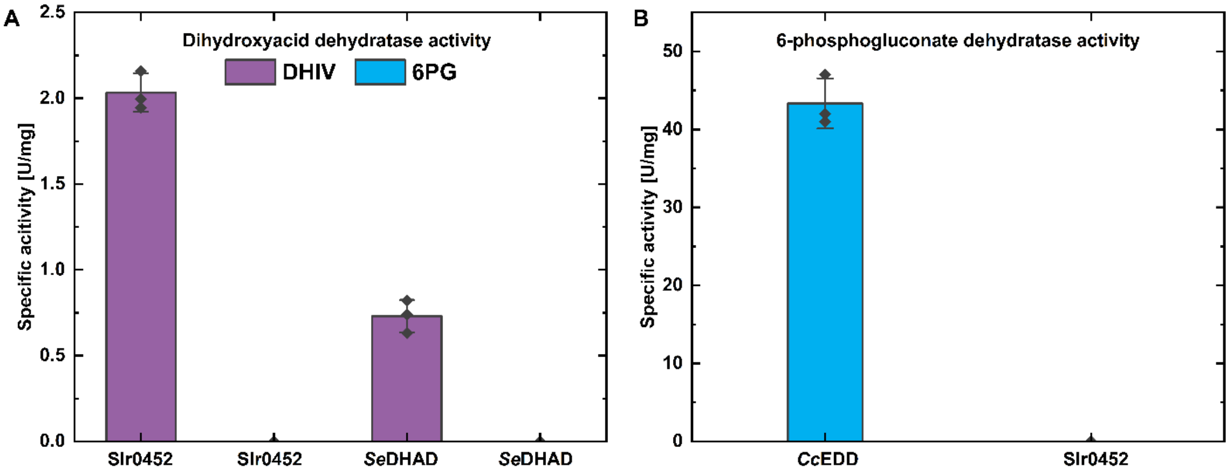


**Figure S3. (A) Substrate specificity of Slr0452 from *Synechocystis* sp. PCC 6803 and DHAD from *Synechococcus elongatus* (*Se*DHAD).** Enzyme activity was measured using 1 mM 2R-dihydroxyisovalerate (DHIV) and 5 mM 6-phosphogluconate (6PG) in the presence of 10 mM $MgCl_2$ and 1 mM DTT using a discontinuous DNPH assay (1). KIV formation was quantified using a calibration curve ranging from 0 to 1 mM. Both DHAD enzymes were specific for DHIV and did not act on 6PG. Data are presented as mean ± SD from three technical replicates (n = 3). **(B) Enzymatic activity of Slr0452 and *Caulobacter crescentus* EDD (*Cc*EDD) with 6-phosphogluconate (6PG) as substrate.** Assays were performed to test EDD activity. Activity was measured continuously using 5 mM 6PG as substrate in the presence of 10 mM $MgCl_2$ and 1 mM DTT. *Cc*EDD served as a positive control for 6PG dehydration. Data are presented as mean ± SD from three technical replicates (n = 3). The biochemical characterization of Slr0452 confirms that this enzyme is exclusively a DHAD and not an EDD as supposed by Evans et. al. 2024 (2)

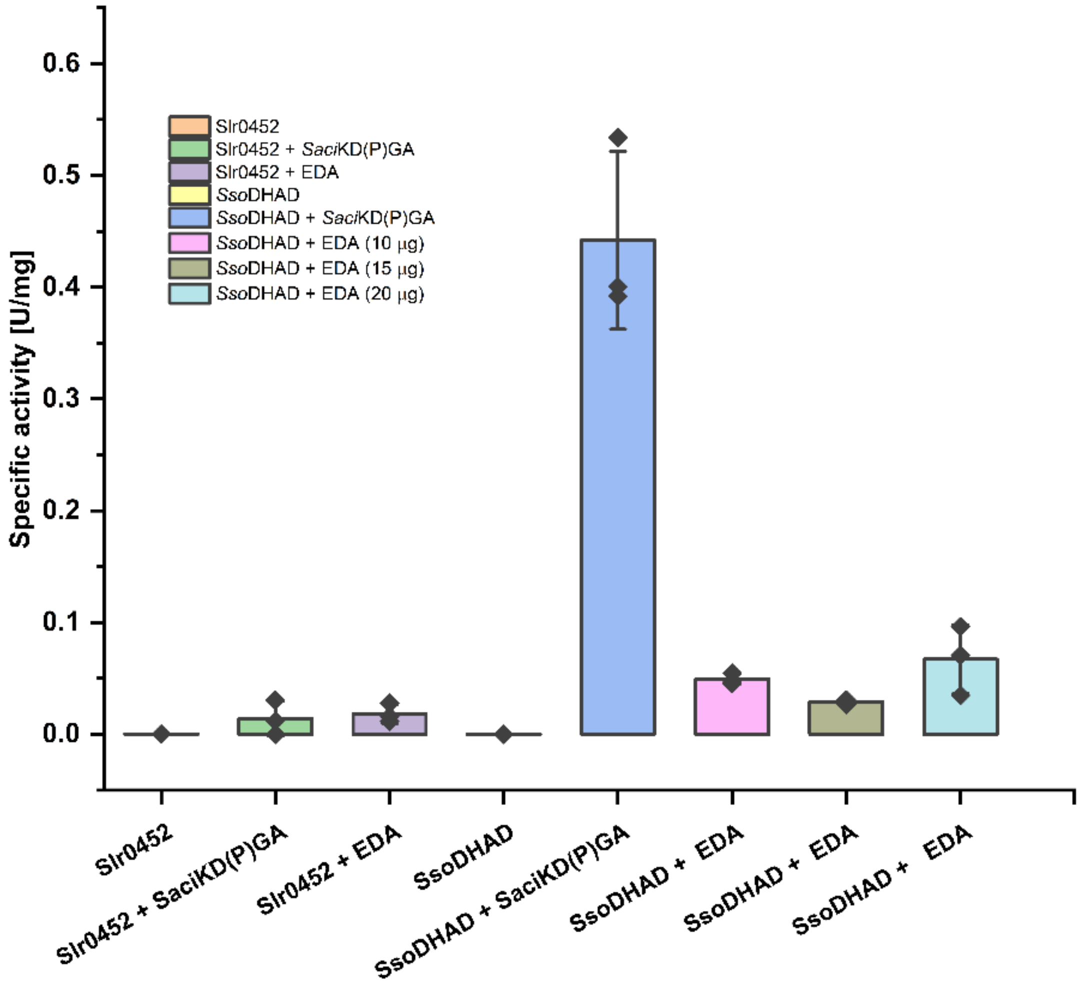


**Figure S4. Slr0452 and EDA (Sll0107) activities with gluconate and 2-keto-3-deoxygluconate (KDG) as substrates.** To evaluate the possibility of a modified ED pathway in *Synechocystis*, Slr0452 and EDA activities were assayed using gluconate and KDG as substrates, respectively. *Sso*DHAD and *Saci*KD(P)GA, previously shown to catalyze reactions with gluconate and KDG, respectively, were used as positive controls (3, 4). Gluconate (20 mM) was incubated with Slr0452 (20 µg) at 30 °C or with *Sso*DHAD (20 µg) at 50 °C for 2 h to allow conversion to KDG. Subsequently, either EDA was added to initiate the reaction at 30 °C, or *Saci*KD(P)GA (10 µg) was added to initiate the reaction at 50 °C. Pyruvate formation was detected as described for the KDPG cleavage assay. Increasing amounts of EDA (10–20 µg) were tested to confirm that EDA does not catalyze KDG as substrate even at higher enzyme concentrations.

**(A) GDH1up-BamHI-XhoI-NdeI-GDH1 (498 bp):**

GACTACTATAGGGCGAATTGGGTACGCTTGGTATTACCCCGATCCCAAACCCAGGGCGGAAAATATTA
AGGGCTACATCGCTTTTTGGCGGGGTGTAAAGGTTGAAGCCTGAATTTTTTTGTAATTATGGGCAATGA
TTTCGACTACATCGCCATAAAAAATGGTAATTTTGTTTTATTGTATCTCCACCCGACTAAGTTTGTAATC
GCCCAAGGCTCAGGAATTTAACTAACTACGGGATCCCTCGAGCATATGGAATTAACTGGAAAAACAGC
CCTGGTCACCGGGGGAGGCATAAGGGTTGGTCGGGCCCTAGTTATGGCCCTGGCGGAAGCCGGTTGTA
ACGTCTTTATTCACTACGGTAATTCCCAGACCGGAGCCCTAGAGGTTCAGCAAGAAGTGGAAGCATTG
GGTAGGAAAGCTTTTACCTACTCCGCCGATTTGAGCGATGCCCAGGCCACCCAAGGCTCGAGAGCTCC
AGCTTTTGTTCCCTTTAGT

**(B) BamHI-EcoRV-petEmod-HT-TEV-Start (494 bp):**

GACTACTATAGGGCGAATTGGGTACCATATGGAGCTCGGATCCGATATCCTAGCATAACCCCTTGGGGCCTCTAAAC
GGGTCTTGAGGGGTTTTTTGCGGCGATCGCCAAAAACAAAGAAAATTCAGCAATTACCGTGGGTAGCAAAAAATCCC
CATCTAAAGTTCAGTAAATATAGCTAGAACAACCAAGCATTTTCGGCAAAGTACTATTCAGATAGAACGAGAAATGAGCT
TGTTCTATCCGCCCGGGGCTGAGGCTGTATAATCTACGACGGGCTGTCAAACATTGTGATACCATGGGCAGAAGAAA
GGAAAAACGTCCCTGATCGCCTTTTTGGGCACGGAGTAGGGCGTTACCCCGGCCCGTTCAACCACAAGTCCCTAT
AGATACAGGAGATTAATAATATGCATCACCATCACCATCACCCCATCGGAGCTAGCGAAAACCTGTATTTTCAGGGCC
ATATGAGCTCCAGCTTTTGTTCCCTTTAGT

**Figure S5: Gene fragments synthesized by Genescript**: GDH1 *sll1709* (A) and modified petE promoter (B). Restriction sites and the 20 bp sequences that overlap with the pBluescript SK(+) vector are highlighted.

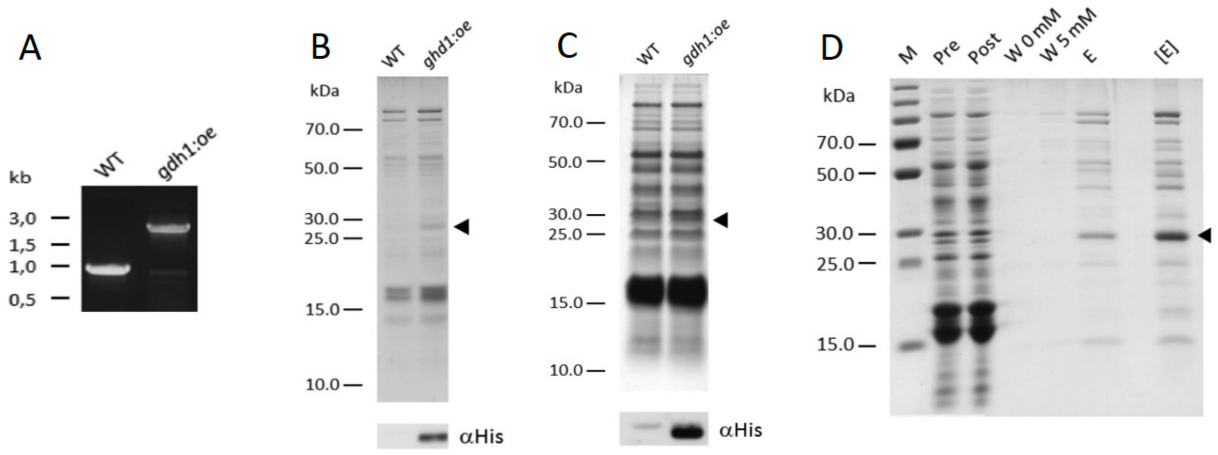


**Figure S6**: **Verification of mutant overexpressing putative GDH1 (Sll1709) in *Synechocystis*** (A) PCR showing segregation of gdh1:oe (*sll1709:oe)*. (B) SDS PAGE analysis followed by immunoblotting of soluble extracts confirmed expression and stable accumulation of putative GDH1 in gdh1:oe (*sll1709*:oe). (C) SDS PAGE and immunoblotting of small-scale His-tag purification of putative GDH1. (D) SDS PAGE analysis of the His-tagged putative GDH1 purification followed by immunoblotting with a His-tagged specific antibody. Soluble extracts before (Pre) and after (Post) incubation with Talon Cobalt resin, wash fraction samples with 0 and 5 mM imidazole, the eluted (7.5 µl of 10 mL) and concentrated eluted (7.5 µl of 800 µl @ ~0.29 mg/mL) fractions were loaded on the gel.

>WP_238987112 glucose 1-dehydrogenase [Lyngbya aestuarii]
MKGLKGKNVLITGATSGIGQAIAVRFAQEGANVAINYRGNPEKAEDTEEMIEQVCNQIRGCGGQDVLVEGDVSNEQDIIRMCEEVVEKLGSLDILINNA
GMQIAEPSDQVKTENFDQVISVNLRGAFLCSREAIKQFLKQNNGGVILNVSSVHEIIPRPEYVSYSMSKGGMGNLTKTLALEYARHGIRVNSIAPGATAT
PINSWTEDEDKRKEVEQHIPMGRVGTSEEMAAITAFLASDDAAYITGQTLYADGGLTLYPDFMKAWSAGE

codon optimized sequence (873 bp):
GTTTAACTTTAAGAAGGAGATATACATATGAAAGGCCTGAAAGGCAAAAACGTGCTGATTACCGGCGCGACGAGCGGCATTGGCCAAGCGATTGCG
GTGCGCTTTGCGCAAGAAGGCGCGAACGTGGCGATTAACTATCGCGGCAACCCGGAAAAAGCGGAAGATACCGAAGAAATGATTGAACAAGTGT
GCAATCAGATTCGCGGCTGCGGCGGCCAAGATGTGCTGGTGGAAGGCGATGTGAGCAACGAACAAGATATTATTCGCATGTGCGAAGAAGTGGTG
GAAAAACTGGGCAGCCTGGATATTCTGATTAACAACGCGGGCATGCAGATTGCGGAACCGAGCGATCAAGTGAAAACCGAAAACTTTGATCAAGTG
ATTAGCGTGAACCTGCGCGGCGCGTTTCTGTGCAGCCGCGAAGCGATTAAACAGTTTCTGAAACAGAACAACGGCGGCGTGATTCTGAACGTGAG
CAGCGTGCATGAAATTATTCCGCGCCCGGAATATGTGAGCTATAGCATGAGCAAAGGCGGCATGGGCAACCTGACCAAAACCCTGGCGCTGGAAT
ATGCGCGCCATGGCATTCGCGTGAACAGCATTGCGCCGGGCGCGACCGCGACCCCGATTAACAGCTGGACCGAAGATGAAGATAAACGCAAAG
AAGTGGAACAGCATATTCCGATGGGCCGCGTGGGCACGAGCGAAGAAATGGCGGCGATTACCGCGTTTCTGGCGAGCGATGATGCGGCGTATAT
TACCGGTCAGACCCTGTATGCGGATGGCGGCCTGACCCTGTATCCGGATTTTATGAAAGCGTGGAGCGCGGGCGAACCCATCGGAGGATCCGAA
AACCTGTATTTTCAGGGCGGC
NdeI(CATATG)target, Linker, BamHI (GGATCC),TEV cleavage siteGSTHis-tag
GATCTCGATCCCGCGAAATTAATACGACTCACTATAGGGAGACCACAACGGTTTCCCTCTAGAAATAATTTTGTTTAACTTTAAGAAGGAGATATACATA
TGAAAGGCCTGAAAGGCAAAAACGTGCTGATTACCGGCGCGACGAGCGGCATTGGCCAAGCGATTGCGGTGCGCTTTGCGCAAGAAGGCGCG
AACGTGGCGATTAACTATCGCGGCAACCCGGAAAAAGCGGAAGATACCGAAGAAATGATTGAACAAGTGTGCAATCAGATTCGCGGCTGCGGCG
GCCAAGATGTGCTGGTGGAAGGCGATGTGAGCAACGAACAAGATATTATTCGCATGTGCGAAGAAGTGGTGGAAAAACTGGGCAGCCTGGATATTC
TGATTAACAACGCGGGCATGCAGATTGCGGAACCGAGCGATCAAGTGAAAACCGAAAACTTTGATCAAGTGATTAGCGTGAACCTGCGCGGCGCG
TTTCTGTGCAGCCGCGAAGCGATTAAACAGTTTCTGAAACAGAACAACGGCGGCGTGATTCTGAACGTGAGCAGCGTGCATGAAATTATTCCGCGC
CCGGAATATGTGAGCTATAGCATGAGCAAAGGCGGCATGGGCAACCTGACCAAAACCCTGGCGCTGGAATATGCGCGCCATGGCATTCGCGTGA
ACAGCATTGCGCCGGGCGCGACCGCGACCCCGATTAACAGCTGGACCGAAGATGAAGATAAACGCAAAGAAGTGGAACAGCATATTCCGATGG
GCCGCGTGGGCACGAGCGAAGAAATGGCGGCGATTACCGCGTTTCTGGCGAGCGATGATGCGGCGTATATTACCGGTCAGACCCTGTATGCGGA
TGGCGGCCTGACCCTGTATCCGGATTTTATGAAAGCGTGGAGCGCGGGCGAACCCATCGGAGGATCCGAAAACCTGTATTTTCAGGGCGGCGCT
AGCATGTCCCCTATACTAGGTTATTGGAAAATTAAGGGCCTTGTGCAACCCACTCGACTTCTTTTGGAATATCTTGAAGAAAAATATGAAGAGCATTTGT
ATGAGCGCGATGAAGGTGATAAATGGCGAAACAAAAAGTTTGAATTGGGTTTGGAGTTTCCCAATCTTCCTTATTATATTGATGGTGATGTTAAATTAACA
CAGTCTATGGCCATCATACGTTATATAGCTGACAAGCACAACATGTTGGGTGGTTGTCCAAAAGAGCGTGCAGAGATTTCAATGCTTGAAGGAGCGGT
TTTGGATATTAGATACGGTGTTTCGAGAATTGCATATAGTAAAGACTTTGAAACTCTCAAAGTTGATTTTCTTAGCAAGCTACCTGAAATGCTGAAAATGTT
CGAAGATCGTTTATGTCATAAAACATATTTAAATGGTGATCATGTAACCCATCCTGACTTCATGTTGTATGACGCTCTTGATGTTGTTTTATACATGGACCC
AATGTGCCTGGATGCGTTCCCAAAATTAGTTTGTTTTAAAAAACGTATTGAAGCTATCCCACAAATTGATAAGTACTTGAAATCCAGCAAGTATATAGCAT
GGCCTTTGCAGGGCTGGCAAGCCACGTTTGGTGGTGGCGACCATCCTCCAAAATCGGATCATCACCATCACCATCACTAGGAATTCGAAGCTTGAT
CCGGCTGCTAACAAAGCCCGAAAGGAAGCTGAGTTGGCTGCTGCCACCGCTGAGCAATAACTAGCATAACCCCTTGGGGCCTCTAAACGGGTCT
TGAGGGGTTTTTTGCTGAAAGGAGGAACTATATCCGGATCTGGCGTAATAGCGAAGAGGCCCGCACCGATCGCCCTTCCCAACAGTTGCGCAGCC
TGAATGGCGAATGGGACGCGCCCTGTAGCGGCGCATTAAGCGCGGCGGGTGTGGTGGTTACGCGCAGCGTGACCGCTACACTTGCCAGCGCC
CTAGCGCCCGCTCCTTTCGCTTTCTTCCCTTCCTTTCTCGCCACGTTCGCCGGCTTTCCCCGTCAAGCTCTAAATCGGGGGCTCCCTTTAGGGTTC
CGATTTAGTGCTTTACGGCACCTCGACCCCAAAAAACTTGATTAGGGTGATGGTTCACGTAGTGGGCCATCGCCCTGATAGACGGTTTTTCGCCCTT
TGACGTTGGAGTCCACGTTCTTTAATAGTGGACTCTTGTTCCAAACTGGAACAACACTCAACCCTATCGCGGTCTATTCTTTTGATTTATAAGGGATTTTG
CCGATTTCGGCCTATTGGTTAAAAAATGAGCTGATTTAACAAATATTTAACGCGAATTTTAACAAAATATTAACGCTTACAATTTAGGTGGCACTTTTCGG
GGAAATGTGCGCGGAACCCCTATTTGTTTATTTTTCTAAATACATTCAAATATGTATCCGCTCATGAGACAATAACCCTGATAAATGCTTCAATAATATTGA
AAAAGGAAGAGTATGAGTATTCAACATTTCCGTGTCGCCCTTATTCCCTTTTTTGCGGCATTTTGCCTTCCTGTTTTTGCTCACCCAGAAACGCTGGTG
AAAGTAAAAGATGCTGAAGATCAGTTGGGTGCACGAGTGGGTTACATCGAACTGGATCTCAACAGCGGTAAGATCCTTGAGAGTTTTCGCCCCGAAG
AACGTTTTCCAATGATGAGCACTTTTAAAGTTCTGCTATGTGGCGCGGTATTATCCCGTATTGACGCCGGGCAAGAGCAACTCGGTCGCCCCATACA
CTATTCTCAGAATGACTTGGTTGAGTACTCACCAGTCACAGAAAAGCATCTTACGGATGGCATGACAGTAAGAGAATTATGCAGTGCTGCCATAACCAT
GAGTGATAACACTGCGGCCAACTTACTTCTGACAACGATCGGAGGACCGAAGGAGCTAACCGCTTTTTTGCACAACATGGGGGATCATGTAACTCG
CCTTGATCGTTGGGAACCGGAGCTGAATGAAGCCATACCAAACGACGAGAGTGACACCACGATGCCTGTAGCAATGCCAACAACGTTGCGCAAA
CTATTAACTGGCGAACTACTTACTCTAGCTTCCCGGCAACAATTAATAGACTGGATGGAGGCGGATAAAGTTGCAGGACCACTTCTGCGCTCGGCC
CTTCCGGCTGGCTGGTTTATTGCTGATAAATCTGGAGCCGGTGAGCGTGGGTCTCGCGGTATCATTGCAGCACTGGGGCCAGATGGTAAGCGCTC
CCGTATCGTAGTTATCTACACGACGGGGAGTCAGGCAACTATGGATGAACGAAATAGACAGATCGCTGAGATAGGTGCCTCACTGATTAAGCATTGG
TAACTGTCAGACCAAGTTTACTCATATATACTTTAGATTGATTTAAAACTTCATTTTTAATTTAAAAGGATCTAGGTGAAGATCCTTTTTGATAATCTCATGAC
CAAAATCCCTTAACGTGAGTTTTCGTTCCACTGAGCGTCAGACCCCGTAGAAAAGATCAAAGGATCTTCTTGAGATCCTTTTTTTCTGCGCGTAATCTG
CTGCTTGCAAACAAAAAAACCACCGCTACCAGCGGTGGTTTGTTTGCCGGATCAAGAGCTACCAACTCTTTTTCCGAAGGTAACTGGCTTCAGCAG
AGCGCAGATACCAAATACTGTCCTTCTAGTGTAGCCGTAGTTAGGCCACCACTTCAAGAACTCTGTAGCACCGCCTACATACCTCGCTCTGCTAATC
CTGTTACCAGTGGCTGCTGCCAGTGGCGATAAGTCGTGTCTTACCGGGTTGGACTCAAGACGATAGTTACCGGATAAGGCGCAGCGGTCGGGCTG
AACGGGGGGTTCGTGCACACAGCCCAGCTTGGAGCGAACGACCTACACCGAACTGAGATACCTACAGCGTGAGCTATGAGAAAGCGCCACGCT
TCCCGAAGGGAGAAAGGCGGACAGGTATCCGGTAAGCGGCAGGGTCGGAACAGGAGAGCGCACGAGGGAGCTTCCAGGGGGAAACGCCTGG
TATCTTTATAGTCCTGTCGGGTTTCGCCACCTCTGACTTGAGCGTCGATTTTTGTGATGCTCGTCAGGGGGGCGGAGCCTATGGAAAAACGCCAGCA
ACGCGGCCTTTTTACGGTTCCTGGGCTTTTGCTGGCCTTTTGCTCACATGTTCTTTCCTGCGTTATCCCCTGATTCTGTGGATAACCGTATTACCGCCT
TTGAGTGAGCTGATACCGCTCGCCGCAGCCGAACGACCGAGCGCAGCGAGTCAGTGAGCGAGGAAGCGGAAGAGCGCCCAATACGCAAACC
GCCTCTCCCCGCGCGTTGGCCGATTCATTAATGCAG
>NEO87985.1 glucose 1-dehydrogenase [Spirulina sp. SIO3F2]
MKLDGKVALVTGSSQGIGQAIAIRLAAAGAKVIINYRSHPEGAEATVNKINTAAGSNNQAYAIQADLGQVENVQRLVKESIEHFGQLDVLVNNAGIEKHA
AFWDVTEKDFDAVLNVNLKGVFFATQAATQHWMATQRPGKVVNISSVHEDLPFPNFAAYCASKGGLKMLTRNLSVELAPHNITINNVAPGAIETPINTK
LLNDPEKLGALLKNIPLGRLGQPGDVASLVEFLASDDAGYITGSTFFVDGGLTWNYQEQ

codon optimized sequence (840 bp):
GTTTAACTTTAAGAAGGAGATATACATATGAAACTGGATGGCAAAGTGGCGCTGGTGACCGGCAGCAGCCAAGGCATTGGCCAAGCGATTGCGATT
CGCCTGGCGGCGGCGGGCGCGAAAGTGATTATTAACTATCGCAGCCATCCGGAAGGCGCGGAAGCGACCGTGAACAAAATTAACACCGCGGC
GGGCAGCAACAACCAAGCGTATGCGATTCAAGCGGATCTGGGCCAAGTGGAAAACGTGCAGCGCCTGGTGAAAGAAAGCATTGAACATTTTGGTC
AGCTGGATGTGCTGGTGAACAACGCGGGCATTGAAAAACATGCGGCGTTTTGGGATGTGACCGAAAAAGATTTTGATGCGGTGCTGAACGTGAACC
TGAAAGGCGTGTTTTTTGCGACCCAAGCGGCGACGCAGCATTGGATGGCGACGCAGCGCCCGGGCAAAGTGGTGAACATTAGCAGCGTGCATG
AAGATCTGCCGTTTCCGAACTTTGCGGCGTATTGCGCGAGCAAAGGCGGCCTGAAAATGCTGACCCGCAACCTGAGCGTGGAACTGGCGCCGC
ATAACATTACCATTAACAACGTGGCGCCGGGCGCGATTGAAACCCCGATTAACACCAAACTGCTGAACGATCCGGAAAAACTGGGCGCGCTGCT
GAAAAACATTCCGCTGGGCCGCCTGGGTCAGCCGGGCGATGTGGCGAGCCTGGTGGAATTTCTGGCGAGCGATGATGCGGGCTATATTACCGG
CAGCACCTTTTTTGTGGATGGCGGCCTGACCTGGAACTATCAAGAACAGCCCATCGGAGGATCCGAAAACCTGTATTTTCAGGGCGGC
NdeI (CATATG)target,Linker,BamHI (GGATCC)TEV cleavage siteGSTHis-tag
GATCTCGATCCCGCGAAATTAATACGACTCACTATAGGGAGACCACAACGGTTTCCCTCTAGAAATAATTTTGTTTAACTTTAAGAAGGAGATATACATA
TGAAACTGGATGGCAAAGTGGCGCTGGTGACCGGCAGCAGCCAAGGCATTGGCCAAGCGATTGCGATTCGCCTGGCGGCGGCGGGCGCGAA
AGTGATTATTAACTATCGCAGCCATCCGGAAGGCGCGGAAGCGACCGTGAACAAAATTAACACCGCGGCGGGCAGCAACAACCAAGCGTATGCG
ATTCAAGCGGATCTGGGCCAAGTGGAAAACGTGCAGCGCCTGGTGAAAGAAAGCATTGAACATTTTGGTCAGCTGGATGTGCTGGTGAACAACGC
GGGCATTGAAAAACATGCGGCGTTTTGGGATGTGACCGAAAAAGATTTTGATGCGGTGCTGAACGTGAACCTGAAAGGCGTGTTTTTTGCGACCCAA
GCGGCGACGCAGCATTGGATGGCGACGCAGCGCCCGGGCAAAGTGGTGAACATTAGCAGCGTGCATGAAGATCTGCCGTTTCCGAACTTTGCG
GCGTATTGCGCGAGCAAAGGCGGCCTGAAAATGCTGACCCGCAACCTGAGCGTGGAACTGGCGCCGCATAACATTACCATTAACAACGTGGCG
CCGGGCGCGATTGAAACCCCGATTAACACCAAACTGCTGAACGATCCGGAAAAACTGGGCGCGCTGCTGAAAAACATTCCGCTGGGCCGCCTG
GGTCAGCCGGGCGATGTGGCGAGCCTGGTGGAATTTCTGGCGAGCGATGATGCGGGCTATATTACCGGCAGCACCTTTTTTGTGGATGGCGGCCT
GACCTGGAACTATCAAGAACAGCCCATCGGAGGATCCGAAAACCTGTATTTTCAGGGCGGCGCTAGCATGTCCCCTATACTAGGTTATTGGAAAATT
AAGGGCCTTGTGCAACCCACTCGACTTCTTTTGGAATATCTTGAAGAAAAATATGAAGAGCATTTGTATGAGCGCGATGAAGGTGATAAATGGCGAAA
CAAAAAGTTTGAATTGGGTTTGGAGTTTCCCAATCTTCCTTATTATATTGATGGTGATGTTAAATTAACACAGTCTATGGCCATCATACGTTATATAGCTGAC
AAGCACAACATGTTGGGTGGTTGTCCAAAAGAGCGTGCAGAGATTTCAATGCTTGAAGGAGCGGTTTTGGATATTAGATACGGTGTTTCGAGAATTGC
ATATAGTAAAGACTTTGAAACTCTCAAAGTTGATTTTCTTAGCAAGCTACCTGAAATGCTGAAAATGTTCGAAGATCGTTTATGTCATAAAACATATTTAAAT
GGTGATCATGTAACCCATCCTGACTTCATGTTGTATGACGCTCTTGATGTTGTTTTATACATGGACCCAATGTGCCTGGATGCGTTCCCAAAATTAGTTTG
TTTTAAAAAACGTATTGAAGCTATCCCACAAATTGATAAGTACTTGAAATCCAGCAAGTATATAGCATGGCCTTTGCAGGGCTGGCAAGCCACGTTTGG
TGGTGGCGACCATCCTCCAAAATCGGATCATCACCATCACCATCACTAGGAATTCGAAGCTTGATCCGGCTGCTAACAAAGCCCGAAAGGAAGCT
GAGTTGGCTGCTGCCACCGCTGAGCAATAACTAGCATAACCCCTTGGGGCCTCTAAACGGGTCTTGAGGGGTTTTTTGCTGAAAGGAGGAACTATA
TCCGGATCTGGCGTAATAGCGAAGAGGCCCGCACCGATCGCCCTTCCCAACAGTTGCGCAGCCTGAATGGCGAATGGGACGCGCCCTGTAGCG
GCGCATTAAGCGCGGCGGGTGTGGTGGTTACGCGCAGCGTGACCGCTACACTTGCCAGCGCCCTAGCGCCCGCTCCTTTCGCTTTCTTCCCTTC
CTTTCTCGCCACGTTCGCCGGCTTTCCCCGTCAAGCTCTAAATCGGGGGCTCCCTTTAGGGTTCCGATTTAGTGCTTTACGGCACCTCGACCCCAA
AAAACTTGATTAGGGTGATGGTTCACGTAGTGGGCCATCGCCCTGATAGACGGTTTTTCGCCCTTTGACGTTGGAGTCCACGTTCTTTAATAGTGGAC
TCTTGTTCCAAACTGGAACAACACTCAACCCTATCGCGGTCTATTCTTTTGATTTATAAGGGATTTTGCCGATTTCGGCCTATTGGTTAAAAAATGAGCT
GATTTAACAAATATTTAACGCGAATTTTAACAAAATATTAACGCTTACAATTTAGGTGGCACTTTTCGGGGAAATGTGCGCGGAACCCCTATTTGTTTATTT
TTCTAAATACATTCAAATATGTATCCGCTCATGAGACAATAACCCTGATAAATGCTTCAATAATATTGAAAAAGGAAGAGTATGAGTATTCAACATTTCCGT
GTCGCCCTTATTCCCTTTTTTGCGGCATTTTGCCTTCCTGTTTTTGCTCACCCAGAAACGCTGGTGAAAGTAAAAGATGCTGAAGATCAGTTGGGTGC
ACGAGTGGGTTACATCGAACTGGATCTCAACAGCGGTAAGATCCTTGAGAGTTTTCGCCCCGAAGAACGTTTTCCAATGATGAGCACTTTTAAAGTTC
TGCTATGTGGCGCGGTATTATCCCGTATTGACGCCGGGCAAGAGCAACTCGGTCGCCCCATACACTATTCTCAGAATGACTTGGTTGAGTACTCACC
AGTCACAGAAAAGCATCTTACGGATGGCATGACAGTAAGAGAATTATGCAGTGCTGCCATAACCATGAGTGATAACACTGCGGCCAACTTACTTCTGA
CAACGATCGGAGGACCGAAGGAGCTAACCGCTTTTTTGCACAACATGGGGGATCATGTAACTCGCCTTGATCGTTGGGAACCGGAGCTGAATGAA
GCCATACCAAACGACGAGAGTGACACCACGATGCCTGTAGCAATGCCAACAACGTTGCGCAAACTATTAACTGGCGAACTACTTACTCTAGCTTCC
CGGCAACAATTAATAGACTGGATGGAGGCGGATAAAGTTGCAGGACCACTTCTGCGCTCGGCCCTTCCGGCTGGCTGGTTTATTGCTGATAAATCT
GGAGCCGGTGAGCGTGGGTCTCGCGGTATCATTGCAGCACTGGGGCCAGATGGTAAGCGCTCCCGTATCGTAGTTATCTACACGACGGGGAGTC
AGGCAACTATGGATGAACGAAATAGACAGATCGCTGAGATAGGTGCCTCACTGATTAAGCATTGGTAACTGTCAGACCAAGTTTACTCATATATACTTTA
GATTGATTTAAAACTTCATTTTTAATTTAAAAGGATCTAGGTGAAGATCCTTTTTGATAATCTCATGACCAAAATCCCTTAACGTGAGTTTTCGTTCCACTGA
GCGTCAGACCCCGTAGAAAAGATCAAAGGATCTTCTTGAGATCCTTTTTTTCTGCGCGTAATCTGCTGCTTGCAAACAAAAAAACCACCGCTACCAG
CGGTGGTTTGTTTGCCGGATCAAGAGCTACCAACTCTTTTTCCGAAGGTAACTGGCTTCAGCAGAGCGCAGATACCAAATACTGTCCTTCTAGTGTAG
CCGTAGTTAGGCCACCACTTCAAGAACTCTGTAGCACCGCCTACATACCTCGCTCTGCTAATCCTGTTACCAGTGGCTGCTGCCAGTGGCGATAAG
TCGTGTCTTACCGGGTTGGACTCAAGACGATAGTTACCGGATAAGGCGCAGCGGTCGGGCTGAACGGGGGGTTCGTGCACACAGCCCAGCTTGG
AGCGAACGACCTACACCGAACTGAGATACCTACAGCGTGAGCTATGAGAAAGCGCCACGCTTCCCGAAGGGAGAAAGGCGGACAGGTATCCGG
TAAGCGGCAGGGTCGGAACAGGAGAGCGCACGAGGGAGCTTCCAGGGGGAAACGCCTGGTATCTTTATAGTCCTGTCGGGTTTCGCCACCTCTG
ACTTGAGCGTCGATTTTTGTGATGCTCGTCAGGGGGGCGGAGCCTATGGAAAAACGCCAGCAACGCGGCCTTTTTACGGTTCCTGGGCTTTTGCTG
GCCTTTTGCTCACATGTTCTTTCCTGCGTTATCCCCTGATTCTGTGGATAACCGTATTACCGCCTTTGAGTGAGCTGATACCGCTCGCCGCAGCCGAA
CGACCGAGCGCAGCGAGTCAGTGAGCGAGGAAGCGGAAGAGCGCCCAATACGCAAACCGCCTCTCCCCGCGCGTTGGCCGATTCATTAATGC
AG

WP_023068790.1 gluconokinase [Lyngbya aestuarii]
MIILVMGVSGSGKTTIGEKLAESLNFQFRDADEFHPDENIKKMANNIPLTDEDRQPWLERMQTTIDQWLQDNQNVVLTCSALKEKYRQILWRDTEKME
LVYLKGSFELIKDRMIKRHDHFMKADLLRSQFEDLEEPTGGIWVDISQTPSEIVQEIKQKLML

codon optimized sequence (873 bp):
GTTTAACTTTAAGAAGGAGATATACATATGATTATTCTGGTGATGGGCGTGAGCGGCAGCGGCAAAACCACCATTGGCGAAAAACTGGCGGAAAGC
CTGAACTTTCAGTTTCGCGATGCGGATGAATTTCATCCGGATGAAAACATTAAAAAAATGGCGAACAACATTCCGCTGACCGATGAAGATCGCCAGC
CGTGGCTGGAACGCATGCAGACCACCATTGATCAGTGGCTGCAGGATAACCAGAACGTGGTGCTGACCTGCAGCGCGCTGAAAGAAAAATATCG

CCAGATTCTGTGGCGCGATACCGAAAAAATGGAACTGGTGTATCTGAAAGGCAGCTTTGAACTGATTAAAGATCGCATGATTAAACGCCATGATCATTT
TATGAAAGCGGATCTGCTGCGCAGCCAGTTTGAAGATCTGGAAGAACCGACCGGCGGCATTTGGGTGGATATTAGCCAGACCCCGAGCGAAATTG
TGCAGGAAATTAAACAGAAACTGATGCTGCCCATCGGAGGATCCGAAAACCTGTATTTTCAGGGCGGC
NdeI(<u>CAT</u>ATG)target,Linker,BamHI (GGATCC)TEV cleavage siteGSTHis-tag
GATCTCGATCCCGCGAAATTAATACGACTCACTATAGGGAGACCACAACGGTTTCCCTCTAGAAATAATTTTGTTTAACTTTAAGAAGGAGATATA<u>CAT</u>A
TGATTATTCTGGTGATGGGCGTGAGCGGCAGCGGCAAAACCACCATTGGCGAAAAACTGGCGGAAAGCCTGAACTTTCAGTTTCGCGATGCGGAT
GAATTTCATCCGGATGAAAACATTAAAAAAATGGCGAACAACATTCCGCTGACCGATGAAGATCGCCAGCCGTGGCTGGAACGCATGCAGACCAC
CATTGATCAGTGGCTGCAGGATAACCAGAACGTGGTGCTGACCTGCAGCGCGCTGAAAGAAAAATATCGCCAGATTCTGTGGCGCGATACCGAAA
AAATGGAACTGGTGTATCTGAAAGGCAGCTTTGAACTGATTAAAGATCGCATGATTAAACGCCATGATCATTTTATGAAAGCGGATCTGCTGCGCAGC
CAGTTTGAAGATCTGGAAGAACCGACCGGCGGCATTTGGGTGGATATTAGCCAGACCCCGAGCGAAATTGTGCAGGAAATTAAACAGAAACTGATG
CTGCCCATCGGAGGATCCGAAAACCTGTATTTTCAGGGCGGCGCTAGCATGTCCCCTATACTAGGTTATTGGAAAATTAAGGGCCTTGTGCAACCCA
CTCGACTTCTTTTGGAATATCTTGAAGAAAAATATGAAGAGCATTTGTATGAGCGCGATGAAGGTGATAAATGGCGAAACAAAAAGTTTGAATTGGGTTT
GGAGTTTCCCAATCTTCCTTATTATATTGATGGTGATGTTAAATTAACACAGTCTATGGCCATCATACGTTATATAGCTGACAAGCACAACATGTTGGGTG
GTTGTCCAAAAGAGCGTGCAGAGATTTCAATGCTTGAAGGAGCGGTTTTGGATATTAGATACGGTGTTTCGAGAATTGCATATAGTAAAGACTTTGAAAC
TCTCAAAGTTGATTTTCTTAGCAAGCTACCTGAAATGCTGAAAATGTTCGAAGATCGTTTATGTCATAAAACATATTTAAATGGTGATCATGTAACCCATCC
TGACTTCATGTTGTATGACGCTCTTGATGTTGTTTTATACATGGACCCAATGTGCCTGGATGCGTTCCCAAAATTAGTTTGTTTTAAAAAACGTATTGAAGC
TATCCCACAAATTGATAAGTACTTGAAATCCAGCAAGTATATAGCATGGCCTTTGCAGGGCTGGCAAGCCACGTTTGGTGGTGGCGACCATCCTCCA
AAATCGGATCATCACCATCACCATCACTAGGAATTCGAAGCTTGATCCGGCTGCTAACAAAGCCCGAAAGGAAGCTGAGTTGGCTGCTGCCACCG
CTGAGCAATAACTAGCATAACCCCTTGGGGCCTCTAAACGGGTCTTGAGGGGTTTTTTGCTGAAAGGAGGAACTATATCCGGATCTGGCGTAATAGC
GAAGAGGCCCGCACCGATCGCCCTTCCCAACAGTTGCGCAGCCTGAATGGCGAATGGGACGCGCCCTGTAGCGGCGCATTAAGCGCGGCGG
GTGTGGTGGTTACGCGCAGCGTGACCGCTACACTTGCCAGCGCCCTAGCGCCCGCTCCTTTCGCTTTCTTCCCTTCCTTTCTCGCCACGTTCGCC
GGCTTTCCCCGTCAAGCTCTAAATCGGGGGCTCCCTTTAGGGTTCCGATTTAGTGCTTTACGGCACCTCGACCCCAAAAAACTTGATTAGGGTGATG
GTTCACGTAGTGGGCCATCGCCCTGATAGACGGTTTTTCGCCCTTTGACGTTGGAGTCCACGTTCTTTAATAGTGGACTCTTGTTCCAAACTGGAACA
ACACTCAACCCTATCGCGGTCTATTCTTTTGATTTATAAGGGATTTTGCCGATTTCGGCCTATTGGTTAAAAAATGAGCTGATTTAACAAATATTTAACGC
GAATTTTAACAAAATATTAACGCTTACAATTTAGGTGGCACTTTTCGGGGAAATGTGCGCGGAACCCCTATTTGTTTATTTTTCTAAATACATTCAAATATG
TATCCGCTCATGAGACAATAACCCTGATAAATGCTTCAATAATATTGAAAAAGGAAGAGTATGAGTATTCAACATTTCCGTGTCGCCCTTATTCCCTTTTTT
GCGGCATTTTGCCTTCCTGTTTTTGCTCACCCAGAAACGCTGGTGAAAGTAAAAGATGCTGAAGATCAGTTGGGTGCACGAGTGGGTTACATCGAAC
TGGATCTCAACAGCGGTAAGATCCTTGAGAGTTTTCGCCCCGAAGAACGTTTTCCAATGATGAGCACTTTTAAAGTTCTGCTATGTGGCGCGGTATTAT
CCCGTATTGACGCCGGGCAAGAGCAACTCGGTCGCCCCATACACTATTCTCAGAATGACTTGGTTGAGTACTCACCAGTCACAGAAAAGCATCTTA
CGGATGGCATGACAGTAAGAGAATTATGCAGTGCTGCCATAACCATGAGTGATAACACTGCGGCCAACTTACTTCTGACAACGATCGGAGGACCGA
AGGAGCTAACCGCTTTTTTGCACAACATGGGGGATCATGTAACTCGCCTTGATCGTTGGGAACCGGAGCTGAATGAAGCCATACCAAACGACGAGA
GTGACACCACGATGCCTGTAGCAATGCCAACAACGTTGCGCAAACTATTAACTGGCGAACTACTTACTCTAGCTTCCCGGCAACAATTAATAGACTG
GATGGAGGCGGATAAAGTTGCAGGACCACTTCTGCGCTCGGCCCTTCCGGCTGGCTGGTTTATTGCTGATAAATCTGGAGCCGGTGAGCGTGGGT
CTCGCGGTATCATTGCAGCACTGGGGCCAGATGGTAAGCGCTCCCGTATCGTAGTTATCTACACGACGGGGAGTCAGGCAACTATGGATGAACGA
AATAGACAGATCGCTGAGATAGGTGCCTCACTGATTAAGCATTGGTAACTGTCAGACCAAGTTTACTCATATATACTTTAGATTGATTTAAAACTTCATTTT
TAATTTAAAAGGATCTAGGTGAAGATCCTTTTTGATAATCTCATGACCAAAATCCCTTAACGTGAGTTTTCGTTCCACTGAGCGTCAGACCCCGTAGAAA
AGATCAAAGGATCTTCTTGAGATCCTTTTTTTCTGCGCGTAATCTGCTGCTTGCAAACAAAAAAACCACCGCTACCAGCGGTGGTTTGTTTGCCGGAT
CAAGAGCTACCAACTCTTTTTCCGAAGGTAACTGGCTTCAGCAGAGCGCAGATACCAAATACTGTCCTTCTAGTGTAGCCGTAGTTAGGCCACCACT
TCAAGAACTCTGTAGCACCGCCTACATACCTCGCTCTGCTAATCCTGTTACCAGTGGCTGCTGCCAGTGGCGATAAGTCGTGTCTTACCGGGTTGGA
CTCAAGACGATAGTTACCGGATAAGGCGCAGCGGTCGGGCTGAACGGGGGGTTCGTGCACACAGCCCAGCTTGGAGCGAACGACCTACACCG
AACTGAGATACCTACAGCGTGAGCTATGAGAAAGCGCCACGCTTCCCGAAGGGAGAAAGGCGGACAGGTATCCGGTAAGCGGCAGGGTCGGAA
CAGGAGAGCGCACGAGGGAGCTTCCAGGGGGAAACGCCTGGTATCTTTATAGTCCTGTCGGGTTTCGCCACCTCTGACTTGAGCGTCGATTTTTGT
GATGCTCGTCAGGGGGGCGGAGCCTATGGAAAAACGCCAGCAACGCGGCCTTTTTACGGTTCCTGGGCTTTTGCTGGCCTTTTGCTCACATGTTC
TTTCCTGCGTTATCCCCTGATTCTGTGGATAACCGTATTACCGCCTTTGAGTGAGCTGATACCGCTCGCCGCAGCCGAACGACCGAGCGCAGCGA
GTCAGTGAGCGAGGAAGCGGAAGAGCGCCCAATACGCAAACCGCCTCTCCCCGCGCGTTGGCCGATTCATTAATGCAG

>NEO86388.1 MAG: gluconokinase [Spirulina sp. SIO3F2]
MGVSGCGKSTVGQALADRLGWPFYDGDDFHPAANIEKMAQGIPLMDSDRFPWLESIHQHIAHCQQTGKAAVLACSALKHIYRQQLRGDLNNIFIVYL
QGGFDLIRARLQARHGHFMKADLLHSQFADLEEPQDAIAVDIDQPLNAMIATIVAGLKDCAIMP

<u>codon optimized sequence:</u>
GTTTAACTTTAAGAAGGAGATATACATATGGGCGTGAGCGGCTGCGGCAAAAGCACCGTGGGCCAGGCGCTGGCGGATCGCCTGGGCTGGCCG
TTTTATGATGGCGATGATTTTCATCCGGCGGCGAACATTGAAAAAATGGCGCAGGGCATTCCGCTGATGGATAGCGATCGCTTTCCGTGGCTGGAAA
GCATTCATCAGCATATTGCGCATTGCCAGCAGACCGGCAAAGCGGCGGTGCTGGCGTGCAGCGCGCTGAAACATATTTATCGCCAGCAGCTGCG
CGGCGATCTGAACAACATTTTTATTGTGTATCTGCAGGGCGGCTTTGATCTGATTCGCGCGCGCCTGCAGGCGCGCCATGGCCATTTTATGAAAGC
GGATCTGCTGCATAGCCAGTTTGCGGATCTGGAAGAACCGCAGGATGCGATTGCGGTGGATATTGATCAGCCGCTGAACGCGATGATTGCGACCAT
TGTGGCGGGCCTGAAAGATTGCGCGATTATGCCGCCCATCGGAGGATCCGAAAACCTGTATTTTCAGGGCGGC
NdeI(<u>CAT</u>ATG)target, Linker, BamHI(GGATCC)TEV cleavage siteGSTHis-tag
GATCTCGATCCCGCGAAATTAATACGACTCACTATAGGGAGACCACAACGGTTTCCCTCTAGAAATAATTTTGTTTAACTTTAAGAAGGAGATATA<u>CAT</u>A
TGGGCGTGAGCGGCTGCGGCAAAAGCACCGTGGGCCAGGCGCTGGCGGATCGCCTGGGCTGGCCGTTTTATGATGGCGATGATTTTCATCCGG
CGGCGAACATTGAAAAAATGGCGCAGGGCATTCCGCTGATGGATAGCGATCGCTTTCCGTGGCTGGAAAGCATTCATCAGCATATTGCGCATTGCC
AGCAGACCGGCAAAGCGGCGGTGCTGGCGTGCAGCGCGCTGAAACATATTTATCGCCAGCAGCTGCGCGGCGATCTGAACAACATTTTTATTGTG
TATCTGCAGGGCGGCTTTGATCTGATTCGCGCGCGCCTGCAGGCGCGCCATGGCCATTTTATGAAAGCGGATCTGCTGCATAGCCAGTTTGCGGA
TCTGGAAGAACCGCAGGATGCGATTGCGGTGGATATTGATCAGCCGCTGAACGCGATGATTGCGACCATTGTGGCGGGCCTGAAAGATTGCGCGA
TTATGCCGCCCATCGGAGGATCCGAAAACCTGTATTTTCAGGGCGGCGCTAGCATGTCCCCTATACTAGGTTATTGGAAAATTAAGGGCCTTGTGCA
ACCCACTCGACTTCTTTTGGAATATCTTGAAGAAAAATATGAAGAGCATTTGTATGAGCGCGATGAAGGTGATAAATGGCGAAACAAAAAGTTTGAATTG

GGTTTGGAGTTTCCCAATCTTCCTTATTATATTGATGGTGATGTTAAATTAACACAGTCTATGGCCATCATACGTTATATAGCTGACAAGCACAACATGTTG
GGTGGTTGTCCAAAAGAGCGTGCAGAGATTTCAATGCTTGAAGGAGCGGTTTTGGATATTAGATACGGTGTTTCGAGAATTGCATATAGTAAAGACTTT
GAAACTCTCAAAGTTGATTTTCTTAGCAAGCTACCTGAAATGCTGAAAATGTTCGAAGATCGTTTATGTCATAAAACATATTTAAATGGTGATCATGTAACC
CATCCTGACTTCATGTTGTATGACGCTCTTGATGTTGTTTTATACATGGACCCAATGTGCCTGGATGCGTTCCCAAAATTAGTTTGTTTTAAAAAACGTATT
GAAGCTATCCCACAAATTGATAAGTACTTGAAATCCAGCAAGTATATAGCATGGCCTTTGCAGGGCTGGCAAGCCACGTTTGGTGGTGGCGACCATC
CTCCAAAATCGGATCATCACCATCACCATCACTAGGAATTCGAAGCTTGATCCGGCTGCTAACAAAGCCCGAAAGGAAGCTGAGTTGGCTGCTGCC
ACCGCTGAGCAATAACTAGCATAACCCCTTGGGGCCTCTAAACGGGTCTTGAGGGGTTTTTTGCTGAAAGGAGGAACTATATCCGGATCTGGCGTAA
TAGCGAAGAGGCCCGCACCGATCGCCCTTCCCAACAGTTGCGCAGCCTGAATGGCGAATGGGACGCGCCCTGTAGCGGCGCATTAAGCGCGG
CGGGTGTGGTGGTTACGCGCAGCGTGACCGCTACACTTGCCAGCGCCCTAGCGCCCGCTCCTTTCGCTTTCTTCCCTTCCTTTCTCGCCACGTTC
GCCGGCTTTCCCCGTCAAGCTCTAAATCGGGGGCTCCCTTTAGGGTTCCGATTTAGTGCTTTACGGCACCTCGACCCCAAAAAACTTGATTAGGGT
GATGGTTCACGTAGTGGGCCATCGCCCTGATAGACGGTTTTTCGCCCTTTGACGTTGGAGTCCACGTTCTTTAATAGTGGACTCTTGTTCCAAACTGG
AACAACACTCAACCCTATCGCGGTCTATTCTTTTGATTTATAAGGGATTTTGCCGATTTCGGCCTATTGGTTAAAAAATGAGCTGATTTAACAAATATTTAA
CGCGAATTTTAACAAAATATTAACGCTTACAATTTAGGTGGCACTTTTCGGGGAAATGTGCGCGGAACCCCTATTTGTTTATTTTTCTAAATACATTCAAA
TATGTATCCGCTCATGAGACAATAACCCTGATAAATGCTTCAATAATATTGAAAAAGGAAGAGTATGAGTATTCAACATTTCCGTGTCGCCCTTATTCCCT
TTTTTGCGGCATTTTGCCTTCCTGTTTTTGCTCACCCAGAAACGCTGGTGAAAGTAAAAGATGCTGAAGATCAGTTGGGTGCACGAGTGGGTTACATC
GAACTGGATCTCAACAGCGGTAAGATCCTTGAGAGTTTTCGCCCCGAAGAACGTTTTCCAATGATGAGCACTTTTAAAGTTCTGCTATGTGGCGCGGT
ATTATCCCGTATTGACGCCGGGCAAGAGCAACTCGGTCGCCCCATACACTATTCTCAGAATGACTTGGTTGAGTACTCACCAGTCACAGAAAAGCAT
CTTACGGATGGCATGACAGTAAGAGAATTATGCAGTGCTGCCATAACCATGAGTGATAACACTGCGGCCAACTTACTTCTGACAACGATCGGAGGAC
CGAAGGAGCTAACCGCTTTTTTGCACAACATGGGGGATCATGTAACTCGCCTTGATCGTTGGGAACCGGAGCTGAATGAAGCCATACCAAACGAC
GAGAGTGACACCACGATGCCTGTAGCAATGCCAACAACGTTGCGCAAACTATTAACTGGCGAACTACTTACTCTAGCTTCCCGGCAACAATTAATAG
ACTGGATGGAGGCGGATAAAGTTGCAGGACCACTTCTGCGCTCGGCCCTTCCGGCTGGCTGGTTTATTGCTGATAAATCTGGAGCCGGTGAGCGT
GGGTCTCGCGGTATCATTGCAGCACTGGGGCCAGATGGTAAGCGCTCCCGTATCGTAGTTATCTACACGACGGGGAGTCAGGCAACTATGGATGA
ACGAAATAGACAGATCGCTGAGATAGGTGCCTCACTGATTAAGCATTGGTAACTGTCAGACCAAGTTTACTCATATATACTTTAGATTGATTTAAAACTTC
ATTTTTAATTTAAAAGGATCTAGGTGAAGATCCTTTTTGATAATCTCATGACCAAAATCCCTTAACGTGAGTTTTCGTTCCACTGAGCGTCAGACCCCGTA
GAAAAGATCAAAGGATCTTCTTGAGATCCTTTTTTTCTGCGCGTAATCTGCTGCTTGCAAACAAAAAAACCACCGCTACCAGCGGTGGTTTGTTTGCC
GGATCAAGAGCTACCAACTCTTTTTCCGAAGGTAACTGGCTTCAGCAGAGCGCAGATACCAAATACTGTCCTTCTAGTGTAGCCGTAGTTAGGCCAC
CACTTCAAGAACTCTGTAGCACCGCCTACATACCTCGCTCTGCTAATCCTGTTACCAGTGGCTGCTGCCAGTGGCGATAAGTCGTGTCTTACCGGGT
TGGACTCAAGACGATAGTTACCGGATAAGGCGCAGCGGTCGGGCTGAACGGGGGGTTCGTGCACACAGCCCAGCTTGGAGCGAACGACCTACA
CCGAACTGAGATACCTACAGCGTGAGCTATGAGAAAGCGCCACGCTTCCCGAAGGGAGAAAGGCGGACAGGTATCCGGTAAGCGGCAGGGTCG
GAACAGGAGAGCGCACGAGGGAGCTTCCAGGGGGAAACGCCTGGTATCTTTATAGTCCTGTCGGGTTTCGCCACCTCTGACTTGAGCGTCGATTT
TTGTGATGCTCGTCAGGGGGGCGGAGCCTATGGAAAAACGCCAGCAACGCGGCCTTTTTACGGTTCCTGGGCTTTTGCTGGCCTTTTGCTCACAT
GTTCTTTCCTGCGTTATCCCCTGATTCTGTGGATAACCGTATTACCGCCTTTGAGTGAGCTGATACCGCTCGCCGCAGCCGAACGACCGAGCGCAG
CGAGTCAGTGAGCGAGGAAGCGGAAGAGCGCCCAATACGCAAACCGCCTCTCCCCGCGCGTTGGCCGATTCATTAATGCAG

>extended EDA Sll0107: [ Synechocystis sp. PCC 6803]
MSIIDDFGCPQESAISALRSPWLKMLRQHRAIAVIRTDSIDLSLQLAKVAIEAGMGLVEITWNSDQPGETIAHLRHQFPHCAIGTGTVLSQEDLLQAVASG
AQFCFTPHSEPELITSAVAHGIPIIPGAFSPTEIVQAWQRGATAVKVFPIKTLGGVDYIQALQGPLGQIPLIPTGGVTLANAQAFLAAGAIAVGLSGQLFPPNL
VAIKAWREIGDLIRTLLKEIQTPRG

Original sequence
ATGTCCATCATTGATGATTTTGGTTGCCCCCAGGAGTCAGCTATCTCCGCCCTCCGATCGCCATGGTTAAAAATGTTGCGACAGCACCGGGCGATC
GCCGTTATTCGCACTGATTCCATCGACCTAAGTCTGCAATTGGCTAAGGTGGCCATCGAAGCAGGCATGGGGCTAGTGGAAATTACCTGGAACAGT
GACCAGCCGGGGGAAACCATTGCTCACCTGCGTCATCAATTTCCCCATTGTGCCATTGGTACGGGTACAGTCCTCTCTCAGGAGGATTTGCTCCAG
GCCGTTGCCAGTGGAGCCCAATTTTGCTTTACCCCCCACAGTGAGCCAGAATTAATTACCAGCGCCGTGGCCCATGGCATCCCCATTATTCCCGG
TGCCTTTTCCCCTACGGAGATTGTCCAAGCCTGGCAGAGGGGAGCCACCGCCGTCAAAGTTTTTCCCATTAAAACCTTGGGAGGAGTGGACTACAT
TCAAGCTCTCCAAGGTCCCCTCGGCCAAATTCCCCTTATTCCCACCGGAGGGGTAACTTTGGCAAATGCCCAAGCTTTCCTAGCGGCCGGAGCG
ATCGCCGTTGGCCTATCTGGACAGCTATTCCCCCCCAATTTAGTAGCAATCAAAGCCTGGAGAGAAATTGGCGATCTAATCCGTACTTTGTTGAAAGA
AATCCAGACACCAAGGGGCTGA

Codon optimized (in red) sequence
ATGAGCATTATCGATGATTTCGGCTGTCCGCAGGAAAGCGCAATTAGCGCCCTGCGCAGTCCGTGGCTGAAAATGCTGCGCCAGCATCGCGCAAT
TGCAGTTATTCGCACCGATAGTATTGATCTGAGCCTGCAGCTGGCCAAAGTGGCAATTGAAGCAGGTATGGGTCTGGTTGAAATTACCTGGAATAGC
GATCAGCCGGGCGAAACCATTGCCCATCTGCGTCATCAGTTTCCGCATTGTGCCATTGGCACCGGTACCGTGCTGAGCCAGGAAGATCTGCTGCA
GGCAGTTGCAAGCGGCGCACAGTTTTGTTTTACCCCGCATAGTGAACCGGAACTGATTACCAGCGCAGTGGCACATGGCATTCCGATTATTCCGG
GCGCATTTTCACCGACCGAAATTGTGCAGGCCTGGCAGCGTGGCGCAACCGCTGTTAAAGTGTTTCCGATTAAGACCCTGGGTGGTGTTGATTATAT
TCAGGCCCTGCAGGGTCCGCTGGGTCAGATTCCGCTGATTCCGACCGGTGGTGTGACCCTGGCCAATGCACAGGCATTTCTGGCCGCCGGTGC
CATTGCAGTTGGTCTGAGCGGTCAGCTGTTTCCGCCGAATCTGGTGGCCATTAAGGCCTGGCGTGAAATTGGTGACCTGATTCGCACCCTGCTGAA
AGAAATTCAGACCCCGCGTGGTTAA

**Figure S7: Cloning strategy and sequences for overexpression of putative GDHs (NEO87985.1and WP_238987112 and) and GKs (NEO86388.1 and WP_023068790.1) from *Spirulina sp. SIO3F2* and *Lyngbya aestuarii* and codon optimized sequence of EDA (Sll0107).**

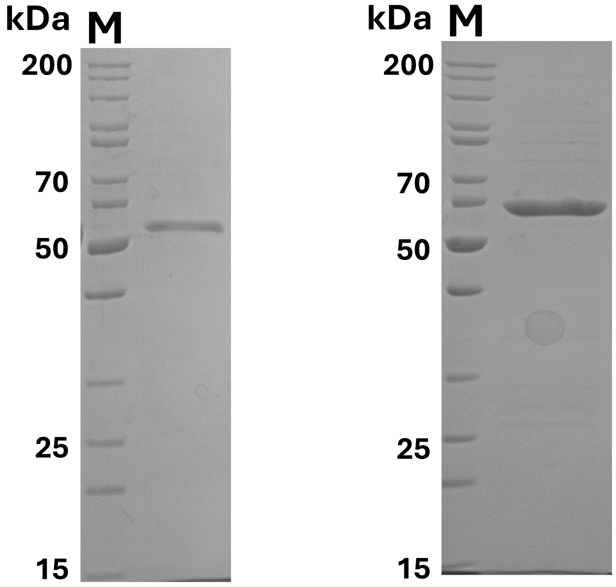


**Figure S8: Purification of the recombinant GDH homologues from *Spirulina* sp. SIO3F2 (NEO87985.1) (left panel) and *Lyngbya aestuarii* (WP_238987112) (right panel) after expression in *E.coli*.** Purified protein (2 µg) following IMAC was analyzed by SDS-PAGE and visualized by staining with Coomassie Brilliant Blue (before TEV cleavage). M, molecular weight marker (PageRuler™ Unstained Protein Ladder, Thermo Fisher Scientific). The apparent molecular masses of GST-tagged GDH from *Spirulina* sp. SIO3F2 (~55 kDa) and *L. aestuarii* (~60 kDa) agree with the calculated masses of 55.5 kDa and 57.1 kDa, respectively.

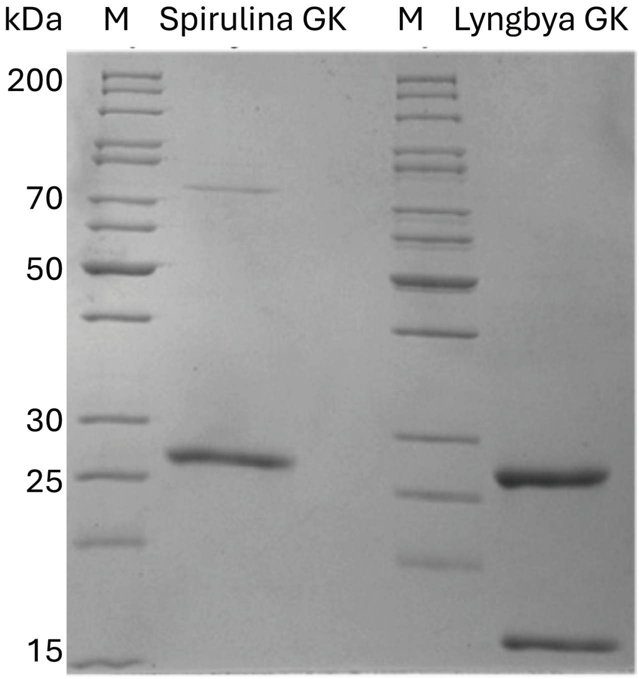


**Figure S9: Purification of the GK from Spirulina sp. SIO3F2 and Lyngbya aestuarii after recombination expression in E. coli**. Both GKs were expressed with GST-His tags and the protein after Immobilized Metal Ion Affinity Chromatography (IMAC) and TEV digestion is shown. Only for L. aestuarii GK the TEV cleavage was successful, and a specific activity of 24.8 U/mg protein could be determined (table 1). The expected size for gluconate kinase is 17-19 kDa.

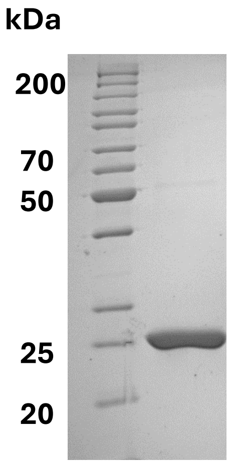


**Figure S10. Purification of the recombinant extended version of Sll0107 (EDA) from *Synechocystis* sp. PCC 6803 after expression in *E. coli*.** Purified protein (2 µg) following IMAC and size-exclusion chromatography was analyzed by SDS–PAGE and visualized with Coomassie Brilliant Blue. M, molecular weight marker (PageRuler™ Unstained Protein Ladder, Thermo Fisher Scientific). The apparent molecular mass (~25 kDa) is consistent with the calculated mass of 23 kDa.

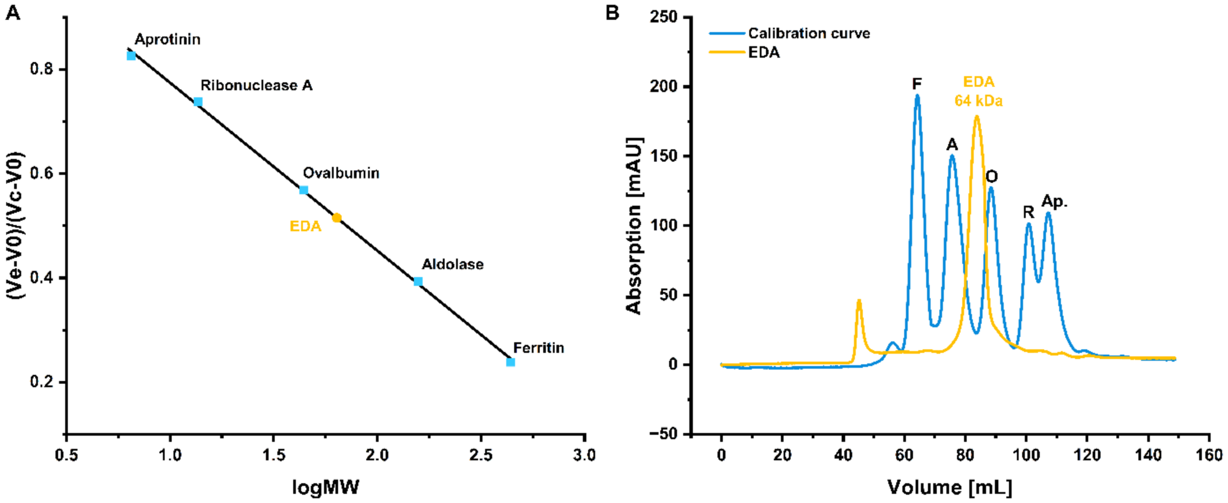


**Figure S11. Native molecular mass of Sll0107 (EDA) from *Synechocystis*.** (A) The native molecular mass of EDA was determined via size exclusion chromatography (HiLoad 16/600 Superdex 200 prep grade, Cytiva, Marlborough, MA, USA)). (B) Calibration curve with five proteins (Aprotinin (6.5 kDa), Ribonuclease A (13.7 kDa), Ovalbumin (44 kDa), Aldolase (158 kDa) and Ferritin (440 kDa) from the LMW and HMW gel filtration calibration kits (GE Healthcare). The native molecular mass revealed a single peak at approximately 64 kDa, consistent with a homotrimeric oligomeric state for Sll0107 (EDA), which is characteristic for bacterial KDPG aldolases (5, 6).

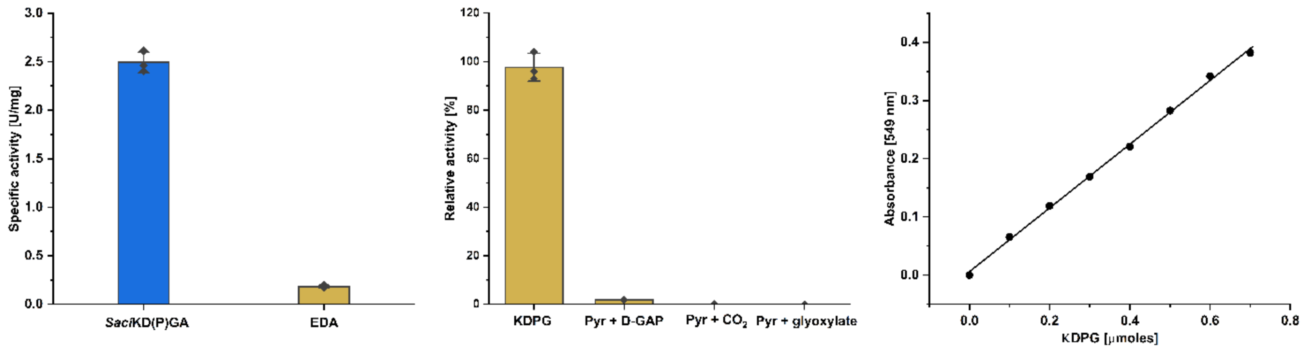


**Figure S12: Enzymatic activity of *Synechocystis* EDA in the condensation (reverse) direction, determined by the thiobarbituric acid (TBA) assay.** (A) Formation of KDPG from 20 mM pyruvate and 10 mM D-GAP was assayed as the condensation reaction. Assays were performed discontinuously in 50 mM Tris-HCL, 300 mM NaCl, pH 7.8. Purified *Sulfolobus acidocaldarius* KD(P)G aldolase (*Saci*KD(P)GA) (3µg) was used as positive control; 15 µg of *Synechocystis* EDA was tested in parallel. Incubation temperatures were 70 °C for *Saci*KD(P)GA and 30 °C for *Synechocystis* EDA. (B) A comparison of relative activities of EDA in the forward (KDPG cleavage) and reverse directions (pyruvate + D-GAP, pyruvate + $CO_2$, and pyruvate + glyoxylate condensation) are shown. (C) A KDPG calibration curve generated under TBA assay conditions was used to quantify the KDPG produced by EDA in the condensation reaction. From these measurements, the specific activity of EDA in the condensation direction (pyruvate + D-GAP → KDPG) was 0.18 U/mg whereas pyruvate + $CO_2$ and pyruvate + glyoxylate did not serve as substrates. Data are presented as mean ± SD of three technical replicates (n = 3).

**Table S1** Result of blast analyses with 261 cyanobacteria next to *Synechocystis* based on a gluconate kinase (GK) of *Escherichia coli* and two glucose dehydrogenases (GDHs) as baits. As GDH baits the NAD$^{+}$-dependent GDH 1GCO from *Bacillus megaterium* and the quinone-dependent GDH from *Acinetobacter calcoaceticus* were used. Only cyanobacteria are listed that contain at least one GK and one GDH homolog that met our threshold settings of a query score > 30 %, an identity score > 30 % and an e-value < $1^{-10}$. In addition, *Synechocystis* sp.PCC 6803 is shown.

QS – Query score in %, IS – Identity score in %, AN – Accession number.

| | | 1GCO NAD+ | | | | 1C9U Chinon | | | | E. coli Gk | | |
|---|---|---|---|---|---|---|---|---|---|---|---|---|
| Organism | QS % | e value | IS % | AN | QS % | e value | IS % | AN | QS % | e value | IS % | AN |
| unclassified Synechocystis | 95 | 1E-48 | 37.2 | WP_010872244.1 | 92 | 2.E-31 | 27.08 | WP_010871649.1 | 16 | 0.39 | 41.38 | WP_010873541.1 |
| | 95 | 1E-45 | 37.35 | WP_041428273.1 | 5 | 2.3 | 38.46 | WP_010873285.1 | | | | |
| | 92 | 2E-31 | 33.6 | WP_010872685.1 | | | | | | | | |
| | 74 | 2E-21 | 31.28 | WP_010873912.1 | | | | | | | | |
| Synechocystis sp. PCC 6803 | 95 | 2E-45 | 37.05 | AGF53506.1 | | | | | | | | |
| Acaryochloridaceae cyanobacterium | 94 | 2E-30 | 33.33 | NJM74450.1 | | | | | 96 | 3E-29 | 37.43 | NJM74503.1 |
| Acaryochloris marina | 94 | 2E-44 | 36.44 | WP_012165802.1 | | | | | 86 | 4E-24 | 35.29 | WP_012164422.1 |
| Acaryochloris sp. CCMEE 5410 | 98 | 2E-82 | 47.73 | WP_050857446.1 | 55 | 1E-19 | 29.12 | WP_039779973.1 | 86 | 4E-23 | 33.55 | WP_010471463.1 |
| Acaryochloris sp. RCC1774 | 98 | 2E-78 | 47.73 | WP_110988267.1 | 55 | 7E-11 | 27.56 | PZD73694.1 | 90 | 3E-35 | 39.62 | WP_110985040.1 |
| Aliterella atlantica | 95 | 8E-51 | 38.15 | WP_045053909.1 | | | | | 91 | 4E-35 | 38.75 | WP_045055410.1 |
| Alkalinema sp. CACIAM 70d | 94 | 2E-68 | 40.23 | OUC14139.1 | 90 | 7E-32 | 28.64 | OUC11796.1 | 86 | 3E-37 | 44.37 | OUC15726.1 |
| Anabaena cylindrica | 96 | 2E-83 | 47.89 | WP_015215175. | | | | | 86 | 6E-36 | 42.76 | WP_015213074.1 |
| Anabaena sp. | 97 | 5E-83 | 46.77 | HFS11400.1 | | | | | 89 | 2E-30 | 35.84 | HFS08018.1 |
| Anabaena sp. 4-3 | 95 | 2E-52 | 38.96 | WP_066423504.1 | 73 | 5E-19 | 25.00 | WP_066426784.1 | 89 | 2E-39 | 43.31 | WP_066425370.1 |
| Anabaena sp. CA = ATCC 33047 | 95 | 5E-52 | 38.55 | WP_066379214.1 | 73 | 4E-19 | 25.00 | WP_066383080.1 | 89 | 4E-39 | 43.31 | WP_066384767.1 |

| | | | | | | | | | | | | |
|---|---|---|---|---|---|---|---|---|---|---|---|---|
| Anabaena sp. CRKS33 | 89 | 7E-28 | 32.49 | OBQ36656.1 | | | | | 92 | 4E-35 | 44.10 | OBQ37897.1 |
| Anabaena sp. UHCC 0204 | 70 | 1E-25 | 35.68 | WP_168496307.1 | | | | | 90 | 2E-36 | 43.40 | WP_168497267.1 |
| Anabaena sp. UHCC 0253 | 72 | 1E-25 | 34.74 | WP_168535874.1 | | | | | 90 | 4E-36 | 43.40 | WP_168538208.1 |
| Anabaenopsis circularis | 93 | 3E-40 | 33.60 | WP_096577178.1 | | | | | 86 | 4E-33 | 38.41 | WP_096579879.1 |
| Aphanizomenon | 94 | 1E-52 | 38.87 | WP_039201825.1 | | | | | 90 | 3E-36 | 43.40 | WP_039200411.1 |
| Aphanocapsa montana | 94 | 3E-69 | 41.46 | WP_166279655.1 | 52 | 6E-11 | 28.68 | WP_166282873.1 | 92 | 7E-32 | 36.36 | WP_166281212.1 |
| Aphanothece hegewaldii | 95 | 5E-57 | 40.40 | WP_106456819.1 | | | | | 92 | 2E-35 | 40.74 | WP_106458100.1 |
| Brasilonema | 97 | 9E-86 | 48.85 | WP_171976650.1 | | | | | 88 | 7E-38 | 45.16 | WP_171977886.1 |
| Brasilonema bromeliae | 94 | 2E-49 | 38.46 | WP_169157777.1 | | | | | 88 | 2E-38 | 45.16 | WP_169155174.1 |
| Brasilonema octagenarum | 95 | 7E-68 | 41.20 | WP_169265890.1 | | | | | 88 | 1E-38 | 45.16 | WP_169266841.1 |
| Brasilonema sp. UFV-L1 | 97 | 4E-86 | 49.24 | WP_169220422.1 | | | | | 88 | 5E-38 | 44.52 | WP_169219012.1 |
| Calothrix brevissima | 96 | 9E-76 | 44.79 | WP_096646716.1 | | | | | 86 | 1E-31 | 39.07 | WP_096647816.1 |
| Calothrix brevissima NIES-22 | | | | | | | | | 88 | 8E-33 | 38.71 | BAY64024.1 |
| Calothrix desertica | 97 | 1E-86 | 49.43 | WP_127082149.1 | 92 | 1E-28 | 28.61 | WP_127079297.1 | 88 | 9E-34 | 40.65 | WP_127080590.1 |
| Calothrix parasitica | 97 | 3E-84 | 48.85 | WP_096655843.1 | 90 | 2E-35 | 28.47 | WP_096657602.1 | 91 | 1E-38 | 41.88 | WP_096660815.1 |
| Calothrix parietina | 95 | 3E-46 | 36.14 | WP_015197185.1 | 73 | 3E-16 | 24.48 | WP_015199424.1 | 77 | 7E-30 | 40.28 | WP_083866341.1 |
| Calothrix sp. CSU_2_0 | 95 | 2E-46 | 34.80 | NJR14504.1 | | | | | 93 | 2E-32 | 38.41 | NJR16580.1 |
| Calothrix sp. HK-06 | 96 | 1E-76 | 45.38 | WP_073619531.1 | 92 | 6E-29 | 28.61 | WP_073623789.1 | 88 | 2E-34 | 41.94 | WP_073624506.1 |
| Calothrix sp. NIES-2098 | 97 | 7E-84 | 47.53 | WP_096590940.1 | 53 | 4E-10 | 27.97 | WP_096592248.1 | 86 | 8E-37 | 40.40 | WP_096595320.1 |
| Calothrix sp. NIES-2100 | 96 | 5E-76 | 44.02 | WP_096602347.1 | | | | | 92 | 1E-35 | 40.12 | BAY26662.1 |
| Calothrix sp. NIES-3974 | 97 | 5E-85 | 48.67 | WP_096620359.1 | 88 | 2E-35 | 30.49 | WP_096620443.1 | 84 | 7E-32 | 38.10 | WP_172892798.1 |
| Calothrix sp. NIES-4071 | 93 | 4E-43 | 35.22 | BAZ09942.1 | | | | | 88 | 4E-33 | 41.29 | BAZ17549.1 |
| Calothrix sp. NIES-4101 | 97 | 1E-82 | 47.15 | BAZ37791.1 | 90 | 1E-26 | 28.30 | BAZ41358.1 | 84 | 5E-30 | 39.46 | BAZ39688.1 |

| | | | | | | | | | | | | |
|---|---|---|---|---|---|---|---|---|---|---|---|---|
| Calothrix sp. PCC 6303 | | | | | | | | | 74 | 9E-29 | 40.71 | AFZ04079.1 |
| Calothrix sp. PCC 7103 | 97 | 3E-87 | 49.43 | WP_019495089.1 | 92 | 2E-30 | 29.08 | WP_019497261.1 | 88 | 2E-33 | 40.65 | WP_019495466.1 |
| Calothrix sp. PCC 7507 | 94 | 3E-74 | 43.03 | WP_015129873.1 | | | | | 89 | 9E-31 | 41.40 | WP_015128726.1 |
| Chamaesiphon minutus | 95 | 7E-49 | 38.65 | WP_015161590.1 | 92 | 4E-30 | 27.17 | WP_157260219.1 | 88 | 1E-39 | 40.00 | WP_015158279.1 |
| Chamaesiphon polymorphus | 97 | 2E-81 | 46.21 | WP_106303519.1 | 92 | 5E-27 | 26.74 | WP_106310332.1 | 88 | 1E-39 | 40.65 | WP_106300099.1 |
| Chlorogloea sp. CCALA 695 | 94 | 4E-74 | 42.28 | WP_106369023.1 | | | | | 90 | 3E-36 | 42.14 | WP_106370661.1 |
| Chlorogloeopsis fritschii | 97 | 1E-82 | 47.91 | WP_016877929.1 | 56 | 8E-15 | 28.52 | WP_016872983.1 | 86 | 2E-32 | 39.07 | WP_016872994.1 |
| Chondrocystis sp. NIES-4102 | 97 | 1E-84 | 49.05 | WP_096717707.1 | | | | | 92 | 6E-37 | 39.75 | WP_096719751.1 |
| Chroococcidiopsis sp. PCC 6712 | 97 | 2E-81 | 45.59 | WP_169245013.1 | | | | | 84 | 1E-32 | 39.60 | WP_169242370.1 |
| Coleofasciculus cht. | 94 | 2E-50 | 39.68 | WP_006101550.1 | 93 | 1E-30 | 26.64 | WP_006104631.1 | 51 | 3E-19 | 45.56 | WP_071777273.1 |
| Coleofasciculus cht. PCC 7420 | 95 | 3E-44 | 35.46 | WP_131115089.1 | | | | | 93 | 3E-32 | 39.88 | WP_131119806.1 |
| Crinalium epipsammum | 97 | 1E-82 | 48.67 | WP_015204932.1 | | | | | 90 | 5E-34 | 38.99 | WP_015202021.1 |
| Crocosphaera chwakensis | 96 | 7E-83 | 47.10 | WP_008275623.1 | 92 | 5E-29 | 28.87 | WP_008276783.1 | 90 | 7E-41 | 41.51 | WP_008273788.1 |
| Crocosphaera subtropica | 97 | 1E-81 | 46.36 | WP_009546719.1 | 91 | 3E-27 | 27.27 | WP_009547689.1 | 90 | 4E-42 | 42.14 | WP_009545789.1 |
| Crocosphaera subtropica ATCC 51142 | 94 | 2E-46 | 37.25 | ACB51761.1 | | | | | 90 | 5E-42 | 42.14 | ACB51335.1 |
| Crocosphaera watsonii | 94 | 5E-48 | 38.06 | WP_007309538.1 | 90 | 2E-23 | 26.38 | WP_156185059.1 | 91 | 4E-44 | 43.12 | WP_007311533.1 |
| Cyanobacteria | 97 | 1E-85 | 49.05 | WP_106869782.1 | | | | | 90 | 3E-37 | 41.36 | WP_039727538.1 |
| Cyanobacteria bacterium 13_1_20CM_4_61_6 | 94 | 3E-66 | 43.32 | OLE96307.1 | | | | | 82 | 6E-30 | 37.50 | OLE97478.1 |
| Cyanobacteria bact. J069 | 97 | 8E-86 | 48.29 | RMF66847.1 | 88 | 5E-29 | 28.61 | RMF63493.1 | 81 | 7E-20 | 39.31 | RMF63274.1 |
| Cyanobacteria bact. UBA11162 | 93 | 5E-43 | 34.82 | HBL13216.1 | | | | | 85 | 6E-37 | 44.00 | HBL10089.1 |
| Cyanobacteria bact. UBA11371 | 97 | 3E-86 | 48.67 | HAZ48147.1 | 90 | 5E-27 | 26.27 | HAZ45289.1 | 92 | 5E-39 | 39.02 | HAZ46796.1 |
| Cyanobacteria bact.UBA11372 | 97 | 4E-86 | 48.29 | HAX77846.1 | | | | | 92 | 1E-38 | 39.02 | HAX74250.1 |
| Cyanobacteria bact. UBA12227 | 90 | 1E-46 | 39.24 | HAG85715.1 | | | | | 85 | 4E-37 | 44.00 | HAG81504.1 |
| cyanobacterium endosymbiont of Epithemia turgida | 94 | 7E-50 | 36.69 | WP_044107707.1 | | | | | 90 | 3E-41 | 42.14 | WP_044107338.1 |

| | | | | | | | | | | | | |
|---|---|---|---|---|---|---|---|---|---|---|---|---|
| cyanobacterium endosymbiont of Epithemia turgida isolate EtSB Lake Yunoko | 94 | 1E-49 | 36.69 | BAP18592.1 | | | | | 88 | 8E-39 | 41.94 | BAP18489.1 |
| cyanobacterium endosymbiont of Rhopalodia gibberula | 94 | 1E-49 | 36.69 | WP_119260189.1 | | | | | 92 | 4E-41 | 41.98 | WP_119260280.1 |
| cyanobacterium PCC 7702 | 95 | 1E-48 | 38.15 | WP_017321921.1 | | | | | 87 | 7E-36 | 41.18 | WP_038315641.1 |
| Cyanothece sp. BG0011 | 94 | 8E-48 | 36.84 | WP_107667288.1 | 91 | 3E-27 | 27.92 | WP_107667132.1 | 90 | 1E-42 | 44.65 | WP_107667392.1 |
| Cyanothece sp. PCC 7425 | 94 | 8E-41 | 36.00 | WP_012627413.1 | 93 | 3E-31 | 27.71 | WP_156793796.1 | 90 | 2E-34 | 37.74 | WP_012626255.1 |
| Cyanothece sp. SIO1E1 | 97 | 3E-80 | 47.33 | NET35852.1 | 57 | 4E-23 | 30.60 | NET34378.1 | 88 | 3E-35 | 40.65 | NET34717.1 |
| Cyanothece sp. SIO2G6 | 94 | 2E-43 | 35.63 | NEQ96279.1 | 92 | 4E-38 | 29.55 | NEQ96349.1 | 81 | 3E-22 | 39.58 | NER00940.1 |
| Cyanothece sp. UBA12306 | 95 | 4E-39 | 34.66 | HAC64137.1 | 90 | 4E-30 | 27.95 | HAC62366.1 | 90 | 3E-42 | 42.14 | HAC64422.1 |
| Cylindrospermum sp. NIES-4074 | 94 | 1E-38 | 34.25 | BAZ33671.1 | | | | | 86 | 3E-37 | 43.05 | BAZ32045.1 |
| Cylindrospermum stagnale | 97 | 8E-75 | 45.25 | WP_015209547.1 | | | | | 88 | 2E-33 | 38.71 | WP_015207172.1 |
| Dolichospermum circinale | 94 | 6E-53 | 38.87 | WP_028089931.1 | | | | | 92 | 7E-34 | 42.86 | WP_035371736.1 |
| Dolichospermum compactum | 89 | 3E-30 | 33.76 | WP_096670462.1 | | | | | 88 | 4E-36 | 43.87 | WP_096666855.1 |
| Dolichospermum planctonicum | 94 | 7E-53 | 38.87 | WP_168357981.1 | | | | | 94 | 2E-34 | 43.64 | WP_137908858.1 |
| Dolichospermum sp. UHCC 0259 | 94 | 1E-52 | 38.46 | WP_168643508.1 | | | | | 88 | 9E-37 | 44.52 | WP_168646187.1 |
| Dolichospermum sp. UHCC 0260 | 95 | 4E-43 | 37.55 | WP_168633154.1 | | | | | 88 | 1E-40 | 42.58 | WP_174783200.1 |
| filamentous cyanobacterium ESFC-1 | 97 | 5E-91 | 50.38 | WP_018398467.1 | 89 | 1E-24 | 26.03 | WP_026222337.1 | 88 | 5E-38 | 40.91 | WP_018400815.1 |
| Fischerella | 70 | 5E-30 | 38.92 | WP_062250401.1 | | | | | 91 | 1E-35 | 41.88 | WP_062249050.1 |
| Fischerella major | 97 | 1E-84 | 48.67 | WP_073554987.1 | | | | | 91 | 1E-36 | 42.50 | WP_073554785.1 |
| Fischerella muscicola | 97 | 2E-84 | 48.67 | WP_016865909.1 | 68 | 9E-18 | 25.08 | WP_016865913.1 | 91 | 4E-35 | 41.25 | WP_016866276.1 |
| Fischerella sp. NIES-3754 | 97 | 3E-85 | 49.05 | WP_062242560. | | | | | 90 | 6E-35 | 41.77 | BAU08163.1 |
| Fischerella sp. NIES-4106 | 94 | 2E-76 | 43.09 | WP_096679645.1 | | | | | 93 | 2E-32 | 39.88 | WP_096675048.1 |
| Fischerella sp. PCC 9339 | 97 | 3E-83 | 48.86 | WP_017307303.1 | | | | | 93 | 2E-31 | 39.26 | WP_017310421.1 |
| Fischerella sp. PCC 9431 | 94 | 1E-69 | 40.23 | WP_026721732.1 | | | | | 90 | 6E-35 | 40.88 | WP_035121874.1 |

| | | | | | | | | | | | | |
|---|---|---|---|---|---|---|---|---|---|---|---|---|
| Fischerella sp. PCC 9605 | 94 | 3E-45 | 37.85 | WP_026735398.1 | | | | | 86 | 2E-33 | 39.74 | WP_035139369.1 |
| Fischerella thermalis | 97 | 1E-84 | 48.67 | WP_102207332.1 | 68 | 2E-18 | 25.71 | WP_102173240.1 | 91 | 2E-37 | 43.12 | WP_102187211.1 |
| Fortiea contorta | 97 | 2E-82 | 47.15 | WP_017654414.1 | 54 | 3E-13 | 26.64 | WP_017652856.1 | 89 | 1E-32 | 40.13 | WP_017651834.1 |
| Geitlerinema sp. FC II | 95 | 8E-48 | 38.96 | PPT05270.1 | | | | | 77 | 3E-35 | 38.97 | PPT08383.1 |
| Geitlerinema sp. P-1104 | 97 | 1E-82 | 48.66 | WP_170191022.1 | 90 | 7E-30 | 30.17 | WP_170188606.1 | 87 | 5E-32 | 37.58 | WP_170190523.1 |
| Geitlerinema sp. PCC 7105 | 95 | 2E-48 | 39.76 | WP_017661427.1 | | | | | 77 | 3E-37 | 40.44 | WP_017663490.1 |
| Geitlerinema sp. PCC 9228 | 95 | 1E-50 | 38.96 | WP_071518111.1 | | | | | 92 | 3E-36 | 38.51 | WP_139240796.1 |
| Gloeobacter kilaueensis | 94 | 7E-74 | 42.28 | WP_023175839.1 | | | | | 86 | 1E-26 | 35.67 | WP_041243890.1 |
| Gloeobacter kilaueensis JS1 | 73 | 3E-22 | 34.36 | AGY59051.1 | | | | | 72 | 3E-20 | 36.64 | AGY58430.1 |
| Gloeobacter violaceus | 98 | 3E-74 | 45.32 | WP_011141114.1 | 56 | 2E-19 | 29.26 | WP_011142258.1 | 85 | 1E-25 | 36.18 | WP_011142812.1 |
| Gloeobacter violaceus SpSt-379 | 94 | 1E-27 | 33.07 | NJR46195.1 | | | | | 88 | 8E-33 | 38.96 | NJR45610.1 |
| Gloeocapsa sp. PCC 73106 | 94 | 2E-70 | 43.58 | WP_006527119.1 | | | | | 78 | 4E-32 | 42.75 | WP_006528895.1 |
| Gloeothece citriformis | 97 | 7E-71 | 47.15 | WP_015955892.1 | | | | | 89 | 2E-41 | 43.59 | WP_015956828.1 |
| Gloeothece verrucosa | 95 | 1E-52 | 38.15 | WP_013324621.1 | | | | | 89 | 6E-40 | 44.23 | WP_013322699.1 |
| Halomicronema hongdechloris | 94 | 5E-48 | 38.06 | WP_007309538.1 | | | | | 91 | 2E-44 | 43.12 | WP_007306380.1 |
| Halomicronema hongdechloris C2206 | 94 | 5E-50 | 38.46 | WP_088431545.1 | | | | | 86 | 3E-29 | 41.72 | WP_080811080.1 |
| Hapalosiphon sp. MRB220 | | | | | | | | | 88 | 1E-30 | 41.29 | ASC70927.1 |
| Hapalosiphonaceae cyanobacterium JJU2 | 96 | 1E-72 | 42.47 | WP_053456449.1 | | | | | 90 | 2E-34 | 40.88 | WP_053456262.1 |
| Hassallia byssoidea | 94 | 1E-78 | 43.90 | RAM49828.1 | | | | | 93 | 9E-32 | 39.26 | RAM49355.1 |
| HrmK Nostoc cycadae WK-1 | 95 | 4E-68 | 40.40 | WP_039744423.1 | | | | | 88 | 2E-39 | 43.87 | WP_039752768.1 |
| Hydrococcus rivularis | | | | | | | | | 88 | 2E-33 | 38.06 | GBE92088.1 |
| Hydrococcus sp. RU_2_2 | 97 | 1E-84 | 47.91 | WP_073598102.1 | | | | | 92 | 1E-32 | 39.51 | WP_073600744.1 |
| Hydrococcus sp. SU_1_0 | 97 | 1E-88 | 49.81 | NJM87157.1 | 93 | 1E-31 | 28.47 | NJM89692.1 | 88 | 2E-39 | 42.86 | NJM87876.1 |
| Hydrocoleum sp. CS-953 | 94 | 5E-40 | 35.22 | NJL53579.1 | | | | | 77 | 1E-26 | 37.04 | NJL53618.1 |

| | | | | | | | | | | | | |
|---|---|---|---|---|---|---|---|---|---|---|---|---|
| Hyella patelloides | 96 | 7E-55 | 40.39 | WP_094673338.1 | 53 | 5E-19 | 29.27 | OZH53923.1 | 88 | 3E-39 | 42.58 | WP_094671305.1 |
| Hyella patelloides LEGE 07179 | | | | | | | | | 74 | 8E-36 | 43.94 | VEP16132.1 |
| Hyellaceae cyanobacterium CSU_1_1 | 97 | 3E-88 | 50.57 | WP_144872141.1 | 92 | 6E-35 | 28.24 | WP_144876011.1 | 76 | 3E-37 | 43.70 | WP_144874985.1 |
| Leptolyngbya frigida | | | | | | | | | 90 | 4E-28 | 35.85 | HGZ85914.1 |
| Leptolyngbya sp. | 94 | 9E-75 | 43.24 | WP_106259022.1 | | | | | 89 | 8E-41 | 43.59 | WP_106259403.1 |
| Leptolyngbya sp. DLM2.Bin15 | 97 | 3E-84 | 48.67 | PZV08269.1 | | | | | 92 | 1E-23 | 34.55 | PZV07176.1 |
| Leptolyngbya sp. DLM2.Bin27 | 94 | 4E-47 | 36.03 | TVQ23819.1 | 92 | 1E-26 | 26.11 | TVQ18876.1 | 86 | 2E-41 | 45.03 | TVQ18967.1 |
| Leptolyngbya sp. KIOST-1 | 97 | 2E-82 | 47.91 | TVQ06243.1 | 91 | 6E-27 | 27.38 | TVQ09982.1 | 89 | 3E-31 | 38.46 | TVQ08450.1 |
| Leptolyngbya sp. O-77 | 97 | 2E-80 | 47.53 | WP_035986605.1 | 91 | 5E-23 | 27.32 | WP_052050691.1 | 91 | 1E-38 | 41.88 | WP_035984714.1 |
| Leptolyngbya sp. O-77 | 94 | 1E-64 | 38.43 | WP_068511348.1 | 67 | 2E-28 | 29.75 | WP_084783046.1 | 90 | 2E-21 | 33.95 | BAU42100.1 |
| Leptolyngbya sp. PCC 7375 | 94 | 1E-64 | 38.43 | WP_068511348.1 | 67 | 2E-28 | 29.75 | WP_084783046.1 | 88 | 5E-20 | 34.81 | WP_068516305.1 |
| Leptolyngbya sp. SIO3F4 | 96 | 2E-48 | 38.10 | WP_006514312.1 | 91 | 6E-29 | 26.71 | EKV00189.1 | 81 | 1E-23 | 38.03 | WP_156828702.1 |
| Leptolyngbya sp. SIOISBB | 94 | 2E-74 | 43.63 | NEQ53329.1 | 93 | 1E-27 | 26.39 | NEQ52188.1 | 81 | 2E-22 | 38.62 | NEQ50907.1 |
| Leptolyngbya valderiana BDU 20041 | 95 | 6E-43 | 36.25 | NEQ43968.1 | 92 | 2E-34 | 28.13 | NEQ42753.1 | 84 | 8E-42 | 47.33 | NEQ43478.1 |
| Leptolyngbyaceae cyanobacterium CCMR0081 | 87 | 3E-19 | 28.70 | OAB63225.1 | | | | | 74 | 3E-33 | 39.69 | OAB59984.1 |
| Leptolyngbyaceae cyanobacterium CCMR0082 | 94 | 9E-44 | 37.50 | WP_163698381.1 | 91 | 1E-27 | 25.70 | WP_163702678.1 | 83 | 8E-27 | 39.04 | WP_163699777.1 |
| Leptolyngbyaceae cyanobacterium CSU_1_3 | 96 | 4E-46 | 37.11 | WP_163659223.1 | 91 | 1E-27 | 25.70 | WP_163668558.1 | 83 | 1E-25 | 38.36 | WP_163664438.1 |
| Leptolyngbyaceae cyanobacterium CSU_1_4 | 97 | 9E-88 | 49.43 | NJR51820.1 | | | | | 88 | 6E-41 | 43.23 | NJR50069.1 |
| Leptolyngbyaceae cyanobacterium SM2_3_12 | 94 | 2E-65 | 38.46 | NJR38065.1 | 60 | 3E-18 | 26.43 | NJR39066.1 | 88 | 1E-37 | 43.87 | NJR40752.1 |
| Leptolyngbyaceae cyanobacterium SM2_5_2 | | | | | | | | | 90 | 6E-41 | 45.28 | NJL45456.1 |
| Lyngbya aestuarii | 97 | 6E-83 | 47.89 | WP_238987112 | 90 | 4E-30 | 27.27 | WP_023068940.1 | 88 | 2E-35 | 40.00 | WP_023068790.1 |
| Mastigocladopsis repens | 97 | 5E-84 | 48.28 | WP_009783137.1 | 90 | 8E-29 | 27.45 | EAW35290.1 | 92 | 4E-37 | 39.51 | WP_009785896.1 |
| Mastigocladus laminosus | 97 | 2E-85 | 48.09 | WP_017314104.1 | | | | | 88 | 1E-36 | 43.87 | WP_026082510.1 |
| Mastigocoleus testarum | 95 | 5E-50 | 39.68 | WP_135107550.1 | | | | | 90 | 9E-31 | 39.62 | WP_035115738.1 |
| Merismopedia glauca | 97 | 8E-83 | 46.97 | WP_027846246.1 | 90 | 1E-32 | 28.30 | WP_027846436.1 | 93 | 4E-43 | 42.33 | WP_027843784.1 |

| | | | | | | | | | | | | |
|---|---|---|---|---|---|---|---|---|---|---|---|---|
| Merismopedia sp. SIO2A8 | 94 | 6E-73 | 42.19 | WP_106290463.1 | | | | | 84 | 3E-34 | 42.18 | WP_106288375.1 |
| Microcoleus sp. IPMA8 | 95 | 2E-48 | 37.75 | NET54256.1 | | | | | 80 | 1E-19 | 35.00 | NET50465.1 |
| Microcoleus sp. IPPAS B-353 | 97 | 3E-76 | 45.04 | WP_172192453.1 | | | | | 88 | 2E-37 | 41.29 | WP_172191685.1 |
| Microcoleus sp. SU_5_3 | 97 | 6E-83 | 47.15 | WP_159786211.1 | 93 | 2E-26 | 27.78 | WP_159789125.1 | 87 | 1E-32 | 38.22 | WP_159789783.1 |
| Microcoleus vaginatus | 93 | 1E-40 | 34.41 | NJK68152.1 | | | | | 89 | 2E-40 | 43.59 | NJK67781.1 |
| Microcystis aeruginosa | 95 | 5E-48 | 38.65 | WP_006634175.1 | | | | | 88 | 2E-36 | 41.29 | WP_006633898.1 |
| Microcystis aeruginosa K13-06 | 95 | 3E-55 | 40.80 | WP_002748972.1 | 92 | 5E-40 | 30.33 | WP_123230411.1 | 83 | 2E-22 | 35.62 | WP_044034309.1 |
| Microcystis aeruginosa Ma_MB_F_20061100_S20 | | | | | | | | | 81 | 3E-22 | 37.32 | NCR76265.1 |
| Microcystis aeruginosa PCC 9808 | | | | | | | | | 81 | 7E-21 | 36.62 | CCI26471.1 |
| Microcystis novacekii Mn_MB_F_20050700_S1 | | | | | 92 | 4E-39 | 30.33 | TRU35748.1 | 81 | 2E-22 | 37.32 | TRU34683.1 |
| Moorea sp. SIO4A1 | 95 | 3E-40 | 34.66 | TRU88955.1 | 92 | 9E-38 | 29.62 | TRU88278.1 | 81 | 1E-22 | 37.32 | TRU80218.1 |
| Myxacorys almedinensis | | | | | | | | | 65 | 4E-24 | 42.86 | NEQ63733.1 |
| Myxosarcina sp. GI1 | 97 | 2E-83 | 47.91 | WP_162425488.1 | 72 | 8E-15 | 23.65 | WP_162424814.1 | 88 | 5E-38 | 40.65 | WP_162421194.1 |
| Nodularia sp. (in Bacteria) | 97 | 4E-85 | 49.43 | WP_036485973.1 | 57 | 2E-09 | 25.63 | WP_052055483.1 | 90 | 1E-32 | 37.71 | WP_036485441.1 |
| Nodularia sp. NIES-3585 | | | | | | | | | 92 | 3E-33 | 38.27 | TVP56687.1 |
| Nodularia sp. NIES-3585 | 97 | 2E-80 | 45.25 | WP_089093690.1 | 56 | 4E-17 | 25.56 | WP_089092999.1 | 92 | 3E-32 | 37.65 | WP_089092975.1 |
| Nodularia spumigena | | | | | 56 | 4E-17 | 25.56 | WP_089092999.1 | 90 | 3E-31 | 37.97 | GAX37826.1 |
| Nodularia spumigena CCY9414 | 97 | 6E-82 | 46.01 | WP_006194082.1 | 72 | 5E-20 | 26.25 | WP_006194451.1 | 92 | 2E-32 | 38.27 | WP_063874063.1 |
| Nostoc azollae' 0708 | 93 | 2E-37 | 33.74 | AHJ27233.1 | 69 | 4E-20 | 26.30 | AHJ31646.1 | 90 | 4E-31 | 38.61 | EAW45645.1 |
| Nostoc calcicola | | | | | | | | | 68 | 9E-28 | 40.00 | ADI65736.1 |
| Nostoc carneum | 96 | 1E-85 | 47.71 | WP_073644990.1 | | | | | 88 | 3E-33 | 38.06 | WP_073645619.1 |
| Nostoc carneum NIES-2107 | 97 | 2E-85 | 48.67 | WP_096729628.1 | 55 | 7E-10 | 25.30 | WP_096731955.1 | 86 | 8E-34 | 40.40 | WP_096731959.1 |
| Nostoc commune | 95 | 7E-38 | 34.14 | BAY34553.1 | 55 | 9E-10 | 25.84 | BAY30768.1 | 88 | 2E-34 | 40.00 | BAY30791.1 |
| Nostoc commune HK-02 | 96 | 3E-85 | 48.85 | WP_109009020.1 | | | | | 88 | 6E-34 | 39.88 | WP_109012677.1 |
| Nostoc cycadae | | | | | | | | | 88 | 1E-33 | 39.88 | BBD68165.1 |
| Nostoc flagelliforme | 96 | 1E-76 | 44.40 | WP_103126066.1 | | | | | 86 | 2E-32 | 38.41 | WP_103124552.1 |

| | | | | | | | | | | | | |
|---|---|---|---|---|---|---|---|---|---|---|---|---|
| Nostoc flagelliforme CCNUN1 | 96 | 8E-84 | 48.09 | WP_100900796.1 | | | | | 88 | 3E-34 | 40.00 | WP_100903263.1 |
| Nostoc linckia | | | | | | | | | 88 | 3E-34 | 40.00 | AUB42704.1 |
| Nostoc linckia z1 | 96 | 6E-85 | 47.51 | WP_099072297.1 | 59 | 2E-21 | 31.10 | WP_143874802.1 | 88 | 2E-33 | 38.06 | WP_096538304.1 |
| Nostoc minutum NIES-26 | | | | | | | | | 86 | 2E-31 | 38.41 | PHJ57043.1 |
| Nostoc piscinale | 97 | 2E-85 | 47.53 | RCJ18721.1 | | | | | 88 | 6E-29 | 39.88 | RCJ22982.1 |
| Nostoc punctiforme | 96 | 1E-75 | 44.02 | WP_062291054.1 | | | | | 86 | 1E-32 | 37.75 | WP_062289723.1 |
| Nostoc sp. 106C | 96 | 2E-82 | 48.47 | RCJ38514.1 | | | | | 88 | 2E-31 | 38.04 | WP_012411601.1 |
| Nostoc sp. 3335mG | 97 | 7E-78 | 45.21 | WP_086764274.1 | | | | | 86 | 5E-36 | 41.72 | WP_086760234.1 |
| Nostoc sp. ATCC 43529 | 93 | 1E-46 | 38.91 | WP_110157102.1 | 88 | 2E-25 | 25.12 | WP_110150531.1 | 92 | 1E-30 | 43.21 | WP_110155645.1 |
| Nostoc sp. ATCC 53789 | 96 | 1E-83 | 45.98 | RCJ24554.1 | | | | | 88 | 2E-32 | 38.06 | RCJ19014.1 |
| Nostoc sp. B(2019) | 96 | 1E-74 | 44.02 | WP_114080702.1 | | | | | 88 | 7E-34 | 40.00 | WP_114083126.1 |
| Nostoc sp. CENA543 | 96 | 1E-83 | 47.71 | WP_162399753.1 | | | | | 92 | 5E-38 | 41.98 | WP_162397461.1 |
| Nostoc sp. HK-01 | 97 | 2E-80 | 46.01 | WP_103138075.1 | | | | | 88 | 2E-32 | 37.43 | WP_103140187.1 |
| Nostoc sp. KVJ20 | 94 | 8E-47 | 37.65 | BBD63483.1 | | | | | 86 | 2E-32 | 38.41 | BBD58761.1 |
| Nostoc sp. 'Lobaria pulmonaria (5183) cyanobiont' | 96 | 9E-86 | 48.85 | WP_069071839.1 | | | | | 88 | 7E-34 | 39.38 | WP_069069359.1 |
| Nostoc sp. MBR 210 | 96 | 3E-75 | 45.17 | WP_104905126.1 | | | | | 88 | 8E-34 | 39.88 | WP_104909130.1 |
| Nostoc sp. NIES-2111 | 97 | 5E-82 | 46.39 | OCQ94408.1 | | | | | 86 | 4E-33 | 37.75 | OCQ90230.1 |
| Nostoc sp. NIES-3756 | 97 | 8E-82 | 50.00 | WP_096681397.1 | | | | | 88 | 2E-34 | 41.29 | WP_096680476.1 |
| Nostoc sp. NIES-4103 | 97 | 3E-82 | 50.38 | WP_067767605.1 | | | | | 88 | 5E-35 | 41.29 | WP_067770169.1 |
| Nostoc sp. NIES-4103 | 96 | 5E-77 | 44.02 | WP_096553559.1 | | | | | 90 | 7E-35 | 44.03 | WP_096553899.1 |
| Nostoc sp. PA-18-2419 | 96 | 5E-77 | 44.02 | WP_096553559.1 | | | | | 88 | 4E-34 | 44.52 | BAZ49694.1 |
| Nostoc sp. PCC 7107 | 96 | 4E-76 | 45.17 | WP_138502447.1 | | | | | 88 | 2E-33 | 39.35 | WP_138498090.1 |
| Nostoc sp. PCC 7120 = FACHB-418 | 95 | 4E-49 | 37.35 | WP_015111850.1 | | | | | 88 | 3E-32 | 37.42 | WP_172641455.1 |
| Nostoc sp. PCC 7524 | 86 | 2E-33 | 35.62 | RUR88738.1 | | | | | 90 | 1E-36 | 40.88 | RUR89562.1 |

| | | | | | | | | | | | | |
|---|---|---|---|---|---|---|---|---|---|---|---|---|
| Nostoc sp. 'Peltigera malacea cyanobiont' DB3992 | 97 | 5E-82 | 46.77 | WP_015136635.1 | | | | | 89 | 2E-38 | 43.31 | WP_015139967.1 |
| Nostoc sp. 'Peltigera membranacea cyanobiont' 210A | 96 | 3E-84 | 47.89 | WP_099102694.1 | | | | | 88 | 4E-34 | 39.88 | WP_099101087.1 |
| Nostoc sp. 'Peltigera membranacea cyanobiont' 232 | 96 | 1E-83 | 47.89 | WP_094351066.1 | | | | | 88 | 4E-35 | 40.65 | WP_094353124.1 |
| Nostoc sp. RF31YmG | 96 | 2E-75 | 44.79 | WP_094341890.1 | | | | | 88 | 4E-31 | 38.04 | WP_094343620.1 |
| Nostoc sp. T09 | 97 | 3E-77 | 44.83 | WP_086837385.1 | | | | | 86 | 7E-37 | 40.40 | WP_086837235.1 |
| Nostoc sp. TCL240-02 | 97 | 5E-86 | 48.67 | WP_086689782.1 | | | | | 88 | 3E-34 | 39.35 | WP_086687115.1 |
| Nostoc sp. UIC 10630 | 96 | 4E-84 | 49.24 | WP_174711313.1 | | | | | 93 | 4E-33 | 38.04 | WP_174713387.1 |
| Nostoc sphaeroides | 96 | 4E-74 | 43.63 | WP_163927577.1 | | | | | 88 | 2E-37 | 41.94 | WP_163928764.1 |
| Nostoc sphaeroides CCNUC1 | 96 | 3E-83 | 47.89 | WP_118166525.1 | | | | | 88 | 4E-31 | 38.65 | WP_118171620.1 |
| Nostocaceae | 75 | 2E-20 | 32.84 | WP_010999482.1 | | | | | 93 | 9E-38 | 40.49 | WP_010997237.1 |
| Nostocales | 73 | 2E-20 | 34.52 | WP_096577165.1 | | | | | 88 | 2E-35 | 40.65 | WP_096575223.1 |
| Nostocales cyanobacterium HT-58-2 | 85 | 9E-57 | 37.93 | QFS50731.1 | | | | | 88 | 4E-31 | 38.65 | QFS46173.1 |
| Okeania hirsuta | 96 | 3E-75 | 44.40 | WP_087538690.1 | | | | | 88 | 2E-37 | 43.23 | WP_087537718.1 |
| Okeania sp. SIO1H6 | 96 | 7E-52 | 38.82 | WP_124154550.1 | 90 | 4E-32 | 27.99 | WP_124155155.1 | 84 | 7E-41 | 45.95 | WP_124143786.1 |
| Okeania sp. SIO2B3 | 95 | 2E-46 | 37.70 | NET17153.1 | | | | | 55 | 4E-21 | 43.30 | NET17604.1 |
| Okeania sp. SIO2C2 | 95 | 1E-47 | 38.49 | NET45732.1 | 90 | 1E-30 | 27.75 | NET41378.1 | 82 | 2E-41 | 47.22 | NET46871.1 |
| Okeania sp. SIO2C9 | 96 | 8E-49 | 39.22 | NEP90059.1 | 90 | 9E-32 | 27.27 | NEP91011.1 | 84 | 2E-40 | 45.95 | NEP89828.1 |
| Okeania sp. SIO2D1 | 96 | 4E-52 | 39.61 | NEQ72396.1 | 58 | 3E-29 | 30.74 | NEQ73014.1 | 84 | 3E-41 | 45.95 | NEQ73389.1 |
| Okeania sp. SIO2F4 | 95 | 8E-47 | 38.19 | NES65406.1 | 90 | 3E-32 | 27.99 | NES63798.1 | 84 | 7E-41 | 46.62 | NES66674.1 |
| Okeania sp. SIO3B3 | 97 | 9E-86 | 49.24 | NES03833.1 | 90 | 7E-30 | 27.27 | NES02411.1 | 89 | 3E-39 | 42.68 | NES03309.1 |
| Okeania sp. SIO3B5 | 96 | 1E-84 | 50.00 | NEP84905.1 | 90 | 1E-31 | 26.79 | NEP81791.1 | 84 | 8E-40 | 45.27 | NEP84377.1 |
| Okeania sp. SIO3H1 | 92 | 4E-32 | 31.62 | NEO53097.1 | | | | | 84 | 2E-40 | 45.27 | NEO56124.1 |
| Okeania sp. SIO3I5 | 95 | 3E-45 | 36.00 | NEN92795.1 | 90 | 2E-31 | 27.03 | NEN88300.1 | 84 | 2E-41 | 45.95 | NEN92322.1 |
| Okeania sp. SIO4D6 | 95 | 7E-48 | 39.92 | NEQ40992.1 | 92 | 3E-36 | 29.04 | NEQ40766.1 | 88 | 1E-40 | 44.52 | NEQ34970.1 |
| Oscillatoria acuminata | 96 | 3E-51 | 39.61 | NEP04696.1 | 90 | 1E-32 | 27.27 | NEP05998.1 | 84 | 2E-41 | 45.95 | NEP05027.1 |

| | | | | | | | | | | | | |
|---|---|---|---|---|---|---|---|---|---|---|---|---|
| Oscillatoria acuminata PCC 6304 | 99 | 1E-84 | 49.62 | WP_015149460.1 | 68 | 1E-17 | 26.33 | WP_044195236.1 | 90 | 2E-34 | 38.36 | WP_044198228.1 |
| Oscillatoria nigro-viridis | | | | | 68 | 1E-17 | 26.33 | AFY82766.1 | 93 | 1E-35 | 38.04 | AFY85409.1 |
| Oscillatoria sp. PCC 10802 | 97 | 3E-76 | 44.66 | WP_015175150.1 | | | | | 88 | 8E-34 | 42.58 | WP_015177042.1 |
| Oscillatoria sp. SIO1A7 | 97 | 2E-77 | 44.66 | WP_017717521.1 | | | | | 94 | 7E-37 | 40.00 | WP_044170098.1 |
| Oscillatoriales cyanobacterium | 98 | 1E-79 | 46.97 | NER35665.1 | | | | | 89 | 8E-38 | 39.74 | NER35059.1 |
| Oscillatoriales cyanobacterium SM2_3_0 | 97 | 2E-87 | 49.81 | HFM98699.1 | | | | | 88 | 1E-39 | 45.16 | TAE82010.1 |
| Oscillatoriales cyanobacterium SpSt-418 | 78 | 1E-19 | 30.88 | HFM98640.1 | | | | | 90 | 5E-33 | 36.48 | HFM98249.1 |
| Phormidesmis priestleyi | 96 | 5E-77 | 46.12 | NJK37169.1 | 88 | 3E-28 | 27.60 | NJK40038.1 | 77 | 7E-31 | 39.26 | NJK38876.1 |
| Phormidium sp. HE10JO | 97 | 2E-80 | 46.77 | WP_073072482.1 | 89 | 2E-26 | 26.76 | PZO48325.1 | 90 | 3E-33 | 38.99 | WP_068817132.1 |
| Phormidium sp. SL48-SHIP | 95 | 1E-48 | 38.40 | WP_087708474.1 | 90 | 4E-30 | 29.50 | WP_087711903.1 | 75 | 2E-38 | 48.15 | WP_087708848.1 |
| Phormidium willei | 97 | 3E-84 | 49.04 | TAN99368.1 | 90 | 6E-28 | 28.71 | TAN96160.1 | 74 | 1E-38 | 49.25 | TAO01427.1 |
| Planktothricoides sp. SpSt-374 | 97 | 4E-77 | 45.80 | HGF99763.1 | | | | | 88 | 1E-43 | 43.87 | HGG00689.1 |
| Planktothrix serta | 95 | 2E-49 | 38.80 | WP_068789296.1 | 90 | 5E-29 | 29.26 | WP_150121656.1 | 75 | 2E-37 | 47.41 | WP_150121654.1 |
| Planktothrix tepida | 95 | 4E-49 | 39.36 | WP_083623841.1 | | | | | 88 | 1E-31 | 41.29 | WP_083625784.1 |
| Planktothrix tepida PCC 9214 | 93 | 4E-37 | 34.01 | CUR30420.1 | | | | | 88 | 1E-29 | 40.65 | CUR30755.1 |
| Pleurocapsa minor | 95 | 9E-49 | 39.46 | WP_072722045.1 | | | | | 85 | 5E-29 | 41.06 | WP_072718661.1 |
| Pleurocapsa sp. CCALA 161 | 85 | 7E-74 | 48.71 | WP_083888297.1 | | | | | 92 | 4E-35 | 38.27 | WP_041392236.1 |
| Pleurocapsa sp. PCC 7319 | 97 | 6E-74 | 46.01 | WP_106238382.1 | 90 | 1E-29 | 28.57 | WP_106234929.1 | 90 | 7E-33 | 39.24 | WP_106242292.1 |
| Pleurocapsa sp. SU_5_0 | 97 | 4E-87 | 49.43 | WP_019503740.1 | 90 | 4E-37 | 29.52 | WP_019509109.1 | 90 | 6E-40 | 40.51 | WP_019507816.1 |
| Pseudanabaena sp. | 97 | 3E-81 | 46.77 | NJK58627.1 | 90 | 2E-28 | 27.62 | NJK56951.1 | 92 | 1E-33 | 37.89 | NJK56026.1 |
| Richelia sp. SM2_1_7 | 97 | 2E-82 | 47.31 | PZV17439.1 | 90 | 6E-29 | 27.34 | PZU95321.1 | 85 | 4E-26 | 36.67 | PZV17064.1 |
| Rippkaea orientalis | 70 | 2E-32 | 37.70 | NJL79557.1 | 59 | 3E-28 | 34.17 | NJL80862.1 | 86 | 2E-39 | 44.37 | NJL79892.1 |
| Rivularia sp. PCC 7116 | 94 | 7E-50 | 37.65 | WP_015785241.1 | 92 | 2E-26 | 26.68 | WP_015784206.1 | 90 | 7E-40 | 38.99 | WP_012595850.1 |
| Scytonema hofmanni UTEX B 1581 | 97 | 2E-82 | 48.09 | WP_015120391.1 | 90 | 2E-38 | 30.31 | WP_015122239. | 93 | 2E-41 | 43.56 | WP_015121982.1 |
| Scytonema hofmannii | | | | | | | | | 90 | 9E-35 | 39.62 | WP_029631955.1 |

| | | | | | | | | | | | | |
|---|---|---|---|---|---|---|---|---|---|---|---|---|
| Scytonema hofmannii PCC 7110 | 97 | 7E-85 | 49.24 | WP_017747811.1 | | | | | 85 | 1E-34 | 40.00 | WP_017739839.1 |
| Scytonema sp. HK-05 | | | | | | | | | 83 | 1E-33 | 40.41 | KYC40844.1 |
| Scytonema sp. NIES-4073 | 94 | 2E-76 | 44.72 | WP_073634910.1 | | | | | 88 | 3E-38 | 44.52 | WP_073627560.1 |
| Scytonema sp. RU_4_4 | 97 | 3E-85 | 48.09 | WP_096562020.1 | | | | | 81 | 2E-36 | 46.15 | WP_096569964.1 |
| Scytonema sp. UIC 10036 | 95 | 2E-46 | 37.55 | NJM71673.1 | | | | | 88 | 4E-37 | 44.87 | NJM69287.1 |
| Scytonema tolypothrichoides | 94 | 3E-75 | 43.82 | WP_155751066.1 | 93 | 9E-35 | 27.04 | WP_155742999.1 | 88 | 2E-35 | 40.65 | WP_155747426.1 |
| Snowella sp. | 94 | 3E-75 | 43.82 | WP_152368908.1 | | | | | 88 | 2E-38 | 45.81 | WP_048867059.1 |
| Spirulina sp. SIO3F2 | 94 | 1E-70 | 41.60 | NEO87985.1 | 89 | 2E-30 | 26.97 | NEO85737.1 | 90 | 1E-44 | 47.17 | NEO86388.1 |
| Stanieria cyanosphaera | 94 | 7E-49 | 38.46 | PZV27181.1 | | | | | 96 | 2E-24 | 32.37 | PZV24310.1 |
| Stanieria sp. NIES-3757 | 97 | 3E-88 | 50.19 | WP_015194130.1 | | | | | 89 | 9E-34 | 37.97 | WP_015191460.1 |
| Symploca sp. SIO2B6 | 97 | 3E-86 | 49.81 | WP_096383972.1 | | | | | 91 | 2E-34 | 36.65 | WP_096382719.1 |
| Symploca sp. SIO2D2 | 94 | 2E-71 | 41.70 | NET09383.1 | | | | | 80 | 1E-20 | 36.88 | NET08794.1 |
| Symploca sp. SIO3C6 | 94 | 3E-51 | 39.68 | NEQ70357.1 | | | | | 77 | 2E-33 | 42.65 | NEQ68681.1 |
| Synechococcaceae cyanobacterium SM2_3_2 | 95 | 2E-45 | 36.80 | NEO30818.1 | | | | | 88 | 6E-40 | 43.87 | NEO28981.1 |
| Synechococcales cyanobacterium C | 97 | 7E-75 | 46.77 | NJL97752.1 | 91 | 1E-25 | 27.52 | NJL98955.1 | 85 | 8E-37 | 45.03 | NJL97528.1 |
| Synechococcus sp. PCC 7336 | 97 | 6E-85 | 46.39 | WP_161827197.1 | | | | | 77 | 2E-27 | 39.26 | WP_161826694.1 |
| Synechocystis sp. PCC 7509 | 99 | 1E-78 | 48.12 | WP_017325474.1 | 89 | 1E-28 | 26.83 | WP_017325515.1 | 91 | 7E-34 | 38.12 | WP_017326127.1 |
| Thermoleptolyngbya sp. PKUAC-SCTA183 | 94 | 2E-74 | 42.68 | WP_009632366.1 | | | | | 85 | 4E-35 | 42.00 | WP_009633957.1 |
| Tolypothrix bouteillei | 96 | 4E-81 | 46.74 | WP_172353582.1 | 64 | 2E-23 | 30.23 | WP_172358739.1 | 88 | 1E-20 | 35.44 | WP_172359099.1 |
| Tolypothrix campylonemoides | 94 | 7E-74 | 43.03 | WP_038081815.1 | | | | | 88 | 4E-35 | 41.29 | WP_038073395.1 |
| Tolypothrix sp. NIES-4075 | 94 | 6E-75 | 43.50 | WP_041037159.1 | | | | | 88 | 1E-38 | 45.16 | WP_041037423.1 |
| Tolypothrix sp. PCC 7601 | 85 | 2E-19 | 27.95 | WP_089129014.1 | | | | | 92 | 1E-38 | 42.59 | WP_089129715.1 |
| Tolypothrix sp. PCC 7910 | 70 | 1E-19 | 33.86 | EKF03756.1 | | | | | 88 | 9E-33 | 38.71 | EKF02310.1 |
| Trichodesmium erythraeum | 97 | 2E-86 | 49.43 | WP_167726011.1 | | | | | 86 | 7E-34 | 39.74 | WP_167727755.1 |

| | | | | | | | | | | | | |
|---|---|---|---|---|---|---|---|---|---|---|---|---|
| Trichormus variabilis | 92 | 2E-19 | 27.13 | WP_011610524.1 | 90 | 4E-30 | 28.10 | WP_011613768.1 | 83 | 1E-38 | 43.84 | WP_011612277.1 |
| Trichormus variabilis ATCC 29413 | 94 | 2E-25 | 29.92 | WP_127056832.1 | 69 | 2E-19 | 25.55 | WP_127053830.1 | 93 | 4E-39 | 44.17 | WP_127055370.1 |
| unclassified Calothrix | 94 | 1E-36 | 31.87 | WP_096687949.1 | | | | | 86 | 3E-32 | 41.72 | WP_096694531.1 |
| unclassified Nostoc | 96 | 2E-34 | 31.10 | WP_086756103.1 | | | | | 88 | 1E-31 | 38.04 | WP_094328614.1 |
| unicellular cyanobacterium SU2 | 95 | 3E-26 | 30.04 | ABA23926.1 | | | | | 90 | 3E-36 | 42.14 | ABA24455.1 |
| unicellular cyanobacterium SU3 | 93 | 5E-48 | 36.59 | WP_085434531.1 | 92 | 6E-30 | 29.51 | WP_085435375.1 | 86 | 4E-45 | 45.70 | WP_085436845.1 |

Table S2: List of *Synechocystis* strains used in this study

| strain | marker of genotype | reference |
|---|---|---|
| Δ*eda* | *sll0107*::gm$^{R}$ | Chen et al 2016 |
| Δ*gnd** (see line below) | *sll0329*::em$^{R}$ | Chen et al 2016 |
| *Δ*gnd* was identified as Δ*gnd zwf*399+A in this study | *sll0329*::em$^{R}$ | This study |
| Δ*zwf* | *slr1843*::cm$^{R}$ | Chen et al 2016 |
| Δ*eda*Δ*gnd* | *sll0107*::gm$^{R}$, *sll0329*::em$^{R}$ | Doello et al 2018 |
| Δ*eda*Δ*gnd*Δ*zwf* | *sll0107*::gm$^{R}$, *sll0329*::em$^{R}$, *slr1843*::cm$^{R}$ | This study (plasmid for deletion of zwf: Chen et al. 2026) |
| Δ*hk* | *sll0539*::sp$^{R}$ | Theune et al. 2020 |
| Δ*hk*::*hk* | *sll0539*::sp$^{R}$ cm | This study |
| Sll1709:oe | *sll1709*::km$^{R}$ | This study |

Table S3: List of used primers for *Synechocystis* used in this study

| mutant | primer | sequence |
|---|---|---|
| Complementation Δ*hk*:*hk* | Hk-out1 | CTATAGGGCGAATTGGGTACCATTGGCCAAAAATAGCGCAAAGGTT |
| | Cm-Hk | ATAATATCGAATTCCTGCAGCTAAGACTGGGCCGCACAAACC |
| | Hk-Cm | TTTGTGCGGCCCAGTCTTAGCTGCAGGAATTCGATATTATTGAA |
| | Cm-hkin2 | GAGGTGCCGCCATCAAGCTTGAAGTGGACATTTTTAGTTGTTAACT |
| | Hkin2-Cm | CAACTAAAAATGTCCACTTCAAGCTTGATGGCGGCACCTCGCTAA |
| | Hkout2 | AGGGAACAAAAGCTGGAGCTACCCCTGGTCTTTGGGCGGAACAA |
| pET15b *Saci* 0025 | Forward | GTCCATATGTGGAAATAATTTCACCTATCATTAC |
| | Reverse | GTCGGATCCATGTACCAGTTCTTGAATCTTTCTC |
| EDAK147G | Forward | GTGGCGCAACCGCTGTTGGCGTGTTTCCGATTAAGACC |
| | Reverse | GGTCTTAATCGGAAACACGCCAACAGCGGTTGCGCCAC |

The nucleotide substitutions introducing the EDAK147G exchange are underlined.
